# Brain-wide representations of prior information in mouse decision-making

**DOI:** 10.1101/2023.07.04.547684

**Authors:** Charles Findling, Felix Hubert, International Brain Laboratory, Luigi Acerbi, Brandon Benson, Julius Benson, Daniel Birman, Niccolò Bonacchi, Sebastian Bruijns, Matteo Carandini, Joana A Catarino, Gaelle A Chapuis, Anne K Churchland, Yang Dan, Felicia Davatolhagh, Eric EJ DeWitt, Tatiana A Engel, Michele Fabbri, Mayo Faulkner, Ila Rani Fiete, Laura Freitas-Silva, Berk Gerçek, Kenneth D Harris, Michael Häusser, Sonja B Hofer, Fei Hu, Julia M Huntenburg, Anup Khanal, Chris Krasniak, Christopher Langdon, Peter E Latham, Petrina Y P Lau, Zach Mainen, Guido T Meijer, Nathaniel J Miska, Thomas D Mrsic-Flogel, Jean-Paul Noel, Kai Nylund, Alejandro Pan-Vazquez, Liam Paninski, Jonathan Pillow, Cyrille Rossant, Noam Roth, Rylan Schaeffer, Michael Schartner, Yanliang Shi, Karolina Z Socha, Nicholas A Steinmetz, Karel Svoboda, Charline Tessereau, Anne E Urai, Miles J Wells, Steven Jon West, Matthew R Whiteway, Olivier Winter, Ilana B Witten, Anthony Zador, Yizi Zhang, Peter Dayan, Alexandre Pouget

**Affiliations:** University of Geneva, Switzerland; University of Helsinki; Stanford University, USA; New York University, USA; University of Washington, USA; Champalimaud Foundation, Portugal; University College London, UK; University of California Los Angeles, USA; University of California Berkeley, USA; Princeton University, USA; USA; Massachusetts Institute of Technology, USA; Sainsbury Wellcome Center, University College London, UK; Max Planck Institute, University of Tübingen, Germany; Cold Spring Harbor Laboratory; Gatsby Computational Neuroscience Unit, UK; Columbia University, USA; Allen Institute for Neural Dynamics, USA; Leiden University, The Netherlands

## Abstract

The neural representations of prior information about the state of the world are poorly understood. To investigate them, we examined brain-wide Neuropixels recordings and widefield calcium imaging collected by the International Brain Laboratory. Mice were trained to indicate the location of a visual grating stimulus, which appeared on the left or right with prior probability alternating between 0.2 and 0.8 in blocks of variable length. We found that mice estimate this prior probability and thereby improve their decision accuracy. Furthermore, we report that this subjective prior is encoded in at least 20% to 30% of brain regions which, remarkably, span all levels of processing, from early sensory areas (LGd, VISp) to motor regions (MOs, MOp, GRN) and high level cortical regions (ACAd, ORBvl). This widespread representation of the prior is consistent with a neural model of Bayesian inference involving loops between areas, as opposed to a model in which the prior is incorporated only in decision-making areas. This study offers the first brain-wide perspective on prior encoding at cellular resolution, underscoring the importance of using large scale recordings on a single standardized task.

---

The ability to combine sensory information with prior knowledge through probabilistic inference is crucial for perception and cognition. In simple cases, inference is performed near-optimally by the brain, following key precepts of Bayesian decision theory (Ernst & Banks, 2002; Jacobs, 1999; Knill & Pouget, 2004; Mamassian et al., 1998; Weiss et al., 2002). For example, when interpreting a visual scene, we assume *a priori* that light comes from above – a sensible assumption which allows us to interpret otherwise ambiguous images (Mamassian et al., 1998).

While much theoretical work has been devoted to the neural representation of Bayesian inference (Echeveste et al., 2020; Ganguli & Simoncelli, 2014; Ma et al., 2006; Soltani & Wang, 2010), it remains unclear where and how prior knowledge is represented in the brain. At one extreme, the brain might combine prior information with sensory evidence in high level decision-making brain regions, right before decisions are turned into actions. This would predict that prior information is encoded only in late stages of processing, as has indeed been reported in parietal, orbitofrontal and prefrontal cortical areas (Forstmann, 2010; Hanks et al., 2011; Hansen et al., 2012; Mulder et al., 2012; Niv, 2019; Nogueira et al., 2017; Rao et al., 2012). At the other extreme, the brain might operate like a very large Bayesian network, in which probabilistic inference is the *modus operandi* in all brain regions and inference can be performed in all directions (Ackley et al., 1985; Berkes et al., 2011; Bondy et al., 2018; Jardri et al., 2017; Kok et al., 2012; Lange & Haefner, 2022). This would allow neural circuits to infer beliefs over variables from observations of arbitrary combinations of other variables. For instance, upon seeing an object, the brain might be able to infer its auditory and tactile properties; but could just as well perform the reverse inference, i.e., predicting its visual appearance upon hearing or touching it. Such a model would predict that prior information should be available throughout the brain, even in low level cortical sensory areas (Berkes et al., 2011; Bondy et al., 2018; Kok et al., 2012; Lange & Haefner, 2022). The current literature offers a contradictory, and thus inconclusive, perspective on whether the prior is indeed encoded in brain regions associated with early processing (Bell et al., 2016; Bondy et al., 2018; Haefner et al., 2016; Han & Helmchen, 2023; Hanks et al., 2011; Ishizu et al., 2023; Mayrhofer et al., 2019; Park et al., 2022; Platt & Glimcher, 1999; Rao et al., 2012; Son et al., 2023). This is because past studies have collectively recorded from only a limited set of areas; and, since they use different tasks, even these results cannot be fully integrated.

To address this problem, we analyzed brain-wide data from the International Brain Laboratory, which provides electrophysiological recordings from 242 brain regions, defined by the Allen Common Coordinate Framework (Wang et al., 2020), as well as from widefield imaging (WFI) data from layers 2/3 of cortex of mice performing the same decision-making task (International Brain Lab et al., 2023; International Brain Laboratory et al., 2021). Our results suggest that the prior is encoded cortically and subcortically, across all levels of the brain, including early sensory regions.

## Mice use the prior to optimize their performance

Mice were trained to discriminate whether a visual stimulus, of different contrasts, appeared in the right or left visual field (Fig. 1a). Importantly, the prior probability that the stimulus appeared on the right side switched in a random and uncued manner between 0.2 and 0.8 in blocks of 20-100 trials (Fig. 1b). Knowledge of the current prior would help the mice perform well; in particular, the prior is the only source of information on zero contrast trials, as the probability of reward on these trials is determined by the block probability. We refer to the experimentally-determined prior as the ‘true block prior’. Since the presence of the blocks was not explicitly cued, mice could only form a subjective estimate of the true block prior from the trial history. At best, they could compute the estimate of the true block prior given full knowledge of the task structure and the sequence of previous stimulus sides since the start of a session. We refer to this hereafter as the Bayes-optimal prior (see Methods, Fig. 1b and 2a).

**Figure 1:**
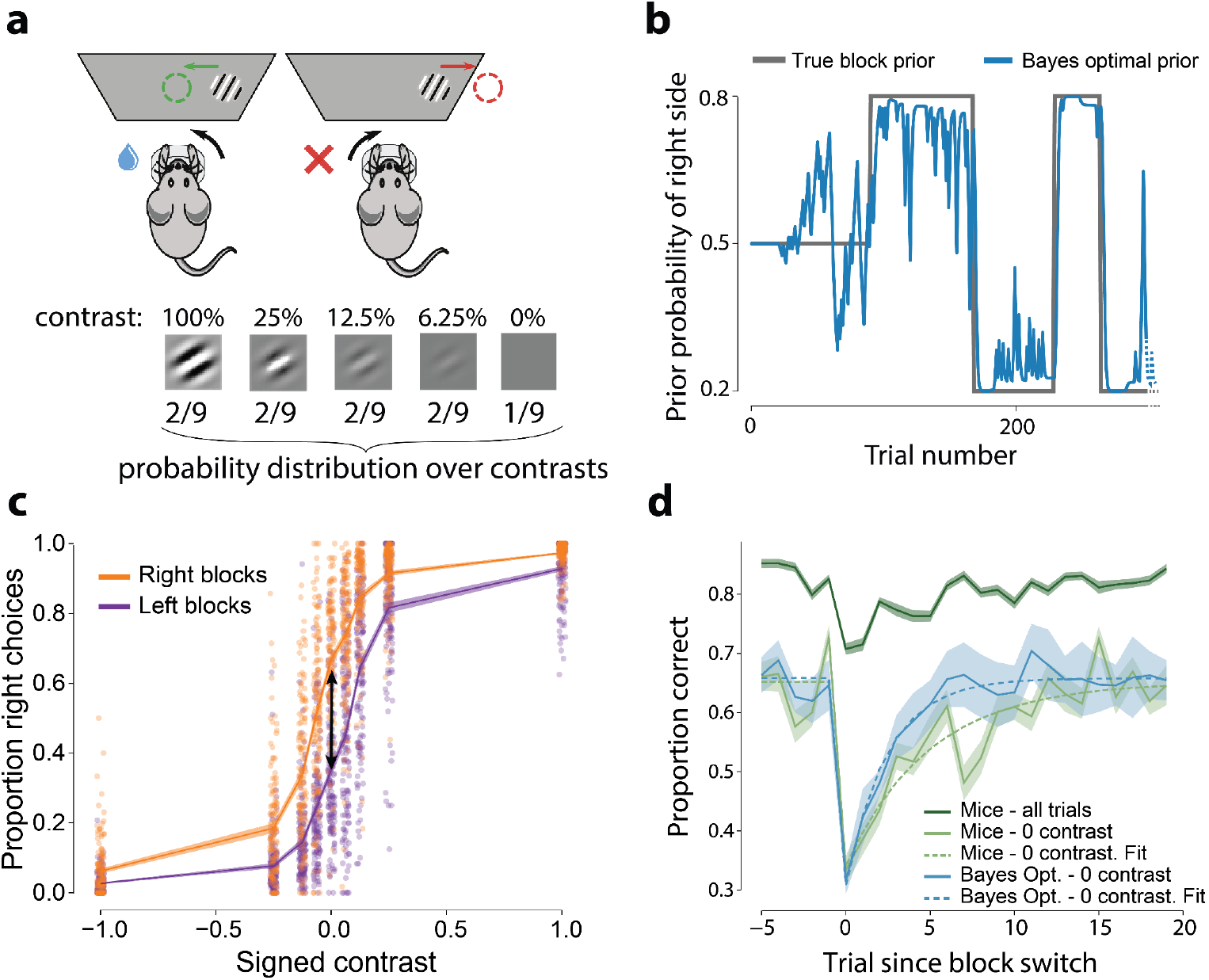
Mice use the block prior to improve their performance. **a**. Mice had to move a visual grating, appearing 35° in the periphery (here shown on the right hand side), to the center of the screen by tuning a wheel with their front paws. The contrast of the visual stimulus varied from trial to trial. **b**. The prior probability that the stimulus appeared on the right side was maintained at either 0.2 or 0.8 over blocks of trials, after an initial block of 90 trials during which the prior was set to 0.5. The length of a block was drawn from a truncated exponential distribution between 20 and 100 trials, with the scale parameter of the exponential set to 60 trials. Following a wheel turn, the mouse was provided with positive feedback in the form of a water reward, or negative feedback in the form of a white noise pulse and timeout. The next trial began after a delay, followed by a quiescence period uniformly sampled between 400ms and 700ms during which the mice had to hold the wheel still. **c**. Psychometric curves averaged across all 139 animals and 459 sessions and conditioned on block identity. The proportion of right choices on zero contrast trials was significantly different across blocks (2-tailed signed-rank Wilcoxon paired test: *t*=15, *p*=2.04e-24, N=139) and displaced in the direction predicted by the true block prior (black double arrow). **d**. Reversal curves showing the percentage of correct responses following block switches. Dark green: average performance across all animals and all contrasts. Light green: same as dark green but for zero contrast trials only. Blue: performance of an observer generating choices stochastically according to the Bayes-optimal estimate of the prior on zero contrast trials. This simulation was limited to zero-contrast trials to focus on the influence of prior knowledge without stimulus information. Shaded region around the mean shows the s.e.m across mice for the curves showing mouse behavior (light and dark green curves) and the standard deviation for the Bayes-optimal model (blue curve), as there is no inter-individual variability to account for.

Analyzing choice behavior revealed that mice leveraged the block structure to improve their performance. Psychometric curves conditioned on right and left blocks, averaged across all animals and all sessions, were displaced relative to each other, in a direction consistent with the true block prior (2-tailed signed-rank Wilcoxon paired test between proportion of right choices on zero contrast trials: *t*=15, *p*=2.0E-24, N=139 mice; Fig. 1c). As a result, animals performed at 58.7% ± 0.4% (mean ± sem) correct for zero contrast trials. This is statistically significantly better than chance (2-tailed signed-rank Wilcoxon *t=89, p*=1.5E-23, N=139 mice), albeit significantly worse than an observer that generates actions by sampling from the Bayes-optimal prior, which would perform at 61.1% ± 1.8% (mean ± std; 2-tailed signed-rank Wilcoxon paired test *t=*2171, *p=*1.5e-8, N=139 mice).

Tracking performance around block switches provided further evidence that the animals estimated and used the prior. Indeed, around block switches performance dropped, presumably because of the mismatch between the subjective and true block prior. Thereafter, performance on zero contrast trials recovered with a decay constant of 4.97 trials (jackknife median, see Methods). This is slower than the observer that generates actions by sampling the Bayes-optimal prior (jackknife median: 2.43, 2-tailed paired t-test *t*=3.35, *p*=0.001, N=139 jackknife replicates; Fig. S1).

### Decoding the prior during the inter-trial interval

In order to determine where the prior is encoded in the mouse’s brain, we used linear regression to decode the Bayes-optimal prior from neural activity during the inter-trial interval (ITI) when wheel movements are minimized (from −600ms to −100ms before stimulus onset, see Methods)(The International Brain Laboratory et al., 2021). Note that decoding the Bayes-optimal prior is more sensible than decoding the true block prior, since mice are not explicitly cued about block identity and, therefore cannot possibly know this latter quantity. We assess the quality of the decoding with an *R*^2^ measure. However, to assess the statistical significance of this value we cannot use standard linear regression methods, since these assume independence of trials, while both neural activity (for instance from slow drift in the recordings stemming from movements of the probes across trials) and the prior exhibit temporal correlations. Instead, we use a “pseudosession” method (Harris, 2020): we first construct a null distribution by decoding the (counterfactual) Bayes-optimal priors computed from stimulus sequences generated by sampling from the same process as that used to generate the stimulus sequence that was actually shown to the mouse (see Methods). A session was deemed to encode the prior significantly if *R*^2^ computed for the actual stimuli is larger than the 95^th^ percentile of the null distribution generated from pseudosessions; effect sizes are reported as a corrected *R*^2^, the difference between the actual *R*^2^ and the median *R*^2^ of the null distribution. All values of *R*^2^ reported in this paper are corrected *R*^2^ unless specified otherwise.

For completeness, we decoded the Bayes-optimal prior, 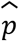, its log odds ratio 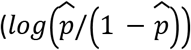 — to test whether neural activity is linearly related to log probabilities as assumed by the theory of probabilistic population codes (Ma et al., 2006) —, the true block prior (fig 1b), and the Bayes-optimal prior on a narrower decoding time window (from −400 to −100ms). For the Bayes-optimal prior, the analysis of the electrophysiological data (Ephys) revealed that around 30.2% of brain areas (73/242 regions), spanning forebrain, midbrain, and hindbrain, encoded the prior significantly (p<.05, pseudosession test; Fisher’s method to combine p-values from multiple recordings of one region, no multiple comparisons correction; Fig. 2b and Fig. S3 for sagittal slices). For example, we could decode the Bayes-optimal prior from a population of *160* neurons in ventrolateral orbitofrontal cortex (ORBvl) with accuracy of *R*^2^ = 0.28 (Fig. 2a, p-value=0.0001). Regions with significant prior encoding include associative cortical areas like the ORBvl and the dorsal anterior cingulate area (ACAd), as well as early sensory areas such as the primary visual cortex (VISp) or the lateral geniculate nucleus (LGd). The Bayes-optimal prior could also be decoded from cortical and subcortical motor areas, such as primary and secondary motor cortex, the intermediate layer of the superior colliculus (SCm), the gigantocellular reticular nucleus (GRN) and the pontine reticular nucleus (PRNr), even though we decoded activity during the inter-trial interval, when wheel movements were minimal (Fig. S2). The encoding of the Bayes-optimal prior is also visible in the PSTH of single neurons (Fig. S4). Decoding the log odds ratio of the Bayes-optimal prior, as opposed to the linear version, revealed consistent findings, with 38.0% (92/242 regions) of regions encoding it significantly across all brain processing levels (Fig. S6). When decoded from a narrower time window (−400ms to −100ms), the Bayes-optimal prior was still significantly decoded across all brain processing levels, albeit with reduced overall decodability (25.6% of regions, 62/242 regions, Fig. S6). An even smaller percentage of regions (19.4%; 47/242 regions, Fig. S6) were found to encode the prior significantly when decoding the true block prior suggesting that the animal’s subjective prior aligns more closely with the Bayes-optimal prior than with the true block prior. This observation is supported by a behavioral analysis, which revealed that a model using the true block as a prior was less effective at explaining behavior compared to the Bayes-optimal model (Fig. S6d). An analysis to determine the necessary number of recordings per region indicated that around 10 recordings per region are required to reach the obtained significance levels (see Fig. S5e). Given that the median number of sessions per region in Ephys is 6 (see Fig. S5d), it is likely that the reported levels of significance are underestimated.

**Figure 2:**
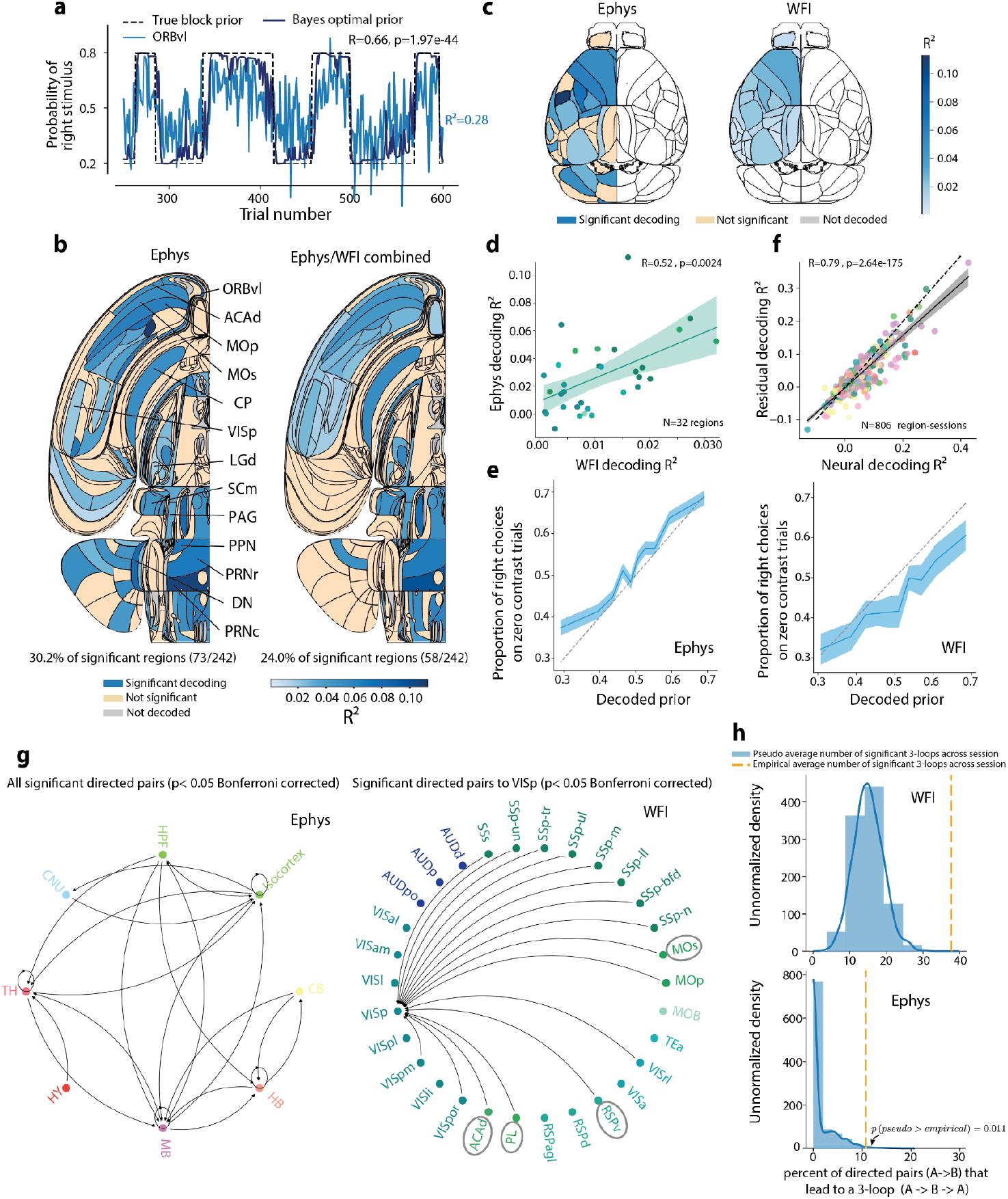
Encoding of the prior across the brain during the inter-trial interval. **a**. Bayes-optimal prior (dark blue) versus Bayes-optimal prior decoded from the ventrolateral orbitofrontal (ORBvl, light blue) on one specific session (corrected *R*^2^=0.28, uncorrected *R*^2^=0.35). Dashed black line: true block prior. **b**. Swanson maps of cross-validated corrected *R*^2^ for areas that have been deemed significant (using Fisher’s method for combining *p*-values, see Methods) Left: Swanson map of *R*^2^ for Ephys data. Right: *R*^2^ across Ephys and WFI. For the left map, a region is deemed significant if the Fisher combined *p*-value is lower than 0.05.

The analysis of widefield calcium imaging data (WFI) suggests an even more widespread encoding of the prior in cortical regions. Indeed, the Bayes-optimal prior was found to be significantly reflected in all dorsal cortical regions (Fig. 2c). This result may reflect a better signal-to-noise ratio, but it might also be due to the calcium signal from axons arising outside these specific areas. However, we also found that the corrected region-specific *R*^2^ for WFI and Ephys modalities were significantly correlated (Spearman correlation R=0.52, p=0.0024, N=32 regions - Fig. 2d). Interpreting the effect size in both Ephys and widefield modalities is challenging due to confounding factors such as the number of sessions and units in Ephys, and the number of pixels in widefield (Fig. S7c,d). To control for correlations between these confounds across modalities (Fig S7e), we corrected the widefield effect size for region sizes. Despite this correction, the correlation between effect sizes across modalities remained significant, and even strengthened (Fig. S7f) - thus suggesting that the effect sizes we decode are, at least partly, specific to the decoded regions.

A quarter of the regions (24.0%, 58/242), still at all levels of brain processing, were found to be significant when merging this larger Ephys data set and WFI data into a single map (using Fisher’s method to combine p-values across Ephys and WFI, see Methods) and applying the Benjamini-Hochberg correction for multiple comparisons with a conservative false discovery rate of 1%, Fig. 2b and Fig. S3 for sagittal slices.

If the decoded prior is truly related to the subjective prior inferred and used by the animal, the amplitude of the decoded prior should be correlated with the animals’ performance on zero contrast trials. Fig. 2e shows that this is indeed the case for both the Ephys and WFI data: on zero contrast trials, the probability that the mice chose the right side was proportional to the cross-validated decoded Bayes-optimal prior of the stimulus appearing on the right (see Methods). Importantly, this relationship remained significant even after controlling for possible drift in the recordings (Fig. S8). Further analysis at the regional level (Fig. S9a) shows a significant relationship in 17.8% of Ephys regions and 90.1% of Widefield regions across all hierarchical levels (LGd, SCm, CP, MOs, ACAd). Additionally, regions that more strongly reflect the prior were more predictive of the animal’s decisions, suggesting the behavioral relevance of the decoded prior (Fig. S9b).

Our results indicate that the Bayes-optimal prior was encoded in multiple areas throughout the brain. However, it is conceivable that mice adjusted their body posture or movement according to the subjective prior and that neural activity in some areas simply reflected these body adjustments. We call this an embodied prior. To test for this possibility, we analyzed video recordings, using Deep Lab Cut (DLC)(Laboratory, International Brain, 2022; Mathis et al., 2018) to estimate the position of multiple body parts, whisking motion energy, and licking during the inter-trial interval (see Methods). We then trained a decoder of the Bayes-optimal prior from these features, and found significant decoding in 38.0% (65/171) of sessions. For these sessions, we found that the *R*^2^ for the prior decoded from video features was correlated with the *R*^2^ for the prior decoded from neural activity (at the brain region level), thus suggesting that the prior signal might be an embodied prior related to body posture (Pearson correlation *R=0*.*18, p=1*.*6E-7*, N=806 region sessions, Supp Fig. S10a). To test for this possibility further, we decoded the prior residual, defined as the Bayes-optimal prior minus the Bayes-optimal prior estimated from video features, from neural activity (again, at the brain region level). If the neural prior simply reflects the embodiment of features extracted by DLC, we should not be able to decode the prior residual from the neural activity and the *R*^2^ of the prior residual should not be correlated with the *R*^2^ of the full prior decoded from neural activity. Crucially, this is not what we observed. Instead, these two values of *R*^2^ are strongly correlated (Fig. 2f, Pearson correlation *R=0*.*89, p=8E-279*, N=806 region sessions) thus suggesting that the neural prior is not an embodied prior, or at least that it cannot be fully explained by the motor features extracted from the video.

To enhance the robustness of our analysis further, we repeated the embodiment study, this time also including eye position data (on sessions on which these were available). This additional step demonstrated that the neural prior could not be entirely attributed to a combination of both motor features and eye position (see Fig. S10b and Fig. S10c for the feature importance). We also specifically checked whether changes in eye position across blocks could account for the significant results in early visual areas such as VISp or LGd. It is indeed conceivable that mice look in the direction of the expected stimulus prior to a trial. If so, what we interpret as a prior signal in these early sensory areas might simply be due to a signal related to eye position. Consistent with this possibility, we found a significant correlation (Pearson correlation *R=0*.*36, p=0*.*0163, N=44* region sessions) between the neural decoding *R*^2^ and the eye position decoding *R*^2^, *i*.*e*, the *R*^2^ for decoding the Bayes-optimal prior from eye position (using sessions in which video was available and recordings were performed in VISp and LGd, *N* = 44 region sessions, Fig. S10d). Following the same approach as for the body posture and motion features, we then decoded the prior residual (Bayes-optimal prior minus Bayes-optimal prior estimated from eye position) from neural activity and found that the residual decoding *R*^2^ was correlated with the neural decoding *R*^2^ (Pearson correlation *R=0*.*8, p=7E-11*, N=44 region sessions Fig. S10d). Therefore, the prior signals found in VISp and LGd did not simply reflect subtle changes in eye position across blocks.

For the right map, combining Ephys and WFI, significance is assessed with the Benjamini-Hochberg procedure, correcting for multiple comparisons, with a conservative false discovery rate of 1%. 30.2% (left) and 24.0% (right) of the areas encode the prior significantly, at all levels of brain processing in both cases. For the full names of brain regions, see [online table](https://github.com/int-brain-lab/paper-brain-wide-map/blob/plotting/brainwidemap/meta/region_info.csv). **c**. Comparison of Ephys and WFI results for dorsal cortex. All areas significantly encode the Bayes-optimal prior in the WFI data (a region is deemed significant if its Fisher combined *p-*value is lower than 0.05). Blue means significant, orange not significant, grey not decoded because we lack quality-controlled data (see Methods), and white not decoded because we have no recordings or because it is out of the analysis’ scope (while we recorded from both hemispheres in WFI, we only decode from the left one here because we only recorded from the left hemisphere in Ephys). **d**. The corrected *R*^2^ for Ephys and WFI are significantly correlated (Spearman correlation *R*=0.52, *p*=0.0024, N=32 regions). Each dot corresponds to one region of the dorsal cortex (see Fig. S5b for color scheme). Significant and non-significant Ephys *R*^2^ were included in this analysis. **e**. Proportion of right choices on zero contrast trials as a function of the cross-validated decoded Bayes-optimal prior of the stimulus appearing on the right, estimated from the neural activity: higher values of the decoded prior are associated with greater proportion of right choices (See Methods for more details). Left: Ephys, Right: WFI. **f**. The corrected *R*^2^ values for decoding the prior from neural activity are correlated with the corrected *R*^2^ values for decoding the residual prior (Bayes-optimal prior minus Bayes-optimal prior decoded from DLC features). This correlation implies that the Bayes-optimal prior decoded from neural activity cannot be explained simply by the motor features extracted by DLC (see Fig. S5a for color scheme). **g**. Right: Granger graph at the Cosmos level in Ephys (p < 0.05 Bonferroni corrected; see Methods). This graph reveals a bidirectional flow of prior information between subcortical and cortical regions. Left: Granger directed pairs targeting VISp in WFI (p < 0.05 Bonferroni corrected). We observe a significant feedback flow of prior information from higher-order areas (highlighted with grey circles) to early sensory regions. **h**. Percentage of instances where a significant directed pair forms a loop of size 3 (orange dashed line), averaged across sessions in WFI (top) and Ephys (below). The flow of prior information forms more loops than what would be expected to occur by chance (blue null distribution).

Our decoding analysis reveals a robust, distributed representation of the Bayes-optimal prior throughout the brain, suggesting a complex network of information flow. To investigate the dynamics of the prior information network, we conducted a Granger Causality analysis during the ITI, between the time series of the decoded prior from one brain region and that of another (see Methods and Supp Fig. S11). This analysis revealed several key findings: 1) The flow of prior information between brain areas is significantly greater than expected by chance (Supp Fig. S11a), 2) This prior flow includes comprehensive communications across the entire brain, from subcortical to cortical areas and vice versa (Fig. 2g left panel), 3) It includes significant feedback connections from higher-order areas to early sensory areas (Fig. 2g right panel), and 4) There is a higher prevalence of loops within this communication network than would be anticipated by chance (Fig. 2h), including between higher-order and early sensory areas (Supp Fig. S11e). These results collectively highlight a loopy and intricate inter-area communication of prior information within the brain.

### Post stimulus prior

We also decoded the Bayes-optimal prior during the 100 ms interval after stimulus onset and found similarities between the encoding of the prior before and after stimulus onset. To avoid confounding the prior with the stimulus identity, two variables that are highly correlated (Spearman correlation *R*=0.40, *p*<1E-308), we first trained a linear decoder of signed contrast from neural activity in each region. We used the output of this decoder to fit two neurometric curves (proportion of decoded right stimulus as a function of contrast, see Methods) conditioned on the Bayes-optimal prior being above 0.7 or below 0.3. We then computed the vertical displacement of the fitted neurometric curves for zero contrast. If an area encodes the prior beyond the stimulus, we expect a shift between these two curves (see Fig. 3a for an example). The null distribution was generated using the pseudosession method previously described. Note that the same analysis can be performed during the ITI, though in this case the neurometric curves are expected to be flat (Fig. 3b) which is indeed what we observed (Fig. S12). This approach allows us to separate the encoding of the prior from the encoding of the stimulus; however, it is possible that some of our results are related to the emergence of the animal’s choices since the animals can respond in less than 100 ms on some trials (The International Brain Laboratory et al., 2021).

**Figure 3:**
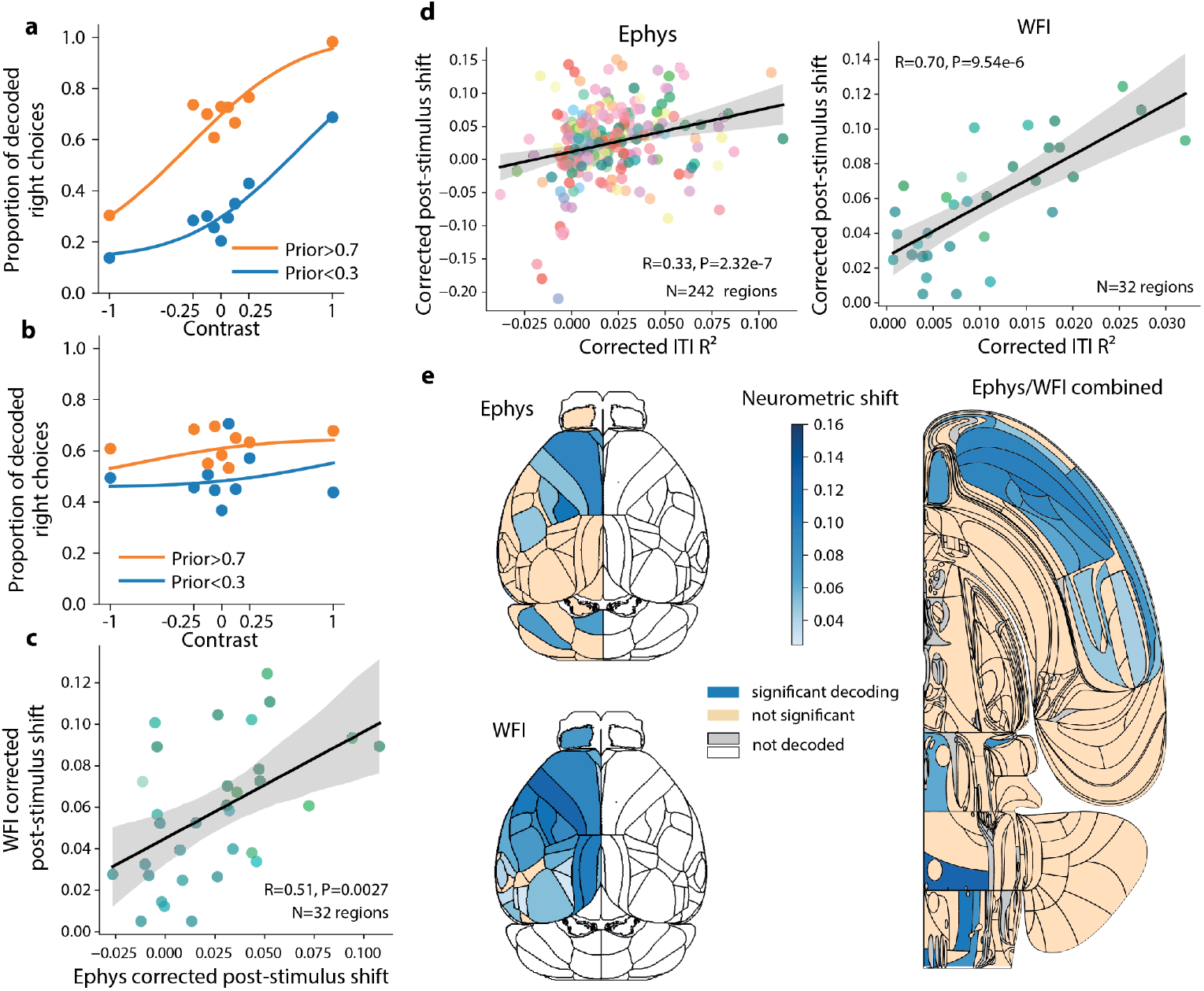
Encoding of the prior across the brain during the post-stimulus period. **a**. Example of neurometric curves for post-stimulus period from a SCm recording. **b**. Same as in **a** for the ITI period. **c**. Correlation between shifts from Ephys and WFI. Each dot corresponds to one cortical region. **d**. The post-stimulus shifts are correlated with the ITI *R*^2^ for both Ephy and WFI (see Fig. S5a for color scheme). **e**. Left: Comparison of corrected post-stimulus neurometric shift of Ephys and WFI for dorsal cortex (a region is deemed significant if its Fisher combined *p-value* is below 0.05). Right: Swanson map of corrected *R*^2^ averaged across Ephys and WFI for areas that have been deemed significant given both data sets (using Fisher’s method for combining *p*-values), and after applying the Benjamini-Hochberg correction for multiple comparisons.

Using this approach, we found that we can detect the prior significantly from 17.8% (43/242) and 84.4% (27/32) of areas during the post-stimulus period for Ephys (Fig. S13a) and WFI respectively (Fig 3e). When applying this methodology to the ITI, we found smaller percentages than with direct decoding in part because this neurometric shift measure is less sensitive (in the ITI, only 15.7% of regions for Ephys and 93.8% for WFI are significant for the Bayes-optimal prior when using the neurometric shift on Ephys/WFI data, versus 30.2% and 100% respectively for conventional decoding). As was the case during the ITI, we found that the Ephys and WFI post-stimulus shifts were correlated (Spearman correlation *R*=0.51, *p*=0.0027, N=32 Fig. 3c). Moreover, the post-stimulus neurometric shift is correlated with the *R*^2^ obtained in the same areas during the ITI period (Spearman correlation *R=0*.*33, p=2*.*32E-7*, N=242 for Ephys, *R*=0.70, *p*=9.54E-6, N=32 for WFI Fig. 3d). In other words, areas encoding the prior in the ITI also tend to do so during the post-stimulus period. This was confirmed by comparing the shifts during the post-stimulus and ITI periods, which were also found to be correlated (Fig. S13b,c).

We obtained similar results when merging the Ephys and WFI data into a single map (using Fisher’s method to combine p-values across Ephys and WFI) and applying the Benjamini-Hochberg correction for multiple comparisons with a false discovery rate of 1% (11.2% of significant regions, 27/242, Fig. 3e). Importantly, as observed during the ITI, areas encoding the prior were found at all levels of brain processing.

In addition, we explored whether regions encoding the stimulus also encoded the prior, as would be expected if these regions are involved in inferring the posterior distribution over the stimulus side. We found that the corrected *R*^2^ for the stimulus decoding was indeed correlated with the corrected *R*^2^ for the Bayes-optimal prior decoding (Spearman correlation *R*=0.29, *p*=2.4e-5, N=201 regions from BWM analysis, (International Brain Lab et al., 2023) Fig. S14a). Moreover, among the 40 areas that were found to encode the stimulus significantly, 25 also encoded the prior significantly, including, once again, areas at all levels of brain processing (e.g., LGd, VISp, SCm, CP, MOs, ACAd, Fig. S14b).

### Decoding the action kernel prior

So far we have established that mice leveraged the block structure and that the Bayes-optimal prior can be decoded from the neural data at all levels of brain processing. However, it remains to be seen whether the animals truly compute the Bayes-optimal prior or, perhaps, use heuristics to compute a subjective, approximate, prior (Schaeffer et al., 2020).

To address this, we developed several behavioral models and used session-level Bayesian cross-validation followed by Bayesian model selection (Stephan et al., 2009) to identify the one that fits the best (see Methods). This analysis suggests that most animals on most sessions estimate what we will refer to as the action kernel prior, which is obtained by calculating an exponentially weighted average of recent past actions (Fig. 4a). The action kernel prior explains the choices of the mice better than the Bayes-optimal prior and than models of behavioral strategies that calculate an exponentially weighted average of recent stimuli (the ‘stimulus kernel’), or assume a one step repetition bias, or a multi-step repetition bias (Urai et al., 2019) or the presence of positivity and confirmation biases (Palminteri & Lebreton, 2022) (see supp Fig. S15a and Supplementary Information). Consistent with the action kernel model, mice updated their subjective prior on the first 90 unbiased trials, even though the true prior is set to 0.5 during that phase (supp Fig S15d). Additionally, mice relied on more than just zero contrast trials to update their subjective prior (supp Fig S15b,c). The decay constant of the exponential action kernel had a median of 5.45 trials across all animals (Fig. 4b, blue histogram), similar to the decay constant of recovery after block switches (4.97 trials, Fig. 1c and Fig. S1). Remarkably, this is close to the value of the decay constant which maximizes the percentage of correct responses, given this (suboptimal) form of prior, losing only 1.9% compared to the performance of the Bayes-optimal version (Fig. 4b). These curves are obtained by simulating the action kernel by varying the decay constant while keeping all other parameters at their best-fitting values.

**Figure 4:**
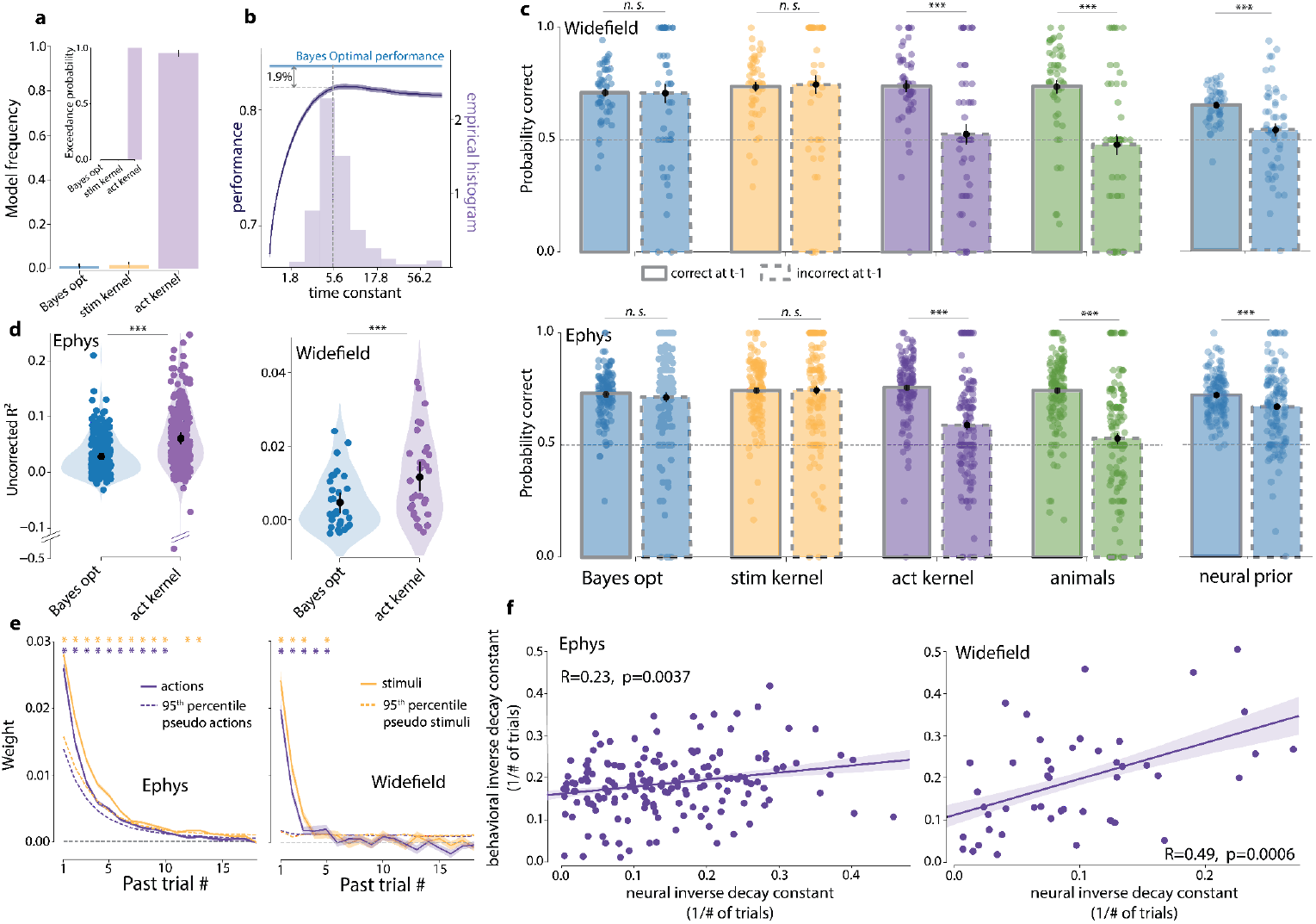
Action kernel prior. **a**. Model frequency (the posterior probability of the model given the subjects’ data) and exceedance probability (the probability that a model is more likely than any other models (Stephan et al., 2009)) for three models of the mice’s subjective prior. Across all sessions and all animals, the best model involves filtering the animal’s last actions with an exponential kernel. Act kernel: action kernel model, Bayes Opt: Bayes-optimal model, Stim Kernel: stimulus kernel model. **b**. Light purple: Histogram of the decay constant for the action kernel model across all sessions and animals. The median (dashed line) is 5.45 trials. Dark purple: probability of correct responses of the action kernel model as a function of the decay constant. The median time constant aligns closely with the optimal performance achievable with the action kernel model, only 1.9% below the Bayes-optimal performance (light blue horizontal line). **c**. Performance on zero contrast trials conditioned on whether the previous action is correct or incorrect for various behavioral models and for the animals’ behavior. Right: same analysis for a simulated agent using the Bayes-optimal prior decoded from neural data (Neural prior) to generate decisions. The decrease in performance between correct and incorrect previous trials for the neural prior suggests that the action kernel model best accounts for neural activity, which is consistent with behavior. Top: WFI. Bottom: Ephys. (*** *p*<0.001, *n*.*s*. not significant with a 2-tailed signed-rank paired Wilcoxon test) **d**. Uncorrected *R*^2^ is higher when decoding the action kernel prior during the ITI than when decoding the Bayes-optimal prior, for both Ephys and WFI (2-tailed signed-rank paired Wilcoxon test for Ephys *t*=2230, *p*=2.6E-30, N=242 regions, and widefield *t*=13, *p*=4.1E-8, N=32 regions). **e**. Average weight of the previous actions (purple) and previous stimuli (yellow) on the decoded Bayes-optimal prior, estimated from neural activity. Ephys: results based on all neurons, and averaged across sessions. WFI: results based on all pixels and averaged across sessions. Stars indicate weights that are significantly different from the pseudosession null distribution. Dashed lines show the 95th percentile of the null distribution. **f**. Correlation between the neural inverse decay constants, obtained by estimating the temporal dependency of the neural signals with respect to previous actions, and the behavioral inverse decay constants, obtained by fitting the action kernel model to the behavior. Neural and behavioral inverse decay constants are correlated for both Ephys (left, Pearson correlation *R*=0.23, *p*=0.0037, N=164 sessions) and WFI (right, Pearson correlation *R*=0.49, *p*=0.0006, N=46 sessions). This analysis included only a subset of sessions—specifically, those involving units or pixels that are part of region-sessions significantly decoding the Bayes-optimal prior (see Methods).

If, as our behavioral analysis suggests, mice use the action kernel prior, then we should find that when we decode the prior inferred from the action kernel, *R*^2^ should be higher than when we decode the prior predicted by any other method. This is borne out by the data in both Ephys and WFI during the ITI (Fig. 4d). Unfortunately, however, and in contrast to the Bayes-optimal prior, we cannot determine which areas encode the action kernel prior significantly, because of the impossibility of generating a null distribution, as this would formally require having access to the exact statistical model of the animal behavior (see section ‘Assessing the statistical significance of the decoding of the action kernel prior’ in Methods).

To explore further whether neural activity better reflects the action kernel prior, as opposed to the stimulus kernel prior or the Bayes-optimal prior, we looked at changes in performance on zero contrast trials following correct and incorrect actions. When considering behavior within blocks, a decision-making agent using an action kernel prior should achieve a higher percentage of correct responses after a correct block-consistent action than an incorrect one, because, on incorrect trials, it updates the prior with an action corresponding to the incorrect stimulus side. Models simulating agents using either the Bayes-optimal prior or the stimulus kernel prior do not show this asymmetry since they perform their updates using the true stimulus, which can always be correctly inferred from the combination of action and reward (see also (Schaeffer et al., 2020)). Mice behavior showed the asymmetry in performance (Fig. 4c). To test whether the neural data shared this asymmetry, we decoded the Bayes-optimal prior from ITI neural activity and simulated the animal’s decision on each trial by selecting a choice according to whether the decoded prior was greater or smaller than 0.5 (i.e., assuming every trial had a zero contrast stimulus). We then asked whether the resulting sequence of hypothetical choices would show the asymmetry. If so, this is a property of the neural data since the predicted quantity, the Bayes-optimal prior, does not show the asymmetry. As shown in Fig. 4c, performance for both modalities, Ephys and WFI, were indeed higher following correct versus incorrect trials, thus strengthening our hypothesis that neural activity more closely reflects the action kernel prior.

Next, we tested the sensitivity of the decoded Bayes-optimal prior, estimated from neural activity, to previous actions (decoding the Bayes-optimal prior instead of the action kernel prior to allow us to test for statistical significance, see Methods). If the prior we estimate from neural activity reflects the subjective prior estimated from behavior, we should find that the neural prior is sensitive to the past 5-6 trials. Using an orthogonalization approach, we estimated the influence of past actions on the decoded Bayes-optimal prior and found that this influence extends at least to the past 5 trials in both Ephys and WFI (Fig. 4e, see Methods section titled ‘Orthogonalization’). A similar result was obtained when testing the influence of the past stimuli (Fig. 4e). These numbers are consistent with the decay constant estimated from behavior (5.45 trials). These results were obtained at the session level, by analyzing all available neurons. Furthermore, we analyzed single regions for which we had a large number of neurons recorded simultaneously (SCm, CP, VPM), or strong imaging signals (MOp, VISp, MOs). In all cases, we found that an asymmetry in the neural data following correct and incorrect choices as well as evidence that the Bayes-optimal prior decoded from these regions is influenced by the past 5-6 actions (Fig. S16).

These analyses also address one potential concern with our decoding approach. It is well known in the literature that animals keep track of the last action or last stimulus (Busse et al., 2011; Lak et al., 2020; Mendonça et al., 2020). It is therefore conceivable that our ability to decode the prior from neural activity is simply based on the encoding of the last action in neural circuits, which indeed provides an approximate estimate of the Bayes-optimal prior since actions are influenced by the prior (Fig. 1c). The fact that we observe an influence of the last 5-6 trials, and not just the last trial, rules out this possibility.

To test this even further, we estimated the temporal dependency of the WFI single pixel and Ephys single-unit activities on past actions directly and compared them to the behavioral sensitivity to past actions on the same sessions (both expressed in terms of neural learning rates, i.e., the inverse of the decay constants, see Methods). Note that this analysis tests whether the temporal dynamics of neural activity is similar to the temporal dynamics of the mouse behavior, defined by fitting the action kernel model, but without regressing first the neural activity against any prior. We found that the inverse decay constants of the neural activity are indeed correlated across sessions with the inverse decay constants obtained by fitting the action kernel model to behavior (Fig. 4f). Critically, this correlation goes away if we perform the same analysis using stimulus kernels instead of action kernels (Fig. S17). Moreover, these results established at a session level remained when accounting for the variability across mice (see Fig. S18).

We next examined the link between behavioral performance and specific brain regions by comparing their neural inverse decay constants with the behavioral inverse decay constants. Notably, associative areas like the secondary motor cortex and retrosplenial areas more closely mirrored these behavioral constants than the primary visual and motor cortex (fig S19a). We also observed that the correlation between behavioral and neural decay constants reflected the prior corrected R2 from the same regions, indicating that regions with higher prior decoding R^2^ scores best align with the animal’s cognitive strategies as measured by the action kernel lengths (fig S19b). This analysis was not extended to electrophysiology recordings due to the limited number of available sessions per region (see fig S7a,b).

## Discussion

To summarize, we report that mice bias their decisions nearly optimally according to their prior expectations. As we have seen, the subjective prior of the animals is based on previous actions, not previous stimuli, a result consistent with past studies in rodents (Bolkan et al., 2022) and primates (Lau & Glimcher, 2005). Interestingly, this subjective prior is encoded, at least to some extent, at all levels of processing in the brain, including early sensory regions (e.g., LGd, VISp), associative regions (ORBvl, ACAd, SCm) and motor regions (MOs, MOp, GRN). Moreover, a Granger analysis revealed the existence of reciprocal loops, communicating specifically the subjective prior between cortical and subcortical regions as well as between sensory and associative cortical areas. These findings lend further support to the hypothesis that information flows across the brain in a way that could support the sort of multidirectional inference apparent in Bayesian networks (Ackley et al., 1985; Berkes et al., 2011; Jardri et al., 2017; Lange & Haefner, 2022).

One might argue that what we call a ‘subjective prior’ might be better called ‘motor preparation’ in motor related areas, or a top-down ‘attentional signal’ in early sensory areas. Ultimately, though, what is important is not the term we use to refer to this signal, but rather that it has properties consistent with the subjective prior: 1-it is predictive of the animal’s choices, particularly on zero contrast trials (Fig. 2e), 2-it depends on previous choices (Fig. 4c), and 3-it reflects more than the last choice or last stimulus, but depends instead on the past 5-6 choices (Fig. 4e). As we have seen, the signals we have recovered throughout the mouse brain fulfill all of these properties.

There are several proposals in the literature as to how probability distributions might be encoded in neural activity. These include: linear probabilistic population codes (Ma et al., 2006), sampling based codes (Echeveste et al., 2020), other activity based codes (Dabney et al., 2020; Ganguli & Simoncelli, 2014; Sahani & Dayan, 2003; Schaeffer et al., 2020; Zemel et al., 1998), and the synaptic weights of neural circuits (Soltani & Wang, 2010). We note that our results are compatible with two requirements of linear probabilistic population codes (Ma et al., 2006; Walker et al., 2020): 1-the log odds of the Bayes-optimal prior is linearly decodable from neural activity (Fig. S5), and 2-changes in the Bayes-optimal prior from trial to trial are reflected in the population activity (Walker et al., 2020).

If the likelihood is also encoded with a linear probabilistic population code, having the prior in the same format would greatly simplify the computation of the posterior distribution, since it would simply require a linear combination of the neural code for the prior and likelihood. As it turns out, it is likely that the likelihood indeed relies on a linear probabilistic population code. Indeed, the neural code for contrast, which is the variable that controls the uncertainty of the visual stimulus in our experiment, has been shown to be compatible with linear probabilistic population code in primates (Ma et al., 2006).

Whether our results are also compatible with sampling based codes is more difficult to assess, as there is still a debate as to which aspects of neural activity correspond to a sample of a probability distribution (Echeveste et al., 2020; Haefner et al., 2016; Hoyer & Hyvärinen, 2003). Moreover, the fact that our prior follows a Bernoulli distribution, which is particularly simple, makes it harder to tease apart the various probabilistic coding schemes.

Ultimately, determining the exact nature of the neural code for the prior will require developing a neural model of Bayesian inference in a large, modular, loopy network - a pressing, remaining task. A critical foundation for this development is the remainder of the extensive data in the International Brain Laboratory brain-wide map (described in the companion paper). This provides a picture, at an unprecedented scale, of the neural processes underpinning decision-making, in which the prior plays such a critical part.

## Methods

The experimental methods concerning the International Brain Laboratory (IBL) data acquisition and preprocessing can be found in the methods of previous IBL publications. For a detailed account of the surgical methods for the headbar implants, see appendix 1 of (The International Brain Laboratory et al., 2021). For a detailed list of experimental materials and installation instructions, see appendix 1 of (Laboratory, International Brain, 2022). For a detailed protocol on animal training, see methods in (Laboratory, International Brain, 2022; The International Brain Laboratory et al., 2021). For details on the craniotomy surgery, see appendix 3 of (Laboratory, International Brain, 2022). For full details on the probe tracking and alignment procedure, see appendices 5 & 6 of (Laboratory, International Brain, 2022). The spike sorting pipeline used at IBL is described in detail in (Laboratory, International Brain, 2022). A detailed account of the widefield imaging data acquisition and preprocessing can be found in Christopher Krasniak’s PhD thesis (Krasniak, 2022).

For the electrophysiological data, we used the 2024 IBL public data release (Laboratory, International Brain, 2023), which is extensively explored in IBL’s latest preprint “Brain-wide neural activity during the IBL task”. It is composed of 699 recordings from Neuropixels 1.0 probes. One or two probe insertions were realized over 459 sessions of the task, performed by a total of 139 mice. For the widefield calcium imaging data, we used another dataset consisting of 52 recordings of the dorsal cortex, realized over 52 sessions of the task, performed by a total of 6 mice.

### Inclusion criteria for the analysis

#### Criteria for trial inclusion

All trials were included except when the animals did not respond to the stimulus (no movement, no response) or when the first wheel movement time (reaction time) was shorter than 80 ms or longer than 2s.

#### Criteria for session inclusion

All sessions were included except sessions with fewer than 250 trials (counting only included trials). 41 sessions in Ephys and 1 session in Widefield did not meet the criteria for the minimum number of trials.

#### Criteria for neural recording inclusion

An insertion was included in the analysis if it had been resolved, that is, if histology clearly revealed the path of the probe throughout the brain, as defined in (Laboratory, International Brain, 2022). A neuron, identified during the spike sorting process, was included if it passed 3 quality control criteria (amplitude > 50 μV ; noise cut-off < 20; refractory period violation). A region recorded along one or two probes was included in the analysis if there were at least 5 units across the session’s probes that passed the quality control (QC). For widefield imaging, we used all the image pixels and included a region recorded during a session in the analysis if there were at least 5 recorded pixels.

For the region-level analysis, after applying these criteria, we were left with 414 sessions for the Ephys dataset. Initially, we considered 418 (=459-41) sessions that had more than 250 included trials, but 4 of these did not have any recorded regions meeting the minimum number of units required. Our region-level analysis spans 242 brain regions, defined by the Allen Common Coordinate Framework (Wang et al., 2020), recorded by at least one included insertion. Our Ephys region-level analysis spans 2289 region-sessions, which are aggregated across sessions to give results at the region level. For the widefield dataset, we were left with 51(=52-1) sessions that had more than 250 included trials. They all had at least one recorded region meeting the minimum number of recorded pixels required. The imaging spans 32 regions of the dorsal cortex – which are included among the 242 regions decoded in the Ephys analysis, for a total of 1539 (=(51*32)-28-65) region-sessions. This total accounts for the fact that not every region was visible in all sessions, summing to 28 non-observed region-sessions. Additionally, 65 region-sessions were excluded because the regions recorded had fewer than 5 pixels

For the session-level analysis, neurons along the whole probes were used and most of the sessions in Ephys (457/459) had at least 5 recorded units that passed the quality control. Taking into account the session inclusion criteria, session-level analysis was performed on 416 sessions. All of the 51 widefield imaging sessions passed the minimal number of trials criteria and were thus included in the analysis.

#### Criteria for the embodiment analysis

Only sessions with available DLC features could be used for the embodiment prior analysis, which requires access to body position. For the Ephys data set, we analyzed the 171 sessions (out of 459) for which the DLC features met the quality criteria defined in (Laboratory, International Brain, 2022), and for which the other inclusion criteria were met. This resulted in a total of 806 region-sessions. Wide field imaging sessions were excluded from this analysis as no video recordings were available.

#### Criteria for the eye position analysis

Reliable tracking of eye position from video recordings was not possible for some sessions because of video quality issues. Thus, we recovered reliable eye position signals from 44 out of the 53 of sessions in which we had recorded from either VISp or LGd, the two regions for which we specifically analyzed the impact of eye position.

#### Joint decoding of DLC features and eye position signals

We performed the joint decoding of DLC features and eye position signals on the 124 sessions where the DLC features met the quality criteria and also where the eye position signals were reliable, for a total of 660 region-sessions.

#### Difference with the Brain Wide Map inclusion criteria

There were two key differences between our inclusion criteria and those used in the Brain Wide Map (International Brain Lab et al., 2023). Firstly, the Brain Wide Map only included regions that had at least two recording sessions, whereas we included regions irrespective of the number of recording sessions. Secondly, we excluded sessions that had fewer than 250 trials after applying trial inclusion criteria, a criterion not applied in the Brain Wide Map.

### Electrophysiological data

Spike counts were obtained by summing the spikes across the decoding window for each neuron and included trial. If there were *U* units and *T* trials, this binning procedure resulted in a matrix of size *U* × *T*. For the intertrial interval, the time window for the main decoding was (−600ms,-100ms) relative to stimulus onset, and for the post-stimulus window, it was (0ms,+100ms) relative to stimulus onset. We used L1-regularized linear regression to decode the Bayes-optimal prior from the binned spike count data, with the scikit-learn function *sklearn*.*linear_model*.*Lasso* (employing one regularization parameter). We used L1 for Ephys because it is more robust to outliers, which are more likely to occur in single cell recordings, notably because of drift. The Bayes-optimal prior was inferred from the sequence of stimuli for each session (see the *Behavioral models* method section and in the Supplementary Information). This decoding procedure yielded a continuous-valued vector of length T.

### Widefield imaging data

For the widefield calcium imaging data, we used L2-regularized regression as implemented by the scikit-learn function *sklearn*.*linear_model*.*Ridge* (one regularization parameter). We used L2 regularization instead of L1 for WFI data because L2 tends to be more robust to collinear features, which is the case across WFI pixels. We decoded the activity from the vector of the region’s pixels for a specific frame of the data. The activity is the change in fluorescence intensity relative to the resting fluorescence intensity ∆*F*/*F*. Data were acquired at 15 Hz. Frame 0 corresponds to the frame containing stimulus onset. For the intertrial interval, we used frame −2 relative to the stimulus onset. This frame corresponds to a time window whose start ranges from −198 to −132ms prior to stimulus onset, and whose end ranges from −132 to −66ms, depending on the timing of the last frame before stimulus onset. For the post-stimulus interval, we used frame +2, which corresponds to a time window whose start ranges from 66 to 132ms from stimulus onset, and extends to 132 to 198ms post onset. This interval is dependent on the timing of the first frame after stimulus onset, which can occur anytime between 0 and 66 ms following onset. If there are *P* pixels and *T* trials, this selection procedure results in a matrix of size P × T.

### Reversal curves

To analyze mouse behavior around block reversals, we plotted the reversal curves defined as the proportion of correct choices as a function of trials, aligned to a block change (Fig. 1d). These were obtained by computing one reversal curve per mouse (pooling over sessions) and then averaging and computing the standard error across the mouse-level reversal curves. For comparison purposes, we also showed the reversal curves for the Bayes-optimal model with a probability-matching decision policy. We did not plot standard errors but standard deviations in this case as there was no variability across agents to account for.

To assess differences between the mouse behavior and the agent that samples actions from the Bayes-optimal prior, we fitted the following parametric function to the reversal curves:

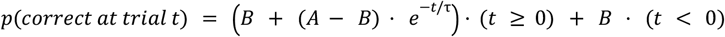

with *t* = 0 corresponding to the trial of the block reversal, τ the decay constant, *B* the asymptotic performance and A the drop in performance right after a block change.

We fitted this curve using only zero contrast trials, between the 5 pre-reversal trials and the 20 post-reversal trials. We restricted our analysis to the zero contrast trials to focus on trials where mice could only rely on block information to decide. This implied that we only used a small fraction of the data. To be precise, across the 459 sessions, we had an average of 10 reversals per session, and the proportion of zero contrast trials is 11.1%. Fitting only on the zero contrast trials around reversals led us to use around 28 trials per session, which accounts for around 3% of the behavioral data — when excluding the first 90 unbiased trials, the average session consists of 555 trials.

To make up for this limited amount of data, we used a jackknifing procedure for fitting the parameters. The procedure involved iteratively leaving out one mouse and fitting the parameters on the N-1=138 zero contrast reversal curves of the held-in mice. Results of the jackknifing procedure are shown in Supplementary Fig. 1.

### Nested cross validation procedure

Decoding was performed using cross-validated, maximum likelihood regression. We used the *scikit-learn* python package to perform the regression (Pedregosa et al., 2011), and implemented a nested cross-validation procedure to fit the regularization coefficient.

The regularization parameter was determined via two nested five-fold cross-validation schemes (outer and inner). We first describe the procedure for the Ephys data. In the ‘outer’ cross-validation scheme, each fold is based on a training/validation set comprising 80% of the trials and a test set of the remaining 20% (random “interleaved” trial selection). The training/validation set is itself split into five sub-folds (‘inner’ cross-validation) using an interleaved 80-20% partition. Cross-validated regression is performed on this 80% training/validation set using a range of regularization weights, chosen for each type of dataset so that the bounds of the hyperparameter range are not reached (see table 1). The regularization weight selected with the inner cross-validation procedure on the training/validation set is then used to predict the target variable on the 20% of trials in the held-out test set. We repeat this procedure for each of the five ‘outer’ folds, each time holding out a different 20% of test trials such that, after the five repetitions, 100% of trials have a held-out decoding prediction. For widefield imaging, the procedure is very similar but we increased the number of outer folds to 50 and performed a leave-one-trial-out procedure for the inner cross-validation (using the efficient *RidgeCV* native sklearn function). We did this because the number of features in widefield (number of pixels) is much larger than in Ephys (number of units): around 167 units in average in Ephys when decoding on a session-level from both probes after applying all quality criteria, vs around 2030 pixels on a session-level in widefield when decoding from the whole brain.

**table 1:** hyperparameter grid for the cross-validation, different for each kind of decoding modality.

| Type of decoding | regularization coefficient range |
| --- | --- |
| Ephys | $\{10^{-5}, 10^{-4}, 10^{-3}, 10^{-2}, 10^{-1}\}$ |
| widefield | $\{10^{-5}, 10^{-4}, 10^{-3}, 10^{-2}\}$ |
| DLC features | $\{10^{-4}, 10^{-3}, 10^{-2}, 10^{-1}, 1, 10, 100\}$ |
| Eye position | $\{10^{-4}, 10^{-3}, 10^{-2}, 10^{-1}, 1, 10, 100, 1000, 10000\}$ |
| DLC features + eye position | $\{10^{-4}, 10^{-3}, 10^{-2}, 10^{-1}, 1, 10, 100, 1000, 10000\}$ |
| Ephys neurometric | $\{10^{-5}, 10^{-4}, 10^{-3}, 10^{-2}, 10^{-1}, 1\}$ |
| widefield neurometric | $\{10^{-5}, 10^{-4}, 10^{-3}, 10^{-2}\}$ |

Furthermore, to average out the randomness in the outer randomization, we ran this procedure 10 times. Each run uses a different random seed for selecting the interleaved train/validation/test splits. We then report the median decoding score R^2^ across all runs. Regarding the decoded prior, we take the average of the predicted priors (estimated on the held-out test sets) across the 10 runs.

### Assessing statistical significance

Decoding a slow varying signal such as the Bayes-optimal prior from neural activity can easily lead to false positive results even when properly cross-validated. For instance, slow drift in the recordings can lead to spurious, yet significant, decoding of the prior if the drift is partially correlated with the block structure (Elber-Dorozko & Loewenstein, 2018; Harris, 2020). To control for this problem, we generated a null distribution of *R*^2^ values and determined significance with respect to that null distribution. This pseudosession method is described in detail in (Harris, 2020).

We denote *X* ∈ *R* ^*N*×*U*^, the aggregated neural activity for a session and *Y* ∈ *R*^*N*^ the Bayes-optimal prior. Here, *N* is the number of trials and *U* the number of units. We generated the null distribution from “pseudosessions”, i.e., sessions in which the true block and stimuli were resampled from the same generative process as the one used for the mice. This ensures that the time series of trials in each pseudosession shares the same summary statistics as the ones used in the experiment. For each true session, we generated *M*=1000 pseudosessions, and used their resampled stimulus sequences to compute Bayes-optimal priors *Y*_*i*_ ∈ *R*^*N*^, with *i* ∈ [1, *M*] the pseudosession number. We generated “pseudoscores” 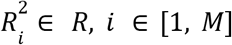 by running the neural analysis on the pair (*X, Y*_*i*_). The neural activity *X* is independent of *Y*_*i*_ as the mouse didn’t see *Y*_*i*_ but *Y*. Any predictive power from *X* to *Y*_*i*_ would arise from slow drift in *X* unrelated to the task itself. These pseudoscores 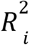 were compared to the actual score *R*^2^ obtained from the neural analysis on (*X, Y*) to assess statistical significance.

The actual *R*^2^ is deemed significant if it is higher than the 95th percentile of the pseudoscores distribution 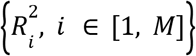. This test was used to reject the null hypothesis of no correlation between the Bayes optimal prior signal *Y* and the decoder prediction. We defined the p-value of the decoding score as the quantile of *R*^2^ relative to the null distribution 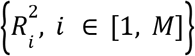.

For each region of the brain that we recorded, we obtained a list of decoding *p*-values where a *p*-value corresponds to the decoding of the region’s unit activity during one session. We used Fisher’s method to combine the session-level *p*-values of a region into a single region-level *p*-value (see the *Fisher’s method* section for more details).

For effect sizes, we computed a corrected *R*^2^, defined as the actual score *R*^2^ minus the median of the pseudoscores distribution, 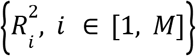. The corrected *R*^2^ of a region is the mean of the corrected *R*^2^ for the corresponding sessions.

### Choosing Between Pearson and Spearman Correlation Methods

In our statistical analyses, we prioritized using Spearman’s correlation when dealing with datasets that included outliers, since it is robust against non-normal distributions. In other cases, we opted for Pearson’s correlation to assess linear relationships.

### Fisher’s Method

Fisher’s method is a statistical technique used to combine independent *p*-values to assess the overall significance. It works by transforming each p-value into a chi-squared statistic and summing these statistics. Specifically, for a set of *p*-values (one per session given a region): *p*_1_, *p*_2_, *p*_3_, … Fisher’s method computes the test statistic:

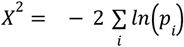

This statistic follows a chi-squared distribution with 2 · *N*_*sessions*_ degrees of freedom, 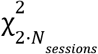, under the null hypothesis that all individual tests are independent and their null hypotheses are true. If the computed test statistic *X*^2^ exceeds a critical value from the chi-squared distribution, the combined *p*-value 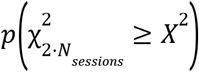 is considered significant and the null hypothesis is rejected.

### Cosmos Atlas

We defined a total of 10 annotation regions for coarse analyses. Annotations include the major divisions of the brain only: isocortex, olfactory areas, hippocampal formation, cortical subplate, cerebral nuclei, thalamus, hypothalamus, midbrain, hindbrain, cerebellum. For a detailed breakdown of the Cosmos atlas, refer to Figure S5.

### Granger causality

To understand how prior information flows between brain regions, we performed a Granger causality analysis on the Ephys and WFI data during the ITI.

For each Ephys session, we considered the neural activity from −600ms to −100ms before stimulus onset, segmented into 50ms bins, yielding 10 bins per region. For each bin, we predicted the Bayes-optimal from the neural activity using the native *LassoCV* sklearn function, with its default regularization candidates. This leads to a decoded Bayes-optimal prior for each region and bin. We then employed a Granger causality analysis to explore whether the prior information in some region Granger causes prior information in other regions. Granger analysis was run with the spectral connectivity python library from the Eden-Kramer lab https://github.com/Eden-Kramer-Lab/spectral_connectivity.

Given a directed pair of regions (e.g., from ACAd to VISp) within a session, the Granger analysis assigns an amplitude to each frequency in the discrete Fourier transform. We calculate an overall Granger score by session by averaging the amplitudes across frequencies (Lima et al., 2020).

To assess significance of the overall Granger score for a directed pair and session, we build a null distribution by applying our analysis to 1000 pseudosessions (see paragraph *Assessing statistical significance*). After decoding these pseudo-priors from neural activity for each region and bin, we perform Granger analysis on these decoded pseudo-priors. This creates 1000 pseudo Granger scores per directed pair and session. Significance is assessed by comparing the actual Granger score against the top 5% of the pseudo scores.

Granger analysis for the widefield data is very similar to that for Ephys. The main differences are to use the last nine frames before stimulus onset as individual bins and to employ the RidgeCV native function from sklearn for the decoding (see the *Widefield imaging data* method section).

Initially, we investigate whether communication between regions exceeds what might be expected by chance. To assess this, we analyze the percentage of significant directed pairs between two regions that significantly reflect the prior, we find an average of 71.6% in widefield and 35.9% in ephys across sessions. We then repeat this analysis for each session across 1000 pseudo-sessions. Subsequently, we assess whether the average percentage of significant pairs across sessions falls within the top 5% of the average percentages calculated from these pseudo-sessions, which indeed it does (see Fig. S11a).

Next, we explore whether the flow of prior information involves more loops than would typically occur by chance. Specifically, we assess whether triadic loops (A −> B −> A) within a session occur more frequently than expected. To evaluate this, we calculate the percentage of instances where a significant Granger pair results in a loop of size 3 for each session. We find that an average (across sessions) of 37.7% in widefield and 10.8% in electrophysiology exceed what would be anticipated by chance, confirming a higher prevalence of loops (see Fig. 2h and Fig. S11b).

To obtain Granger graphs at the region level, we apply Fisher’s method to combine the session-level *p*-values of a directed pair. Lastly, to construct the Granger causality graph at the Cosmos level, we further combine the *p*-values from each directed pair using Fisher’s method once again.

### Controlling for region size when comparing decoding scores across Ephys and WFI

With WFI data, the activity signal of a region has always the same dimension across sessions, corresponding to the number of pixels. To control for the effect of region size on the region *R*^2^, we performed linear regression across 32 recorded regions to predict the decoding *R*^2^ from the number of pixels per region. We found a significant correlation between *R*^2^ and the size of the regions (Fig S7d, *R*=0.82 *p*=9.1E-9). To determine whether this accounts for the correlations between Ephys and WFI *R*^2^ correlation (Fig. 2d), we subtracted the *R*^2^ predicted by region’s size from the WFI *R*^2^ and re-computed the correlation between Ephys *R*^2^ and these size-corrected WFI *R*^2^ (Fig. S7f).

### Number of recording sessions per region required to reach significance

For each region showing significance in prior decoding, we conducted a subsampling process to see how many recorded sessions were necessary on average to reach significance. A region *r* is associated with set of *N*_*r*_ sessions in which its activity was recorded. For a particular region, we can randomly subselect *N* ∈ [1, *N*_*r*_] sessions from this set and test the significance of the region given only these *N* prior decodings.

For each significant region and each possible number *N*, we repeated this procedure 1000 times, getting a distribution of *p*-values. We report 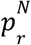, the median of the *p*-value distribution, as a measure of the significance for prior decoding of the region r given that only *N* recording sessions are available. For each number of available sessions *N*, we report the fraction of the total number of significant regions for which the statistic 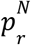 is less than 0.05, as a measure of the number of recordings per region required to reach back our obtained significance levels.

### Assessing the significance of decoding weights

Let *w*_*i*_ be the weight associated with the neuron *i* — where i ranges from 1 to *U*, with *U* the number of units, for the decoding of *U* neurons’ activity on a particular region-session. We determine whether the weight is significant by comparing it to the distribution of pseudoweights 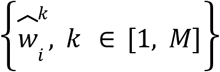 derived from decoding the *M* = 200 pseudosessions priors based on neural activity on that same region-session.

*w*_*i*_ is deemed significant if it is higher than the 97.5th percentile of the pseudoweights distribution or lower than the 2.5th percentile.

### Proportion of right choices as a function of the decoded prior

To establish a link between the decoded prior, estimated from the neural activity, and the mouse behavior, we plotted the proportion of right choices on zero contrast trials as a function of the decoded Bayes-optimal prior. For each electrophysiology (or widefield) session, we decoded the Bayes-optimal prior using all neural activity from that session. We focused this analysis on test trials, held-out during training, following the procedure described in the paragraph *Nested cross validation procedure*. For the main Fig. 2e, we then pooled over decoded priors for all sessions, assigned them to deciles and computed the associated proportion of right choices. In other words, we computed the average proportion of right choices on trials where the decoded prior is part of each decile.

To quantify the significance of this effect on a session-level (see Fig. S8), we additionally performed a logistic regression predicting the choice (right or left) as a function of the decoded prior. Let j be the session number, we predict the actions on that session 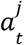 (with t the trial number) as a function of the decoded prior:

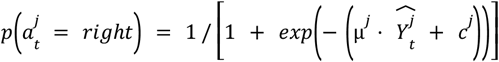

with 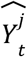 the cross-validated decoded Bayes-optimal prior, μ^*j*^ the slope (coefficient of the logistic regression associated with the decoded prior) and *c*^*j*^ an intercept. The logistic regression fitting was performed with the default sklearn *LogisticRegression* function, which uses L2 regularization on weights with regularization strength *C* = 1.

To assess the statistical significance of these slopes, μ^*j*^, we generated null distributions of slopes over *M=200* pseudosessions (pseudosessions are defined in the paragraph above titled *Assessing statistical significance*). For each pseudosession, we computed the slope of the logistic regression between proportion of correct choices as a function of the decoded pseudo Bayes-optimal prior. The decoded pseudo Bayes-optimal prior was obtained by, first, computing the pseudo Bayes-optimal prior for each pseudosession, and then using the neural data from the original session to decode this pseudo Bayes-optimal prior. The percentage of correct choice was more complicated to obtain on pseudosessions because it requires simulating the mice choices as accurately as possible. Since we don’t have a perfect model of the mice choices, we had to approximate this step with our best model, i.e., the action kernel model. We used the action kernel model fitted to the original behavior session and simulated it on each pseudosession to obtain the actions on each trial of the pseudosessions.

From the set of decoded pseudo Bayes-optimal priors and pseudoactions, we obtained *M* pseudoslopes 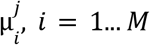 using the procedure described above. Because the mouse did not experience the pseudosessions or perform the pseudoactions, any positive coefficient 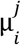 has to be the result of spurious correlations. Formally, to assess significance, we ask if the mean slope 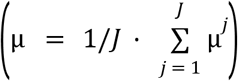 is within the 5% top percentile of the averaged pseudoslopes: 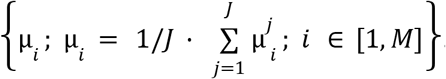. Fig. S8 shows this set of *M* averaged pseudoslopes as a histogram. The red vertical dashed line is the average slope μ.

When applying this null-distribution procedure in Ephys and WFI data, we find that the pseudoslopes in Ephys are much more positive than in WFI. This is due to the fact that spurious correlations in Ephys are mostly induced by drift in the Neuropixels probes, while WFI data barely exhibits any drift.

### Neurometric Curves

We used the same decoding pipeline described for the Bayes-optimal prior decoding to train a linear decoder of the signed contrast from neural activity in each region, for the intertrial interval [−600, −100]ms and post-stimulus [0, 100]ms intervals. There are 9 different signed contrasts {-1, −0.25, −0.125, −0.0625, 0, 0.0625, 0.125, 0.25, 1} where the left contrasts are negative and the right contrasts are positive. Given a session of *T* trials, we denote {*s*_*i*_}_*i*∈[1,*T*]_ the sequence of signed contrasts, 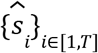 the cross-validated decoder output given the neural activity *X*, and {*p* _*i*_} _*i*∈[1,*T*]_ the Bayes-optimal prior. We defined two sets of trial indices for each session based on the signed contrast c and the Bayes-optimal prior: 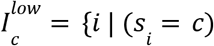 and *p*_*i*_ < 0. 5} and 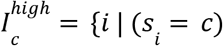 and *p*_*i*_ > 0. 5} corresponding to the trials with signed contrast *c* and a Bayes-optimal prior lower or higher than 0.5 respectively.

For these sets, we computed the proportions 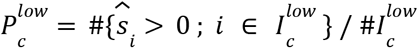 and 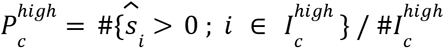. These are the proportions of trials decoded as right stimuli conditioned on the Bayes-optimal prior being higher or lower than 0.5. We fitted a *low prior* curve to 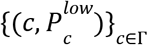 and a *high prior* curve to 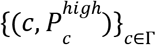, which we called neurometric curves. We used an *erf()* function from 0 to 1 with two lapse rates for the curves fit to obtain the neurometric curve:

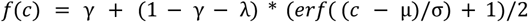

Where γ is the low lapse rate, λ is the high lapse rate, μ is the bias (threshold) and σ is the rate of change of performance (slope). Importantly, we assumed some shared parameters between the low-prior curve and the high-prior curve : γ, λ and σ are shared, while the bias μ is free to be different for low and high prior curves. This assumption of shared parameters provides a better fit to the data compared to models with independent parameters for each curve, as evidenced by lower Bayesian Information Criterion (BIC) scores during both pre-stimulus (ΔBIC = BIC(independent parameters) - BIC(shared parameters) = 6482 for ephys, ΔBIC = 822 for widefield) and post-stimulus periods (ΔBIC = 6435 for ephys, ΔBIC = 812 for widefield). We used the psychofit toolbox to fit the neurometric curves using maximal likelihood estimation (https://github.com/cortex-lab/psychofit). Finally, we estimated the vertical displacement of the fitted neurometric curves for the zero contrast *f*^*high*^ (*c* = 0) − *f*^*low*^ (*c* = 0), which we refer to as the neurometric shift.

We used the pseudosession method to assess the significance of the neurometric shift, by constructing a neurometric shift null distribution. *M*=200 pseudosessions are generated with their signed contrast sequences, which are used as target to linear decoder on the true neural activity. We fitted neurometric curves to the pseudosessions decoder outputs, conditioned on the Bayes-optimal prior inferred from the pseudosessions contrast sequences.

### Stimulus side decoding

To compare the representation of prior information across the brain to the representation of stimulus, we used the stimulus side decoding results from our companion paper (International Brain Lab et al., 2023). The decoding of the stimulus side was performed using cross-validated logistic regression with L1 regularization, on a time window of [0,100]ms after stimulus onset. Only regions with at least two recorded sessions were included, a criteria applied in companion paper. The right panel of Figure S15 is a reproduction from figure 5a of our companion paper.

### Embodiment

Video data from two cameras were used to extract 7 behavioral variables which could potentially be modulated according to the mice’s subjective prior (see (Laboratory, International Brain, 2022)): licking, whisking left & right, wheeling, nose position and paw position left & right. If we are able to significantly decode the Bayes-optimal prior from these behavioral variables during the [−600 ms, −100 ms] inter-trial interval, we say that the subject embodies the prior. For the decoding, we used L1-regularized maximum likelihood regression with the same cross-validation scheme used for neural data (see paragraph *Nested Cross-Validation procedure*). Sessions and trials are subject to the same QC as for the neural data, so that we decode the same sessions and the same trials as the Ephys session-level decoding. For each trial, the decoder input is the average over the ITI [−600, −100]ms of the behavioral variables. For a session of *T* trials, the decoder input is a matrix of size *T*x7 and the target is the Bayes-optimal prior. We use the pseudosession method to assess the significance of the DLC features decoding score *R*^2^.

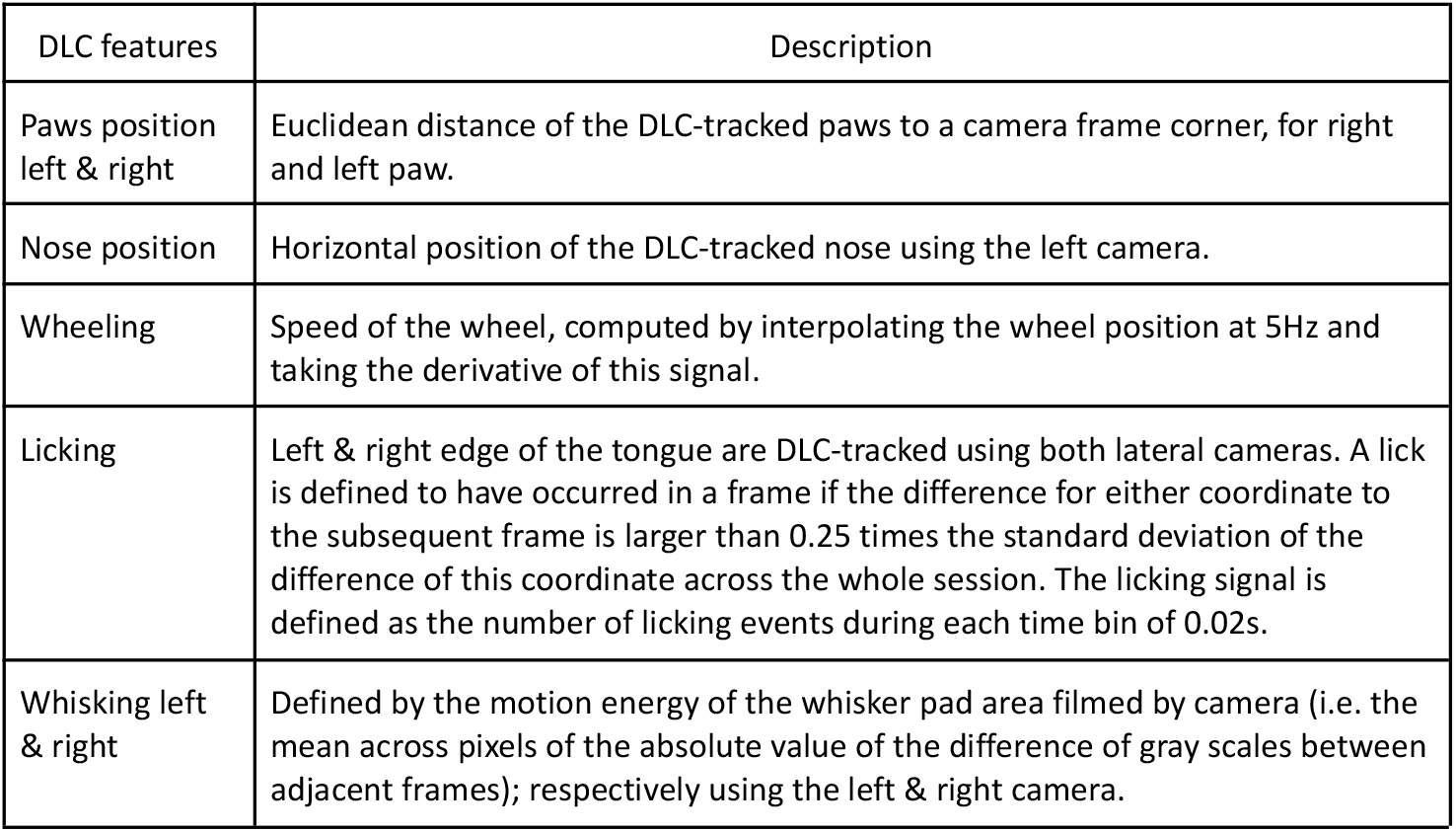

To investigate the embodiment of the Bayes-optimal prior signal, we compared session-level decoding of the prior signal from DLC regressors to region-session-level decoding of the prior signal from the neural activity of each region during the session.

### DLC Residual analysis

The DLC prior residual signal is the part of the prior signal which was not explained away by the DLC decoding, defined as the prior signal minus the prediction of the DLC decoding. We decoded this DLC prior residual signal from the neural activity, using the same linear decoding schemes as previously described.

### Eye position decoding

Video data from the left camera were used to extract the eye position variable, a 2D-signal corresponding to the position of the center of the mouse pupil relative to the video border. The camera as well as the mouse’s head were fixed. DeepLabCut was not able to achieve sufficiently reliable tracking of the pupils; therefore we used a different pose estimation algorithm (Biderman et al., 2023), trained on the same labeled dataset used to train DeepLabCut. For the decoding, we used L2-regularized maximum likelihood regression with the same cross-validation scheme used for neural data, during the [−600 ms, −100 ms] inter-trial interval.

The eye-position prior residual signal is the part of the prior signal which is not explained away by the eye position decoding, defined as the prior signal minus the prediction of the eye position decoding. We decode this eye position prior residual signal from the neural activity of early visual areas (LGd & VISp), using the same linear decoding schemes as previously described.

### Contribution of DLC and eye-position features to prior embodiment: feature importance

To assess the contribution of DLC and eye position features to prior embodiment, we performed a leave-one-out decoding procedure of the DLC + eye position features. There are five different types of DLC features : licking, wheeling, nose position, whisking and paws positions. Additionally, with the x and y coordinates of the eye position, we had a total of seven types of variables for which we individually performed a separate leave-one-out decoding analysis. The difference between the full decoding R^2^ and the leave-one-out decoding R^2^ is a measure of the importance of the knocked-out variable in the full decoding.

### Behavioral models

To determine the behavioral strategies used by the mice, we developed several behavioral models and used Bayesian model comparison to identify the one that fits best. We considered three types of behavioral models which differ as to how the integration across trials is performed (how the subjective prior probability that the stimulus will be on the right side is estimated based on history). Within a trial, all models compute a posterior distribution by taking the product of a prior and a likelihood function (probability of the noisy contrast given the stimulus side, see Supplementary information).

Among the three types of models of the prior, the first, called the *Bayes-optimal model*, assumes knowledge of the generative process of the blocks. Block lengths are sampled as follows:

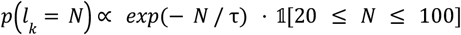

With *l*_*k*_ the length of block *k* and 𝟙 the indicator function. Block lengths are thus sampled from an exponential distribution with parameter τ = 60 and constrained to be between 20 and 100 trials.

When block *k* − 1 comes to an end, the next block *b*_*k*_, with length *l*_*k*_, is defined as a “right” block (where the stimulus is likely to appear more frequently on the right) if block *b*_*k*−1_ was a “left” block (where the stimulus was likely to appear more frequently on the left) and conversely. During “left” blocks, the stimulus is on the left side with probability γ = 0. 8 (and similarly for “right” blocks). Defining *s*_*t*_ as the side on which the stimulus appears on trial *t*, the Bayes-optimal prior probability the stimulus will appear on the right at trial *t, p* (*s*_*t*_ | *s* _1:(*t*−1)_) is obtained through a likelihood recursion (Scott, 2002).

The second model of the subjective prior, called the *stimulus kernel model* (Norton et al., 2019), assumes that the prior is estimated by integrating previous stimuli with an exponentially decaying kernel. Defining *s*_*t*−1_ as the stimulus side on trial *t* − 1, the prior probability that the stimulus will appear on the right π_*t*_ is updated as follows:

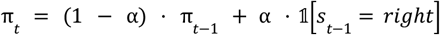

with π_*t*−1_ the prior at trial *t* − 1 and α the learning rate. The learning rate governs the speed of integration: the closer α is to 1, the more weight is given to recent stimuli *s*_*t*−1_.

The third model of the subjective prior, called the *action kernel model*, is similar to the *stimulus kernel model* but assumes an integration over previous chosen actions with, again, an exponentially decaying kernel. Defining *a*_*t*−1_ as the action at trial *t* − 1, the prior probability that the stimulus will appear on the right π_*t*_ is updated as follows:

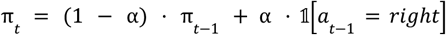

For the *Bayes-optimal* and *stimulus kernel* models, we additionally assume the possibility of capturing a simple autocorrelation between choices with immediate or multi-step repetition biases or choice- and outcome-dependent learning rate (Palminteri & Lebreton, 2022; Sugawara & Katahira, 2021). See Supplementary Information for more details on model derivations.

### Model comparison

To perform model comparison, we implemented a session-level Bayesian cross validation procedure. In this procedure, for each mouse with multiple sessions, we held out one session *i* and fitted the model on the held-in sessions. For each mouse, given a held-out session *i*, we fitted each model *k* to the actions of held-in sessions, denoted here as *A*^\*i*^ and obtained the posterior probability, *p* (θ_*k*_ |*A*^\*i*^, *m*_*k*_), over the fitted parameters θ_*k*_ through an adaptive Metropolis-Hastings (M-H) procedure (Andrieu & Thoms, 2008). 4 adaptive M-H chains were run in parallel for a maximum of 5000 steps, with the possibility of early stopping (after 1000 steps) implemented with the Gelman-Rubin diagnostic (Brooks & Gelman, 1998). θ_*k*_ typically includes sensory noise parameters, lapse rates and the learning rate (for *stimulus* and *action Kernel* models) - see Supplementary Information for the formal definitions of these parameters. Let {θ _*k,n*_; *n* ∈ [1, *N* _*MH*_]} be the *N*_*MH*_ samples obtained with the M-H procedure for model *k* (after discarding the burn-in period). We then computed the marginal likelihood of the actions on the held-out session, denoted here as *A*^*i*^.

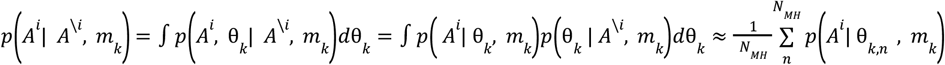

For each subject, we obtained a score per model *k* by summing over the log-marginal likelihoods *log p* (*A*^*i*^ | *A*^\*i*^, *m*_*k*_), obtained by holding out one session at a time. Given these subject-level log-marginal likelihood scores, we performed Bayesian model selection (Stephan et al., 2009) and reported the model frequencies (the expected frequency of the *k*-th model in the population) and the exceedance probabilities (the probability that a particular model *k* is more frequent in the population than any other considered model).

### Assessing the statistical significance of the decoding of the action kernel prior

Given that the action kernel model better accounts for the mice’s behavior, it would be desirable to assess the statistical significance of the decoding of the action kernel prior. Crucially, since assessing significance involves a null hypothesis (the neural activity is independent of the prior), a rigorous construction of the corresponding null distribution is key.

For the Bayes optimal prior decoding, constructing the null distribution is straightforward. It requires that we generate stimulus sequences with the exact same statistics as those experienced by the mice. We do this by simulating the same generative process used to generate the stimulus during the experiment, yielding what we called pseudosessions in previous sections.

However, for the action kernel prior (and contrary to the Bayes optimal model), we also need to generate action sequences with the same statistics as those generated by the animals. In turn, this would require a perfect model of how the animals make decisions. Since we lack such a model, we would need to come up with an approximation. There are multiple approximations that we could use, including:

1. Synthetic sessions, in which we use the action kernel model, using the parameters fitted to each mouse on each session, to generate fake responses. However, the action kernel model is not a perfect model of the animal’s behavior, it is merely the best model we have among the ones we have tested. Additionally, there could be some concerns about the statistical validity of using a null distribution, which assumes that the action kernel is the perfect model when testing for the presence of this same model in the mouse’s neural activity
2. Imposter sessions, in which we use responses from other mice. However, other animals are most unlikely to have used the exact same model/parameters as the mouse we are considering. This implies that the actions in these imposter sessions do not have the same statistics as the decoded session. There is indeed a large degree of between-session variability, as can be seen from the substantial dispersion in the fitted action kernel decay constants shown in Fig. 4b.
3. Shifted sessions, in which we decode the action kernel prior on trial M, using the recording on trial M+N, with periodic boundaries for the ‘edges’ (Harris, 2020). The problems here are two-fold. First, N must be chosen large enough such that the block structure of the shifted session is independent of the block structure of the non-shifted session. Because blocks are about 50 trials long, N must be large for the independence assumption to hold. This adds a constraint on the number of different shifted sessions that we can generate, leading to a poor null distribution with little diversity (made from only a few different shifted sessions). Second, it has been shown that there is within-session variability (Ashwood et al., 2022) such that when N is chosen to be large, we can not consider the shifted actions to have the same statistics as the non-shifted actions.

There may be other options. However, since they would all rely on approximations, the degree of statistical inaccuracy associated with their use would be unclear. We would not even know which one to favor, as it is hard to establish the quality of the approximations. Overall, we have access to the exact generative process to construct the null distribution for the Bayes optimal prior, versus only approximations for the action kernel prior. As a result, we decided to err on the side of caution and focus primarily on the Bayes optimal prior decoding whenever possible. In analyses involving animal behavior — such as in Fig. 2e, see the *Proportion of right choices as a function of the decoded prior* Method section, and Fig. 4e, see the *Orthogonalization* Method section — we had to rely on an approximation. For these, we employed the synthetic session approach to establish a null distribution.

### Orthogonalization

To assess the dependency on past trials of the decoded Bayes-optimal prior from neural activity, we performed stepwise linear regression as a function of the previous actions (or previous stimuli). The Bayes-optimal prior was decoded from neural activity on a session-level, thus considering the activity from all accessible cortical regions in WFI and all units in Ephys.

The stepwise linear regression involved the following steps. We started by linearly predicting the decoded Bayes-optimal prior on trial *t* from the previous action (action on trial *t*-1), which allows us to compute a first-order residual, defined as the difference between the decoded neural prior and the decoded prior predicted by the last action. We then used the second-to-last action (action at trial t-2) to predict the first-order residual, to then compute a second-order residual. Next, we predicted the second-order residual with the third-to-last action and so on. We use this iterative stepwise procedure in order to take into account possible autocorrelations in actions.

The statistical significance of the regression coefficients is assessed as follows. Let us use *T*^*j*^ to denote the number of trials of session 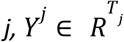 the decoded Bayes-optimal prior, and 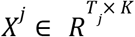 the chosen actions, where K is the number of past trials considered in the stepwise regression. When running the stepwise linear regression, we obtain a set of weights 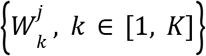, with 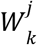 the weight associated with the *k*^th^-to-last chosen action. We test for the significance of the weights for each step k, using as a null hypothesis that the weights associated with the *k*^th^-to-last chosen action are not different from weights predicted by the ‘pseudosessions’ null distribution.

To obtain a null distribution, we followed the same approach as in the section entitled *Proportion of right choices as a function of the decoded prior*. Thus, we generated decoded pseudo Bayes-optimal priors and pseudoactions. For each session, these pseudovariables are generated in the following way: first, we fitted the action kernel model (our best fitting-model) to the behavior of session *j*. Second, we generated *M* pseudosessions (see the *Assessing the statistical significance* section). Lastly, we simulated the fitted model on the pseudosessions to obtain pseudoactions. Regarding the decoded pseudo Bayes-optimal priors, we first infer with the Bayes-optimal agent, the Bayes-optimal prior of the pseudosessions, and second, we decoded this pseudoprior with the neural activity. For each session *j* and pseudo *i*, we have generated a decoded pseudo Bayes-optimal prior 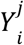 as well as pseudoactions 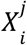. When applying the stepwise linear regression procedure to the couple 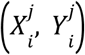, we obtain a set of pseudoweights 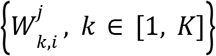. Because the mouse did not experience the pseudosessions or perform the pseudoactions, any non-zero coefficients 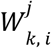 must be the consequence of spurious correlations. Formally, to assess significance, we ask if the average of the coefficients over sessions 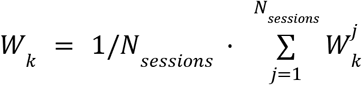 is within the 5% top percentile of 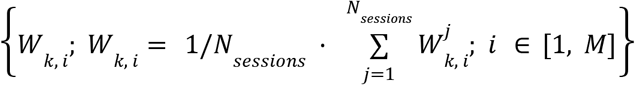.

The statistical significance procedure when predicting the decoded Bayes-optimal prior from the previous stimuli is very similar to the one just described for the previous actions. The sole difference is that, for this second case, we do not need to fit any behavioral model to generate pseudostimuli. Pseudostimuli for session *j* are defined when generating the *M* pseudosessions. Pseudoweights are then obtained by running the stepwise linear regression predicting the decoded pseudo Bayes-optimal prior from the pseudostimuli. Formal statistical significance is established in the same way as for the previous actions case.

When applying this null-distribution procedure to Ephys and WFI, we find that the strength of spurious correlations (as quantified by the amplitude of pseudoweights *W*_*k,i*_) for Ephys is much greater than for WFI data. This is due to the fact that spurious correlations in electrophysiology are mainly produced by drift in the Neuropixels probes, which is minimal in WFI.

### Behavioral signatures of the action kernel model

To study why the Bayesian model selection procedure favors the action kernel model, we sought behavioral signatures that can be explained by this model but not the others. Since the action kernel model integrates over previous actions (and not stimuli sides), it is a self-confirmatory strategy. This means that if an action kernel agent was incorrect on a block-conformant trial (trials where the stimulus is on the side predicted by the block prior), then it should be more likely to be incorrect on the subsequent trial (if it is also block-conformant). Other models integrating over stimuli, such as the Bayes-optimal or the stimulus Kernel model are not more likely to be incorrect following an incorrect trial, because they can use the occurrence or non-occurrence of the reward to determine the true stimulus side, which could then be used to update the prior estimate correctly. To test this, we analyzed the proportion correct of each session at trial *t*, conditioned on whether it was correct or incorrect at trial *t* − 1. To isolate the impact of the last trial, and not previous trials or other factors such as block switches and structure, we restricted ourselves to:

- zero contrast trials
- trial *t, t* − 1, and *t* − 2, had stimuli which were on the “expected”, meaning “block-conformant” side
- on trial *t* − 2, the mouse was correct, meaning that it chose the block-conformant action
- on trials that were at least 10 trials from the last reversal

### Neural signature of the action kernel model from the decoded Bayes-optimal prior

To test if the behavioral signature of the action kernel model discussed in the previous section is also present in the neural activity, we simulated an agent whose decisions are based on the cross-validated decoded Bayes-optimal prior and tested whether this agent also shows the same action kernel signature. The decoded Bayes-optimal prior was obtained by decoding the Bayes-optimal prior from the neural activity (see *Nested cross validation procedure* section) on a session-level basis, considering all available widefield pixels or electrophysiology units.

Note that if the decoded Bayes-optimal agent exhibits the action kernel behavioral signature, this must be a property of the neural activity since the Bayes-optimal prior on its own cannot produce this behavior.

The agent is simulated as follows. Let us denote *Y* ∈ *R*^*N*^ the Bayes-optimal prior with *N* is the number of trials. When performing neural decoding of the Bayes-optimal prior *Y*, we obtain a cross-validated decoded Bayes-optimal prior 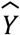. We define an agent which, on each trial, greedily selects the action predicted by the decoded Bayes-optimal prior 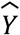, meaning that the agent chooses right if 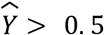, and left otherwise.

On sessions which significantly decoded the Bayes optimal prior, we then test whether the proportion of correct choices depends on whether the previous trial was correct or incorrect. We do so at the session level, applying all but one criteria of the behavioral analysis described previously in the *Behavioral signatures of the action kernel model* paragraph:

- trial *t, t* − 1, and *t* − 2, had stimuli which were on the “expected”, meaning on the “block-conformant” side
- on trial *t* − 2, the mouse was correct, meaning that it chose the block-conformant action
- on trials that were at least 10 trials from the last reversal

Note that, given that the neural agent uses the pre-stimulus activity to make its choice, we do not need to restrict ourselves to zero contrast trials.

#### Neural Decay Rate

To estimate the temporal dependency of the neural activity in Ephys and WFI, we assumed that the neural activity was the result of an action kernel (or stimulus kernel) integration and fitted the learning rate (inverse decay rate) of the kernel to maximise the likelihood of observing the neural data.

We first describe the fitting procedure for widefield imaging data. Given a session, let us call *X*_*t,n*_ the widefield calcium imaging activity of the *n*-th pixel for trial *t*. Similarly to the procedure we used for decoding the Bayes-optimal prior, we took the activity at the second-to-last frame before stimulus onset. We assumed that *X*_*t,n*_ is a realization of Gaussian distribution with mean *Q*_*t,n*_ and with standard deviation σ_*n*_, *X*_*t,n*_ ∼ *N*(*Q*_*t,n*_, σ_*n*_). *Q*_*t,n*_ was obtained through an action kernel (or stimulus kernel) integration process:

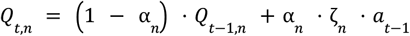

with α_*n*_ the learning rate, *a* _*t*−1_ ∈ {− 1, 1} the action at trial *t* − 1 and ζ_*n*_ a scaling factor. α_*n*_, ζ_*n*_ and σ_*n*_ are found by maximizing the probability of observing the widefield activity *p*(*X*_1:*T,n*_ |*a*_1:*T*_ ; α_*n*_, ζ _*n*_, σ_*n*_), with 1: *T* = {1, 2, … *T*} and *T* the number of trials in that session. If a trial is missed by the mouse, which occurs when reaction time exceeds 60 seconds (1.5% of the trials, see companion paper), *Q*_*t,n*_ is not updated. For the electrophysiology now, let us call *X*_*t,n*_ the neural activity of unit *n* at trial *t*. Similarly to what we did when decoding the Bayes-optimal prior, we took the sum of the spikes between −600 and −100ms from stimulus onset. We assumed here that *X*_*t,n*_ is a realization of a Poisson distribution with parameter *Q*_*t,n*_, *X*_*t,n*_ ∼ *Poisson*(*Q* _*t,n*_). *Q*_*t,n*_ was obtained through an action kernel (or stimulus kernel) integration process:

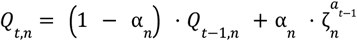

with α_*n*_ the learning rate and 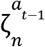 scaling factors, one for each possible previous action. Note that in this Ephys case, as *Q*_*t,n*_ can only be positive, two scaling factors are necessary to define how *Q*_*t,n*_ is adjusted after a right or left choice. 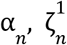 and 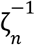 are found by maximizing the probability of observing the Ephys activity 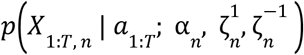, with 1: *T* = {1, 2, … *T*} and *T* the number of trials in that session. For electrophysiology, we added constraints on the units. Specifically, we only considered units 1-whose median (pre-stimulus summed) spikes was not 0, 2-with at least 1 spike every 5 trials and 3-where the distribution of (pre-stim summed) spikes was different when the Bayes-optimal prior is greater versus lower than 0.5 (significance is asserted when the p-value of a Kolmogorov Smirnov test was below 0.05).

To restrict our analysis to units (or pixels) which are likely to reflect the subjective prior, we only assessed those that are part of regions-sessions that significantly decoded the Bayes-optimal prior. Significance is assessed according to the pseudosession methodology (see the *Assessing statistical significance* section), which accounts for spurious correlations (which a unit-level Kolmogorov Smirnov test would not). Then, to obtain a session-level neural learning rate, we averaged across pixel-level or unit-level learning rates.

To compare neural and behavioral temporal timescales, we correlated the session-level neural learning rate with the behavioral learning rate, obtained by fitting the action kernel to the behavior.

In both Ephys and WFI, when considering that the neural activity is a result of the stimulus kernel, the calculations were all identical except replacing actions *a*_1:*T*_ with stimuli side *s*_1:*T*_.

This analysis (presented in Fig. 4f) makes the assumption that sessions could be considered as independent from another - an assumption which can be questioned given that we have a total of 459 sessions across 139 mice in electrophysiology and 52 sessions across 6 mice in widefield. To test the presence of the correlation between neural and behavioral timescales while relaxing this assumption, we developed a hierarchical model which takes into account the two types of variability, within mice and within sessions given a mouse. This model defines session-level parameters, which are sampled from mouse-level distributions, which are themselves dependent on population-level distributions. See Supplementary information for the exact definition of the hierarchical model. This hierarchical approach confirmed the session-level correlation between neural and behavioral timescales (see Fig. S18).

## Data availability

All the data that support the findings of the present study are available at https://int-brain-lab.github.io/iblenv/notebooks_external/data_release_brainwidemap.html and https://int-brain-lab.github.io/iblenv/notebooks_external/loading_widefield_data.html. Users are allowed to distribute, remix, adapt, and build upon the material in any medium or format, so long as attribution is given to the creator (data license CC-BY).

## Code availability

The code associated with this paper can be found in the following github repository: https://github.com/int-brain-lab/prior-localization/tree/main.

## Acknowledgements

This work was supported by grants from the Wellcome Trust (216324), the Simons Foundation, The National Institutes of Health (NIH U19NS12371601), the National Science Foundation (NSF 1707398), the Gatsby Charitable Foundation (GAT3708), and by the Max Planck Society and the Humboldt Foundation. We would like to thank the University of Geneva for providing computational resources and support that contributed to these research results.

## Supplementary Figures

**Figure S1.**
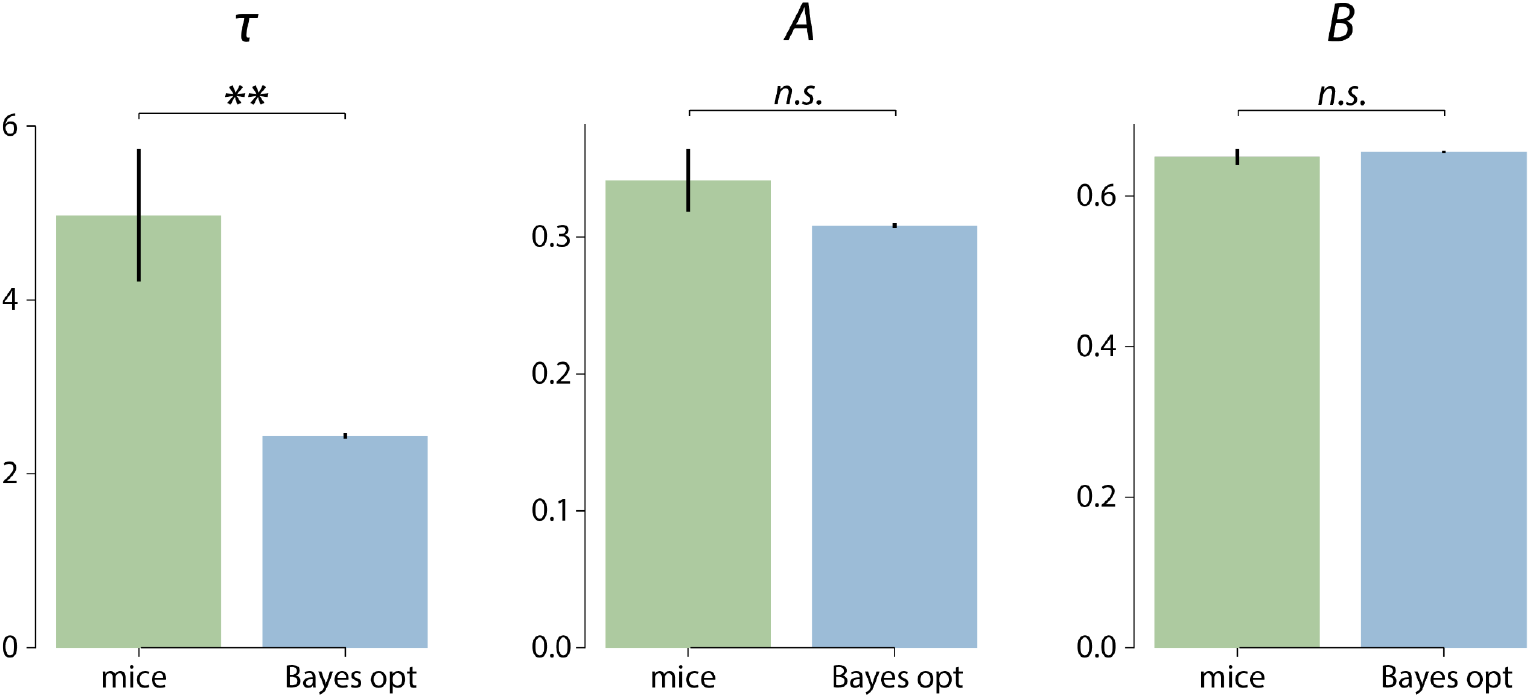
Histograms of the decay constant (τ), amplitude (*A*) and asymptote (*B*) of the zero contrast reversal curves across all mice. The parameters are obtained by fitting the following parametric curve: *p*(*correct at trial t*) = *B* on the zero contrast pre-reversal trials (the 5 trials before a block switch) and *p*(*correct at trial t*) = *B* + (*A* − *B*) · *e*^−*t*/τ^ on the zero contrast post-reversal trials (the 20 trials after a block switch). τ reflects the reversal timescale. To make up for the limited amount of available zero contrast reversal trials, we fit these curves using a jackknife procedure (see Methods). Bars and error bars indicate jackknife means ± SEM (jackknifing was applied on N = 139 mice). Mice have a significantly longer mean recovery decay constant than the Bayes-optimal observer (4.97 vs 2.43, 2-tailed paired t-test t=2.94, *p*=0.001), while the other parameters are not significantly different. (for *A:* t=1.43, *p*=0.15 and for *B*: t=-0.64, *p*=0.53) (** *p*<0.01, n.s. not significant)

**Figure S2.**
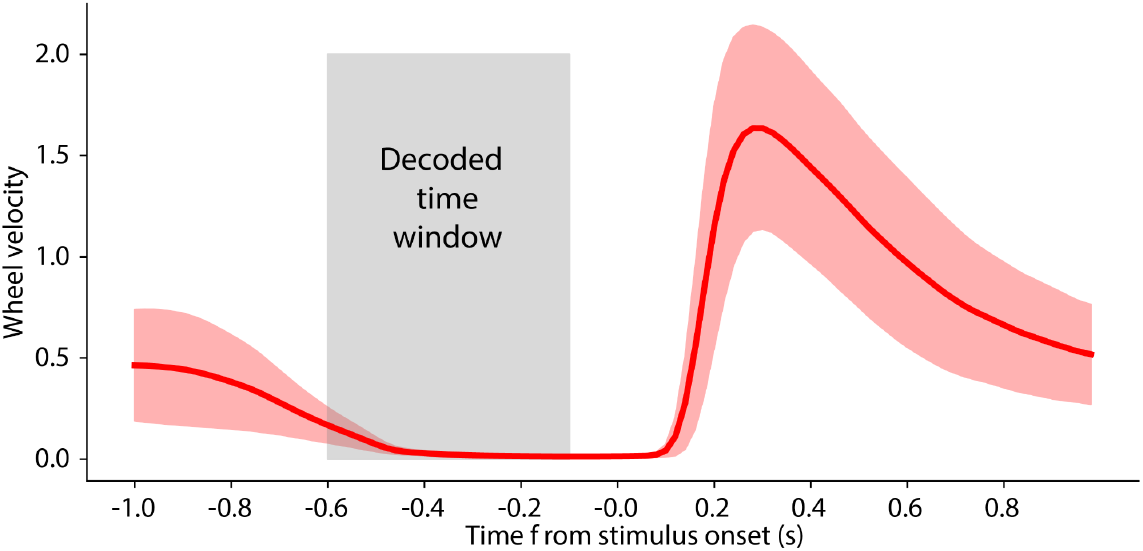
Average wheel speed averaged across sessions before and after stimulus onset. The decoded time window used for Ephys data is indicated in light gray. For WFI, the data was decoded on the second-to-last frame relative to the stimulus onset, corresponding to a time window that ranges from −198 to −132ms at the start to −132 to −66ms at the end, depending on the timing of the last frame before the stimulus onset (this last frame can occur anytime between −132 and −66ms before the stimulus onset).

**Figure S3.**
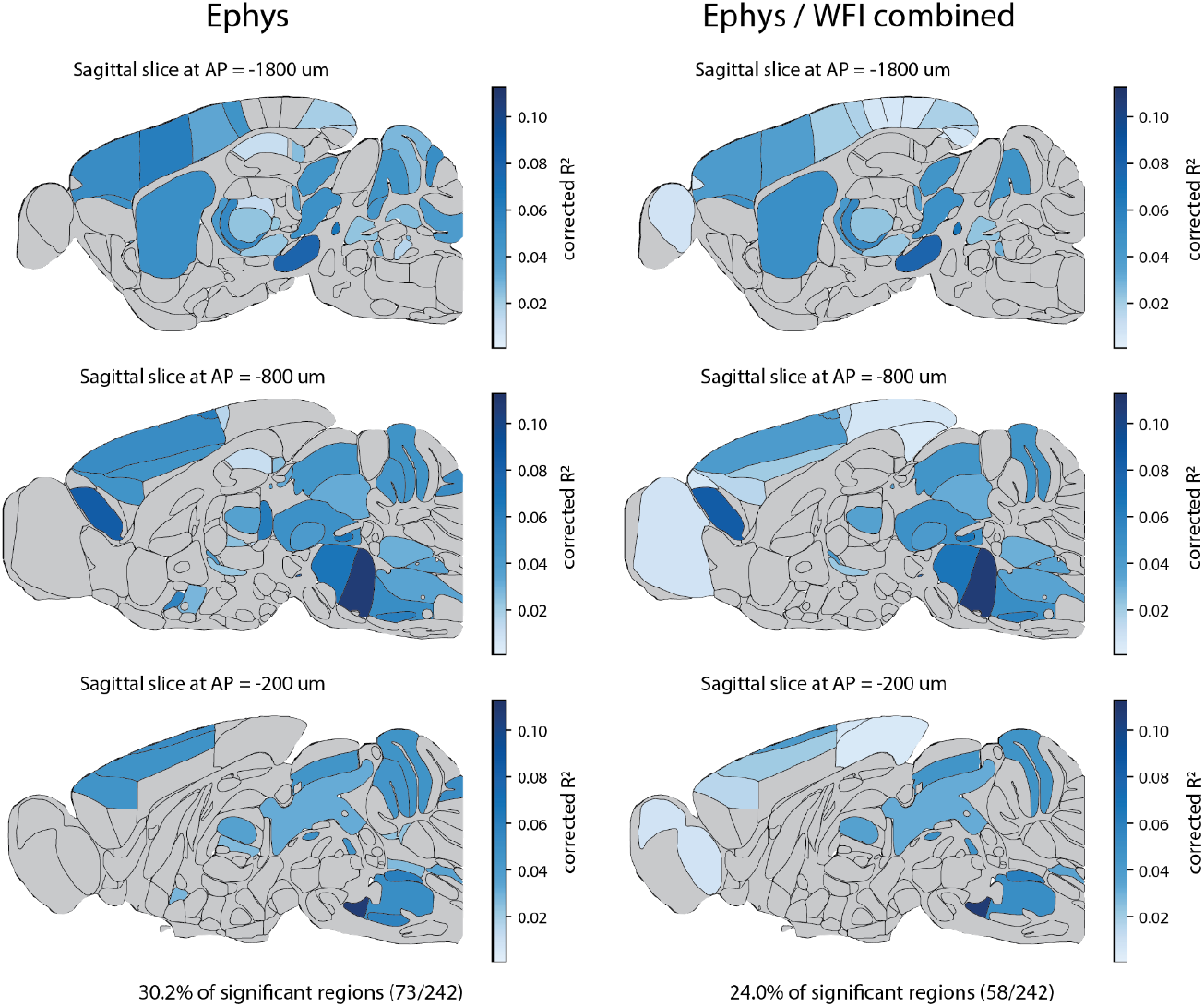
Encoding of the prior across the brain during the inter-trial interval. Sagittal slices corresponding to the main decoding figure presented Fig. 2b. Left: Ephys only. A region is deemed significant if the Fisher combined *p*-value is lower than 0.05. Right: Ephys and Widefield combined. Significance for regions is assessed with the Benjamini-Hochberg procedure, correcting for multiple comparisons, with a conservative false discovery rate of 1%.

**Figure S4.**
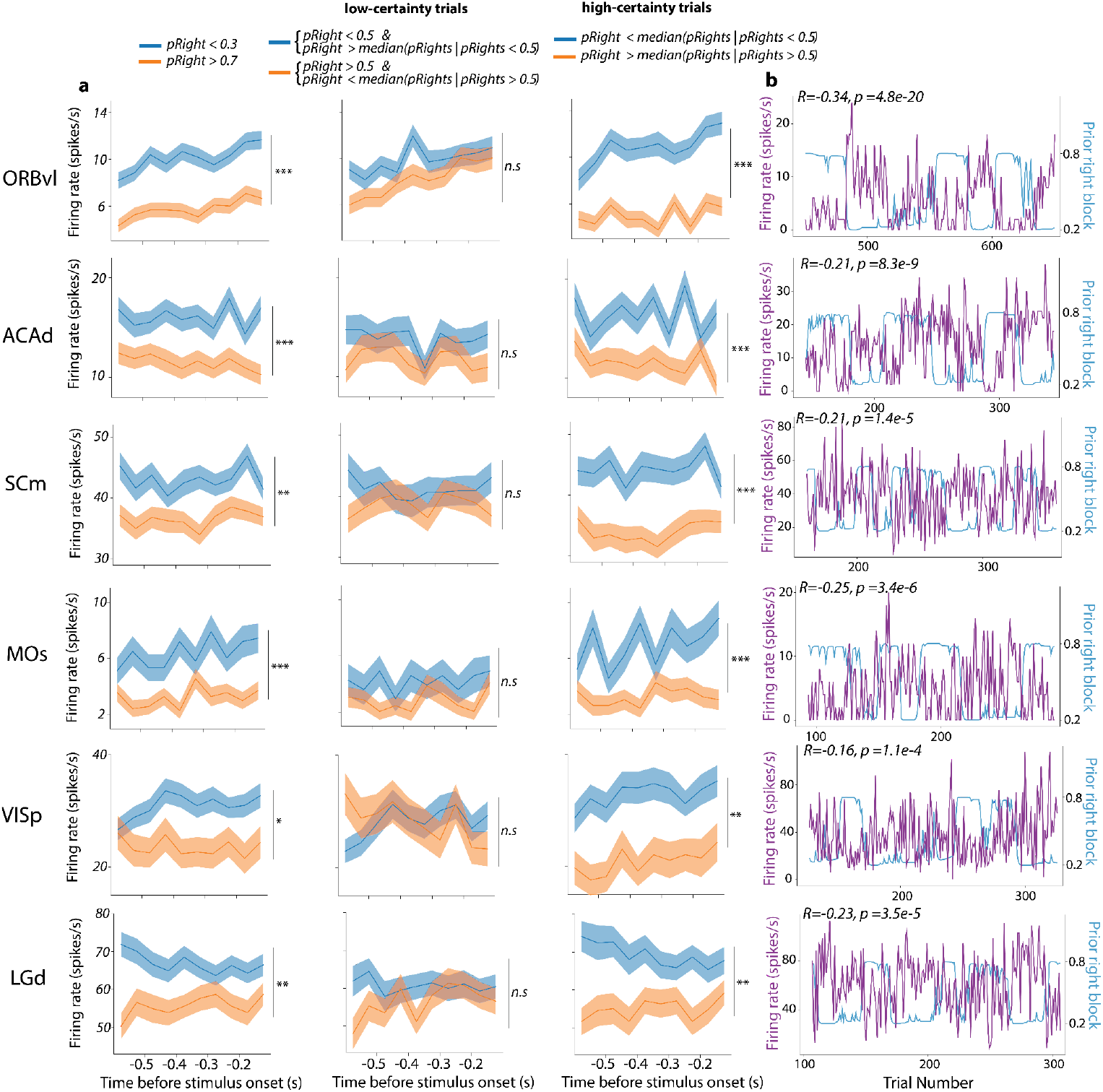
Six examples of neurons encoding the Bayes-optimal prior significantly (*** p<0.001, ** p<0.01, * p<0.05). **a**. Peri-Stimulus Time Histograms (PSTHs) segmented by trials throughout the session. Left column conditions on the Bayes-optimal prior for the right side being less than 0.3 (blue) vs greater than 0.7 (orange). The middle and right columns depict PSTHs for trials under conditions of low certainty (pRight close to 0.5) and high certainty (pRight far from 0.5), respectively. Significance is assessed by testing the difference between the trial wise firing rates (averaging across time bins) of “left” (blue) and “right” (orange) trials with a two-sample Kolmogorov-Smirnov test. **b**. Spike counts of the neurons (purple line) during the intertrial interval in the [−600, −100] millisecond time window before stimulus onset, along with the Bayes-optimal prior (blue) for a subset of trials within the session (Spearman correlations of the full session are reported on the graphs). All neurons on this panel show a preference for the left side though, at the population level, we did not observe a bias for either the right or left side. Indeed, we examined the distribution of decoding weights and detected no discernible trend concerning the weight distribution. Testing the significance of the decoder weight in each region yielded adjusted p-values all above 0.2 (Wilcoxon test), after adjusting for multiple comparisons using the Benjamini-Hochberg correction. Additionally, a combined analysis of all weights from the six regions lead to the same conclusion (two tailed signed Wilcoxon test: t=16732, p-value=0.31).

**Figure S5.**
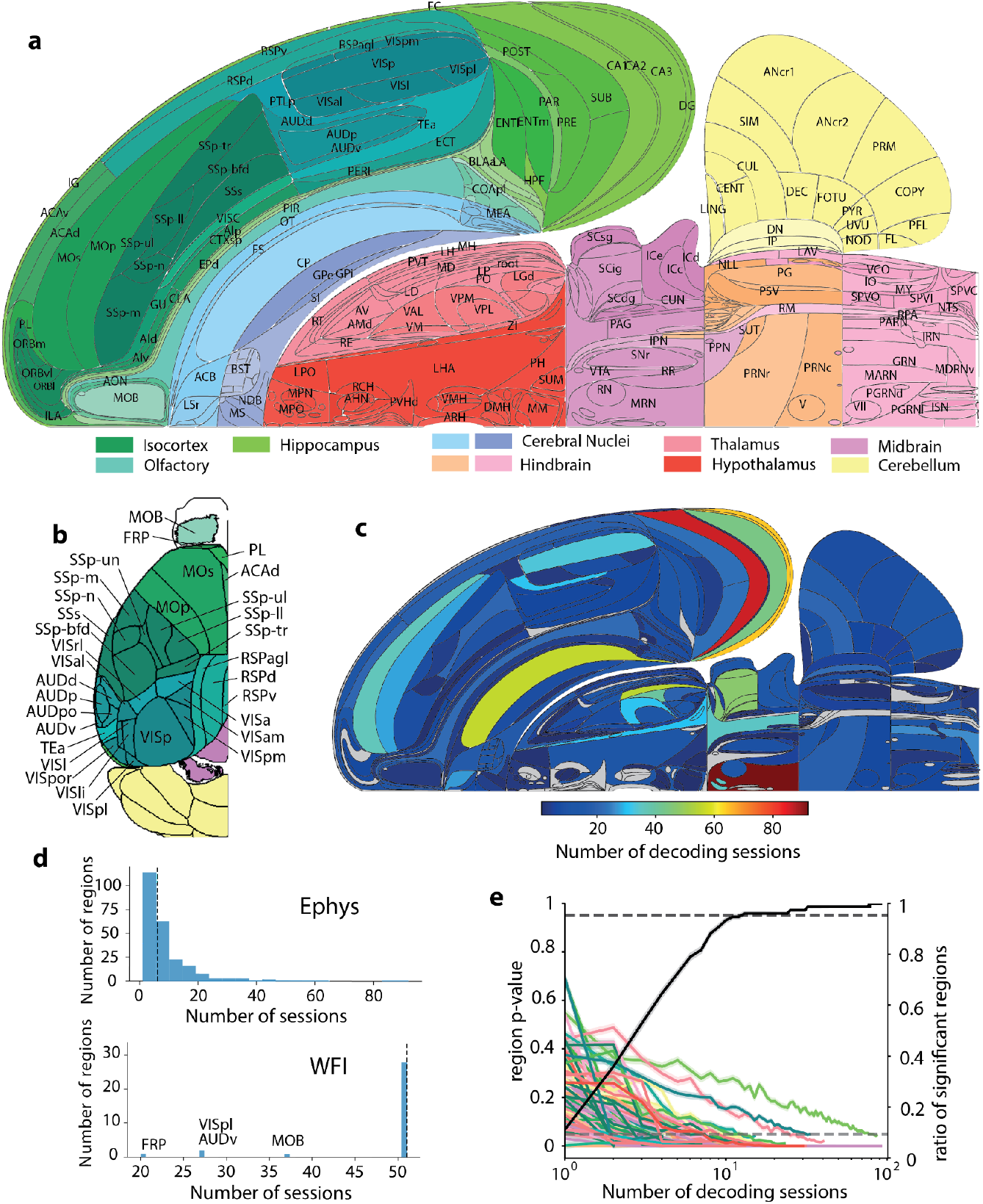
**a**. Swanson map color-coded to indicate distinct anatomical regions. **b**. Dorsal brain slice recorded with widefield imaging, utilizing the same color scheme for regional identification. **c**. Distribution of decoding sessions across different regions as mapped in the Swanson brain atlas. **d**. Histogram detailing the number of sessions per recording type: Electrophysiology (Ephys) and Widefield Imaging (WFI). The vertical lines indicate median values, with 6 sessions for Ephys and 51 for WFI. **e**. Dual-axis graph displaying the p-value of regional significance as a function of the number of decoding sessions (left axis) and the ratio of significant regions relative to the total number of sessions (right axis). It is estimated that approximately 10 sessions per region are necessary to identify 95% of significant regions highlighted in the main decoding analysis (refer to Methods section for more details). It’s important to recognize that this analysis has limitations: it assumes uniformity across recordings and regions without considering, e.g., variations in effect size or number of units per recording. Despite these limitations, we concentrated on the number of recordings because it is a primary factor that experimenters can directly control.

**Figure S6.**
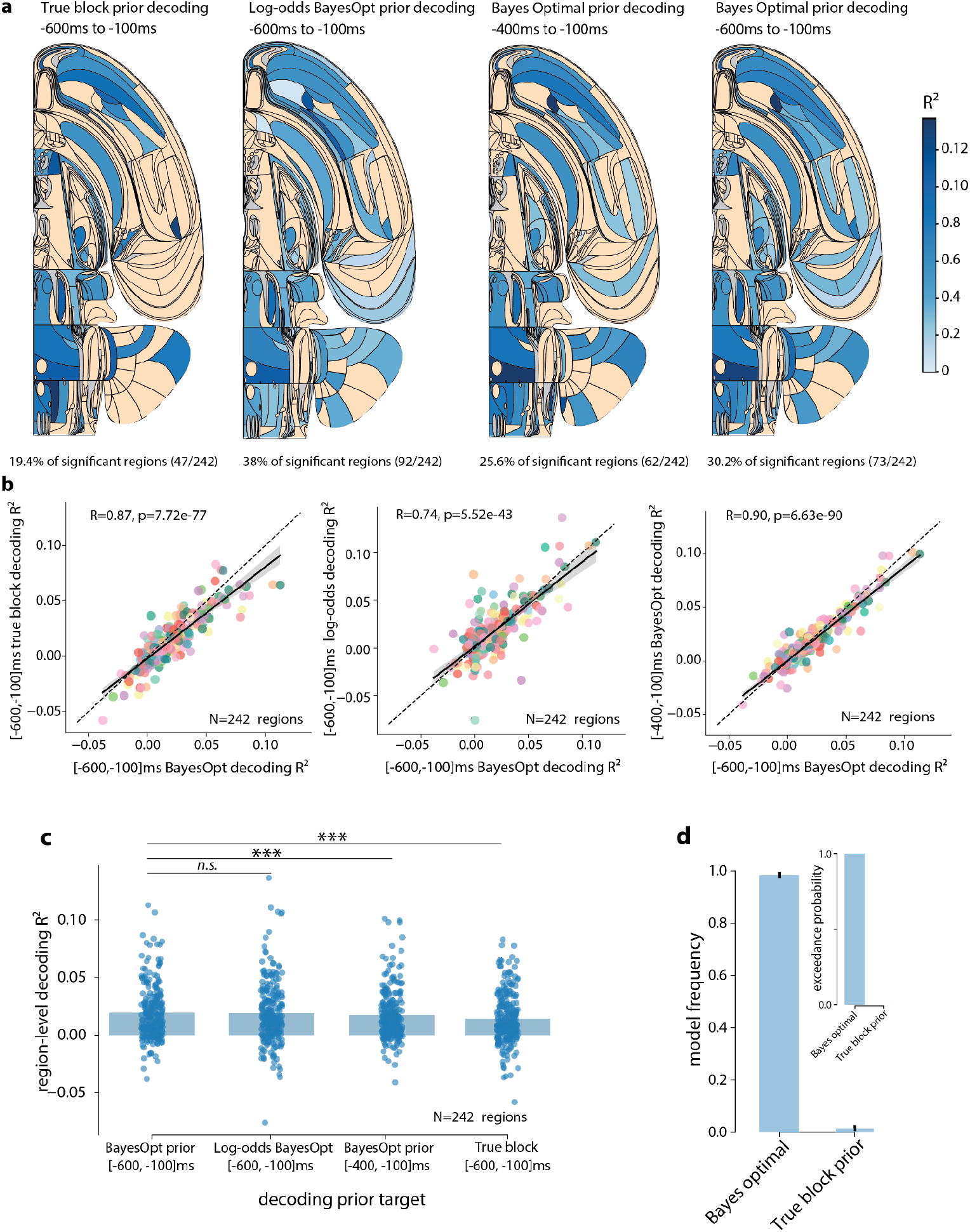
**a**. Swanson maps showing corrected decoding *R*^2^ values for various decoding priors. From left to right: True block prior, log odds ratio of the Bayes-optimal prior, Bayes-optimal prior on a narrower time window (−400ms to −100ms), and the Bayes-optimal prior from main Fig. 2b. A region is deemed significant if the Fisher combined *p*-value is lower than 0.05. **b**. Correlation analysis comparing Bayes optimal decoding from the extended window (shown in Fig. 2b) with the true block decoding (left panel), the log odds prior (middle panel), and the Bayes-optimal prior from the narrower window (right panel). In the three cases, we have a large correlation between corrected *R*^2^ **c**. Comparison of the corrected *R*^2^ across the four decodings, testing whether the points panel **b**. are over or below the diagonal (2-tailed signed-rank paired Wilcoxon test, *n*.*s*. not significant, *** p<0.001). **d**. Bayesian model comparison for 2 behavioral models, the Bayes optimal model, which infers a prior from past observations (see Methods and Supplementary Information), and a model that assumes the true block prior, which is not accessible to the mice. Our analysis shows that the Bayes optimal model more effectively explains the behavior, with an exceedance probability greater than 0.999.

**Figure S7:**
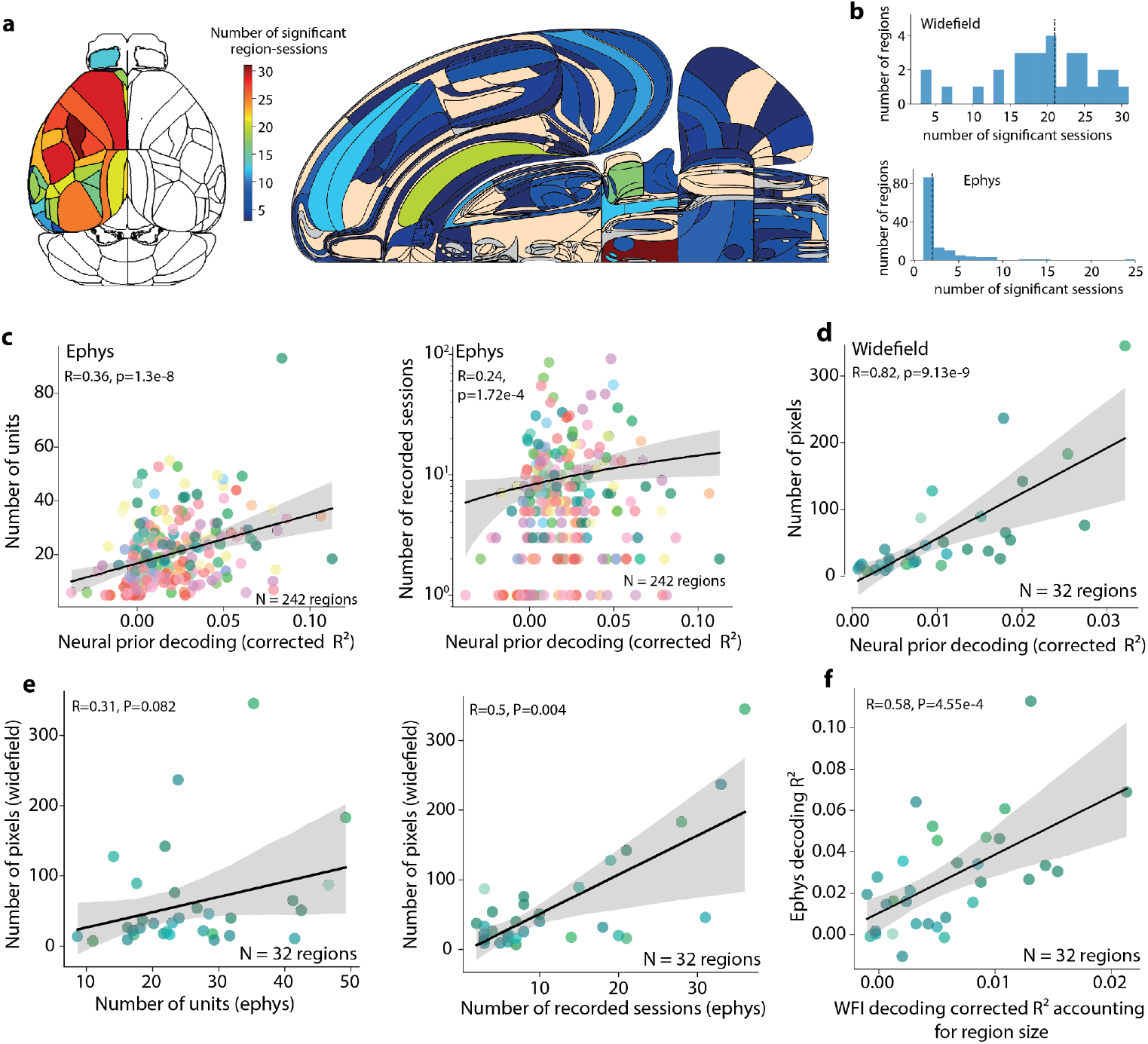
**a**. Number of significant sessions for each region represented on the dorsal map for WFI and the Swanson map for Ephys. **b**. Histograms representing the distribution of the number of significant sessions for each region in both WFI (top) and Ephys (bottom). Note that the number of significant sessions per region in Ephys is low, which prevents us from making robust claims at the regional level. **c**. Number of units (left) and number of recorded sessions (right) as a function of the decoded *R*^*2*^ for the Bayes-optimal prior in Ephys. **d**. Number of pixels as a function of the decoded *R*^*2*^ for the Bayes-optimal prior in WFI. **e**. Correlations between confounds across modalities. Left panel: Number of pixels in WFI as a function of the number of units in Ephys. Right: Number of pixels as a function of the number of recorded sessions in Ephys. **f**. Corrected *R*^*2*^ for Ephys as a function of the corrected *R*^*2*^ for WFI after correcting the WFI *R*^*2*^ data for region size. Correcting for the region size in WFI was performed by subtracting the size-predicted *R*^*2*^ (from panel **d**) from the WFI *R*^*2*^. Each dot corresponds to one region. All Ephys regions (significant and non-significant) were included in this analysis.

**Figure S8.**
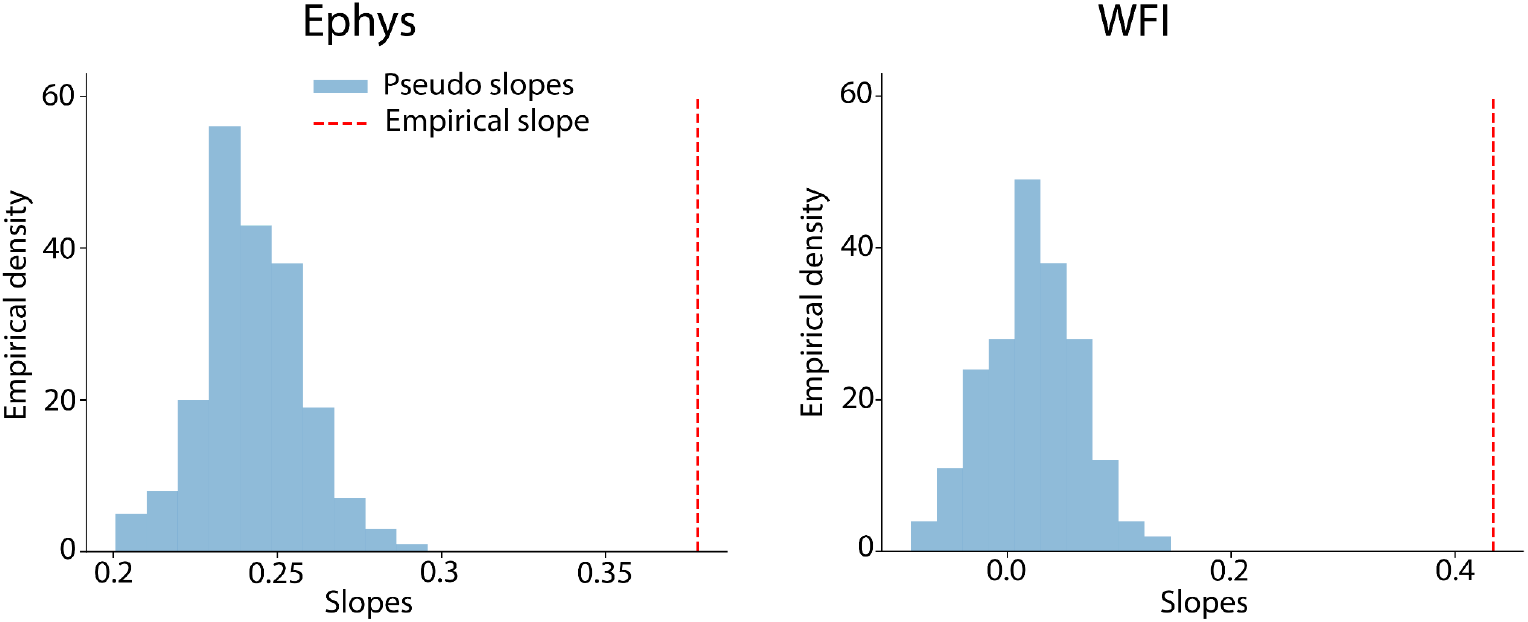
Null distribution of the slopes for the proportion of right choice vs decoded prior on zero contrast trials. Slopes were estimated using logistic regression to predict the choice (left or right) as a function of the decoded prior. The null distribution was calculated using 200 pseudosessions. For each pseudosession, pseudoactions were generated from an action kernel behavioral model that was fit to each real session (see Methods for more details). We then obtained pseudoslopes by predicting (with logistic regression) the pseudoactions as a function of the decoded prior. The null distribution was obtained by averaging the pseudoslopes across all sessions (we thus obtain 200 averaged pseudoslopes). The empirical average slope (red dashed lines, values corresponding to Fig. 2e) does not overlap with the null distribution obtained with pseudosessions (blue histogram). Therefore the correlations between the predicted prior and proportion of right choice can not be explained away by spurious temporal correlations or drift in the neural recordings.

**Figure S9.**
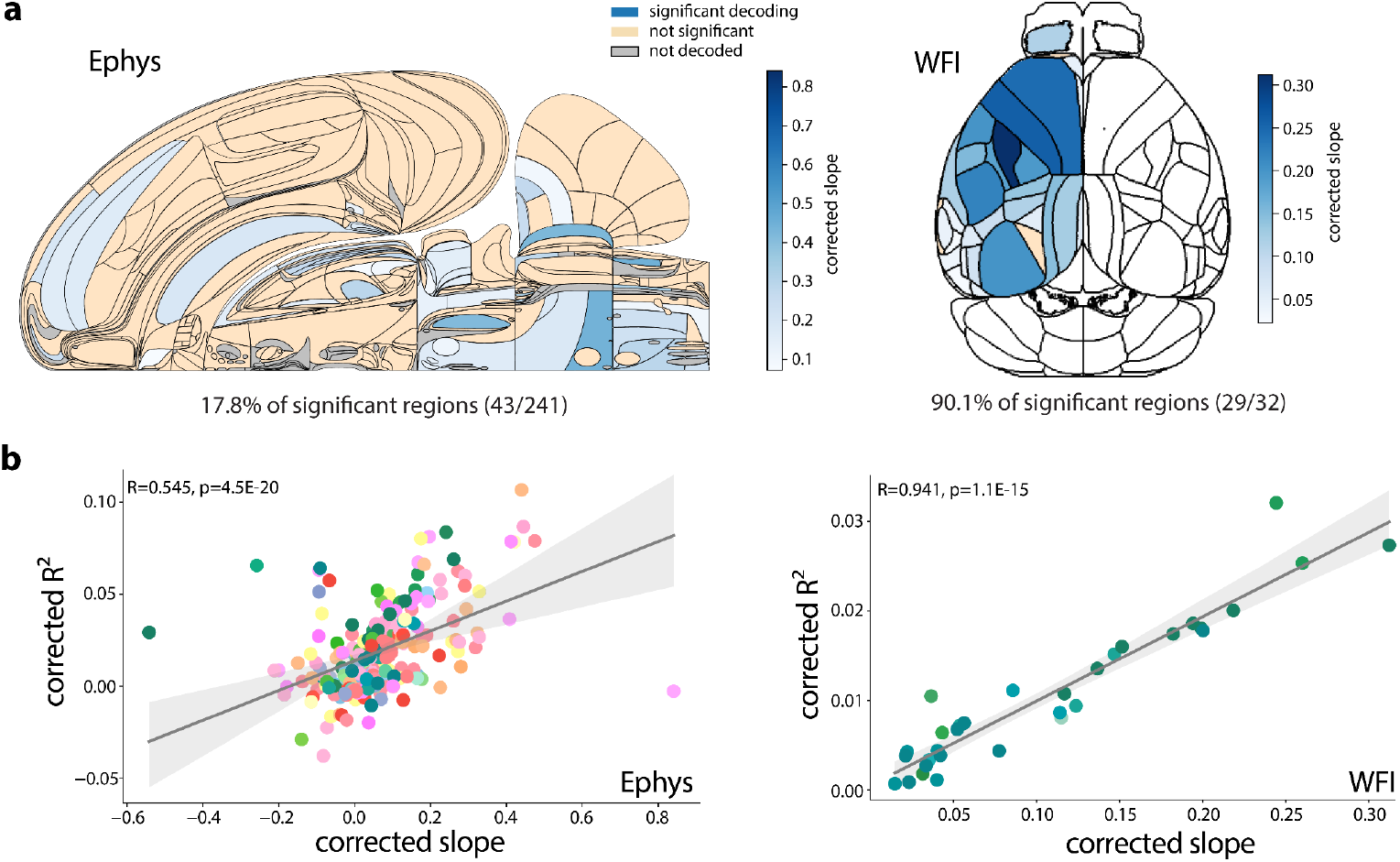
Proportion of right choices on zero contrast trials as a function of the decoded region-level Bayes-optimal prior. We decoded the Bayes-optimal prior for each region and computed the slope of this decoded prior as a function of the proportion of right choices (corrected using pseudo-sessions). This is the analysis presented in Fig 2e but at a region level. **a**. Region-level corrected slopes for Ephys and Widefield (significance is assessed when the region-level p-values < 0.05, using Fisher’s method for combining p-values). We observed that the slopes are significant in 17.8% of the regions in Ephys and 90.1% in Widefield, spanning every level of the hierarchy, including LGd, SCm, CP, MOs, and ACAd. It should be noted that the analysis for Ephys includes only 241 regions due to the exclusion of two sessions where the mouse made the same choice on every zero contrast trial. **b**. Correlation at the regional level between the decoded R^2^ values and the corrected slopes. We find a correlation in both modalities. These correlations prompt further investigation into whether they could be explained away by differences in how the Bayes optimal prior versus the action kernel model account for behavior across sessions. Specifically, sessions that more closely follow the action kernel model could potentially show lower corrected *R*^2^ and slopes, as these metrics are calculated using the Bayes optimal prior. In Ephys, we found no correlation between the log Bayes Factor (the difference in the marginal log likelihood between the action kernel and Bayes optimal models at the session level) and the corrected slopes (Spearman correlation: R=0.05, p=0.29, N=412 sessions), with the corrected slopes averaged across regions for each session. In widefield, a small correlation was detected (Spearman correlation: R=-0.34, p=0.014, N=51 sessions). However, even after adjusting for the log Bayes factor (by removing the linear prediction of the log Bayes factor from the corrected slope), the correlation between the corrected *R*^2^ and the adjusted corrected slope remained strong (Spearman correlation: R=0.935, p=4.7E-15, N=32 regions). This suggests that the type of behavioral strategy the mice used does not confound the correlation between the corrected *R*^2^ and the corrected slope.

**Figure S10.**
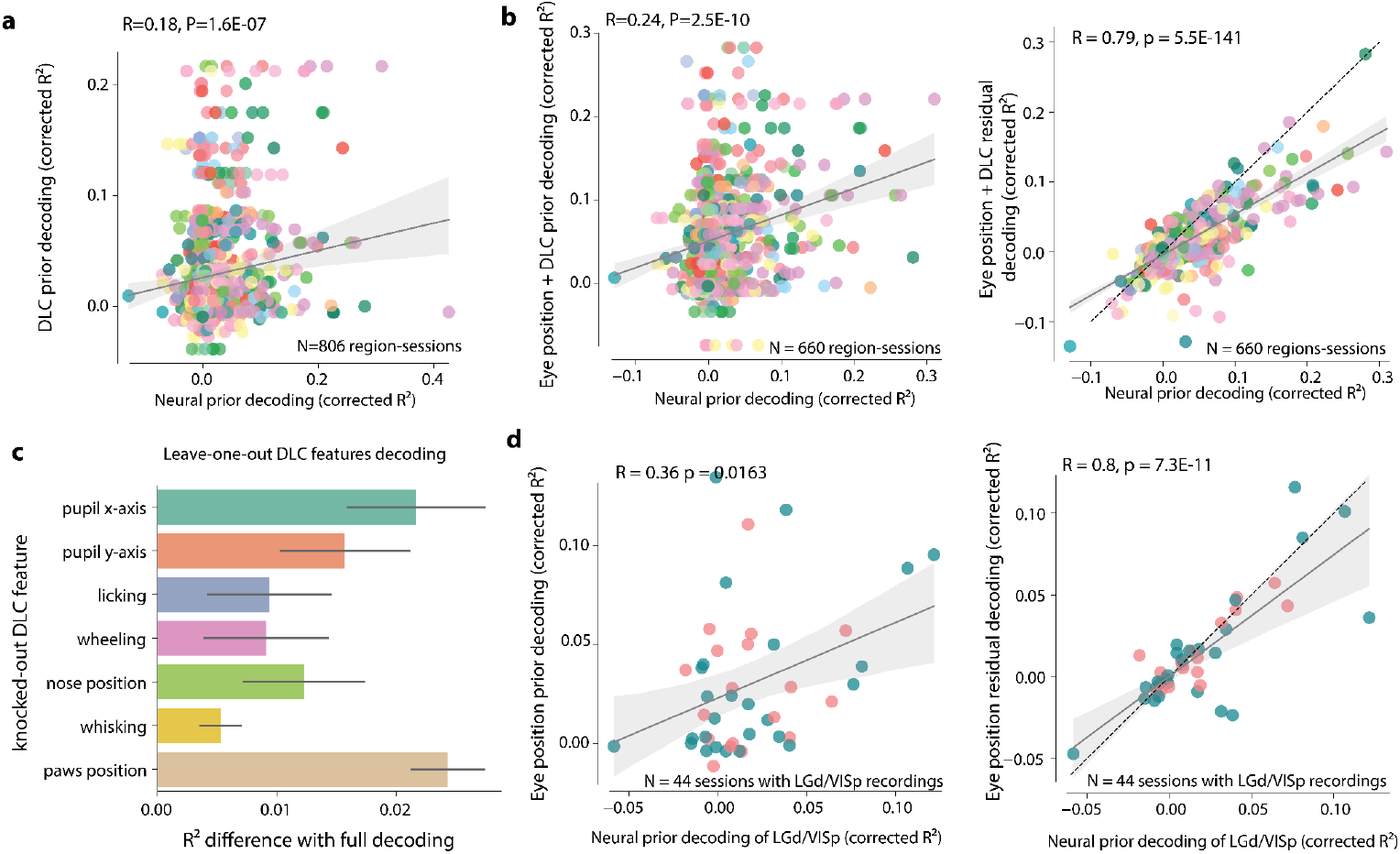
**a**. The decoding *R*^2^ for the Bayes-optimal prior from neural activity is significantly correlated with the decoding *R*^2^ for the Bayes-optimal prior from DLC features (Pearson correlation R=0.18, p=1.6E-7). **b**. Embodiment analysis accounting for both the DLC features and the eye position. Left: Decoding *R*^2^ for the Bayes-optimal prior from neural activity against decoding *R*^2^ for the Bayes-optimal prior from DLC features and eye position. The correlation between these two quantities is significant (Pearson correlation *R=0*.*24, p=2*.*5E-10*). Right: DLC + eye position residual decoding *R*^2^ against neural decoding *R*^2^. The residual decoding *R*^2^ values are obtained by first regressing the Bayes-optimal prior from DLC features and eye position, and then regressing the prior residual (Bayes-optimal prior minus Bayes-optimal prior estimated from DLC features and eye position) against neural activity. The neural decoding *R*^2^ corresponds to the *R*^2^ when decoding the Bayes-optimal prior from neural activity. The two quantities are strongly correlated (Pearson correlation *R=0*.*79, p=5*.*5E-141*), suggesting that the prior cannot be entirely attributed to a combination of both DLC features and eye position. **c**. Regressor elimination approach: for each feature, we remove it to measure the decrease in the decoding score compared to the full model (see Methods). The first feature to impact the model significantly when removed is the paw position. In this task, the paws are typically engaged to manipulate the wheel, which in turn adjusts the stimulus. It appears that the paws are positioned differently—likely on the wheel—depending on whether the prior suggests the next side will be left or right. The second key feature was the x-coordinate of the eye position, which aligns with the task setup where the stimulus is positioned along a horizontal plane, indicating that the mice tend to look in the direction suggested by the Bayes-optimal prior. **d**. Left: decoding *R*^2^ for the Bayes-optimal prior from neural activity in VISp and LGd against decoding *R*^2^ for the Bayes-optimal prior from eye position. The correlation between these two quantities is significant (Pearson correlation *R=0*.*36, p=0*.*0163*). Right: residual decoding *R*^2^ against neural decoding *R*^2^. The residual decoding *R*^2^ values are obtained by first regressing the Bayes-optimal prior against eye position and then regressing the prior residual (Bayes-optimal prior minus Bayes-optimal prior estimated from eye position) against neural activity in VISp and LGd (see Methods). The neural decoding *R*^2^ corresponds to the *R*^2^ when decoding the Bayes-optimal prior from neural activity. The two quantities are strongly correlated (Pearson correlation *R=0*.*8, p=7*.*3E-11*), suggesting that the prior signals in LGd and VISp are not solely due to the position of the eyes across blocks.

**Figure S11.**
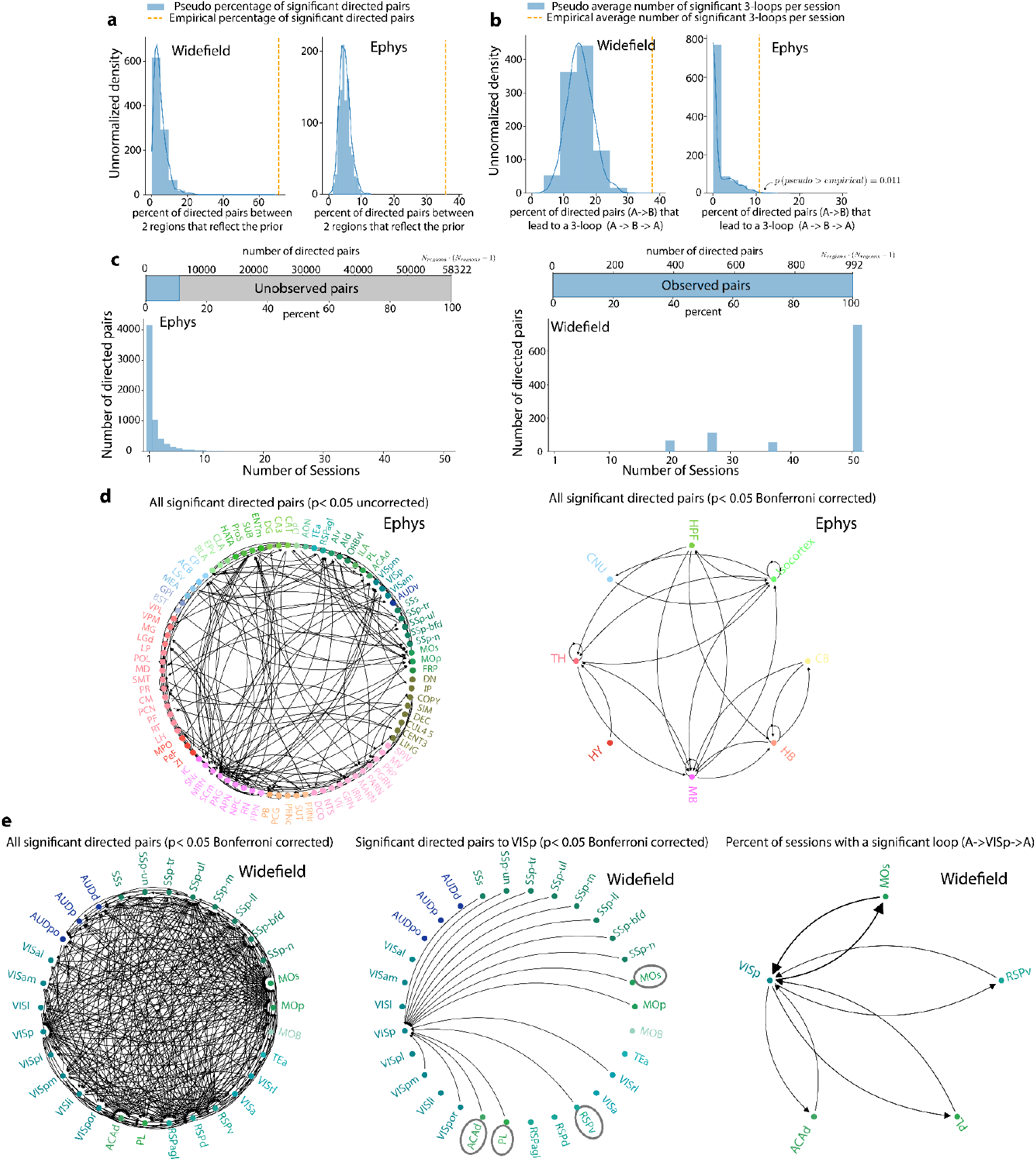
Granger causality analysis **a**. Average percentage of significant directed pairs between two regions that reflect the prior across sessions. When considering all pairs of regions encoding the prior significantly, and for which we had simultaneous recordings, we observed that information was significantly exchanged between 71% of these pairs in Widefield imaging and 36% in Ephys. Blue histograms: null distribution. **b**. Average percentage of significant directed pair (A->B) which is reciprocated within the same session by its counterpart (B->A), we found this to occur 38% of the time in Widefield and 11% in Ephys. Blue histograms: null distribution. (same as in main Fig. 2h). **c**. Histogram showing the number of sessions for each directed pair and barplot showing the percentage of observed directed pairs (directed pairs with at least one session) versus unobserved pairs. Right: In Ephys, with a total of 242 observed regions, the possible number of pairs amounts to 242 × 241 = 58,322. Of these, approximately 10% of the directed pairs had been recorded simultaneously, but the vast majority (75%) of these pairs appeared in two or fewer sessions, highlighting their scarcity. Left: Widefield provides a richer dataset, because, with 32 regions recorded simultaneously, we can analyze a total of 992 possible directed pairs (32 × 31), most of them available on all sessions. **d**. Left: Complete connectivity graph from Ephys (p < 0.05 uncorrected for multiple comparisons). When correcting for multiple comparisons, none of the links remains significant. This lack of significant findings post-correction is likely due to the sparse nature of the observations in Ephys (see panel c.). Right: Connectivity graph in Ephys across Cosmos regions (p < 0.05 Bonferroni corrected). *p*-values across directed pairs of regions are aggregated at the Cosmos level with Fisher’s method (see Methods, identical to main Fig. 2g left). **e**. Left: Complete connectivity graph from Widefield (p < 0.05 Bonferroni corrected). The graph is densely populated and consequently difficult to interpret. Middle: A partial connectivity graph from Widefield, highlighting significant directed pairs projecting to the Primary Visual Cortex (VISp), as shown in Figure 2g (right). We uncover feedback connections from higher-order areas such as the Motor Cortex (MOs), Ventral Retrosplenial Cortex (RSPv), Prelimbic Cortex (PL), and Anterior Cingulate Area Dorsal (ACAd) — these regions are marked with grey circles for emphasis — to the early sensory area, the Primary Visual Cortex (VISp). Left: Percentage of sessions exhibiting significant reciprocal connections (A->VISp->A) for sessions in which the Bayes optimal prior could be significantly decoded from both VISp and the previously identified higher-order regions (MOs, RSPv, PL and ACAd). The size of the arrow is proportional to the percentage. Our findings indicate the existence of reciprocal connections in these sessions: 33.3% between MOs and VISp, 16.7% between ACAd and VISp, 20% between PL and VISp, and 18.75% between RSPv and VISp.

**Figure S12:**
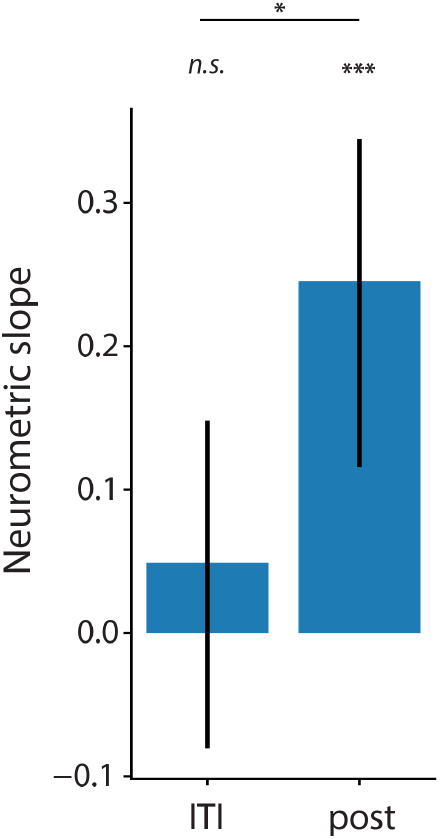
The average slope of the neurometric curves is significantly different from 0 during the post stimulus period (2-tailed signed-rank Wilcoxon test, t=9833, *p=1*.*2E-5, N=242 regions*) but not during the ITI (t=13547, *p=0*.*29, N=242 regions*). Also, neurometric slopes are significantly greater during the post-stimulus period than during the ITI (2-tailed signed-rank paired Wilcoxon test t=12306, *p=2*.*8E-2, N=242 regions*) (*** *p*<0.001, * *p*<0.05, *n*.*s*. not significant)

**Figure S13:**
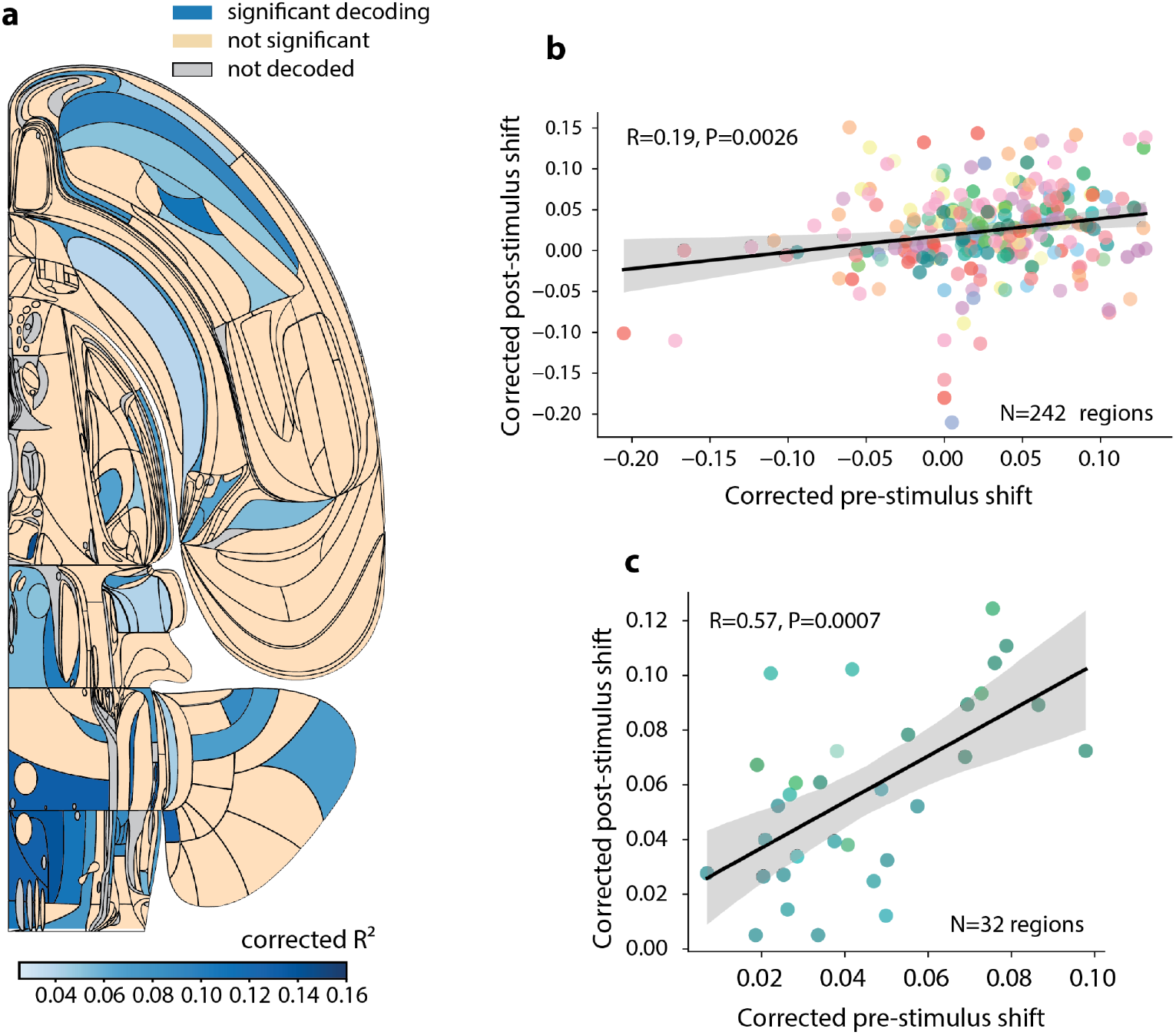
**a**. Swanson map of corrected neurometric post-stimulus shifts for Ephys data. **b**. The corrected post-stimulus shifts and corrected ITI shifts are significantly correlated in both Ephys (Spearman correlation *R=0*.*19, p=0*.*0026, N=242* regions) and WFI (Spearman correlation *R*=0.57, *p*=0.0007, N=32 regions).

**Figure S14:**
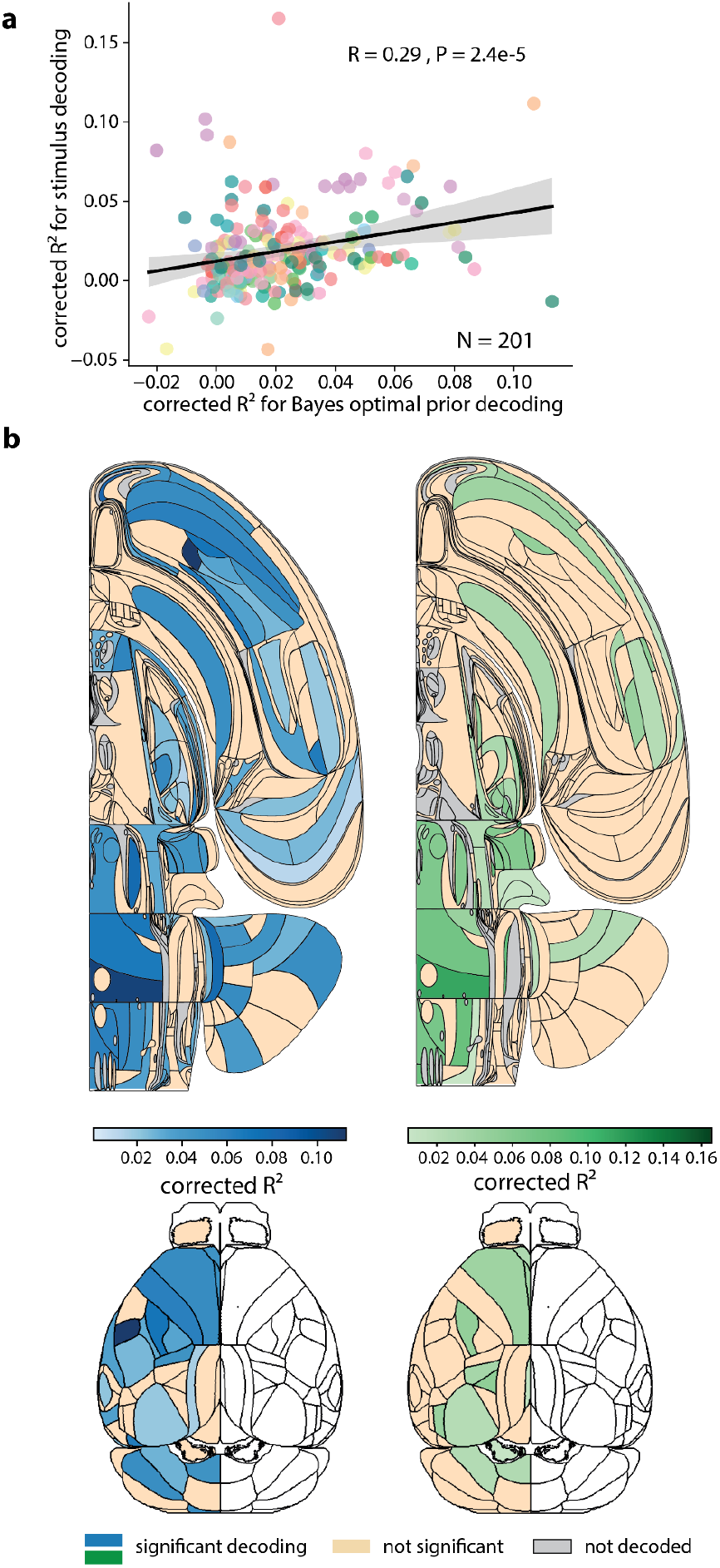
**a**. The neural decoding *R*^2^ for the stimulus side and the Bayes-optimal prior are significantly correlated across brain regions (Spearman correlation *R*=0.29, *p*=2.4e-5). **b**. Swanson maps and dorsal cortical views of brain regions encoding the Bayes-optimal prior (blue, left) and the stimulus side (green, right) significantly based on Ephys data.

**Figure S15.**
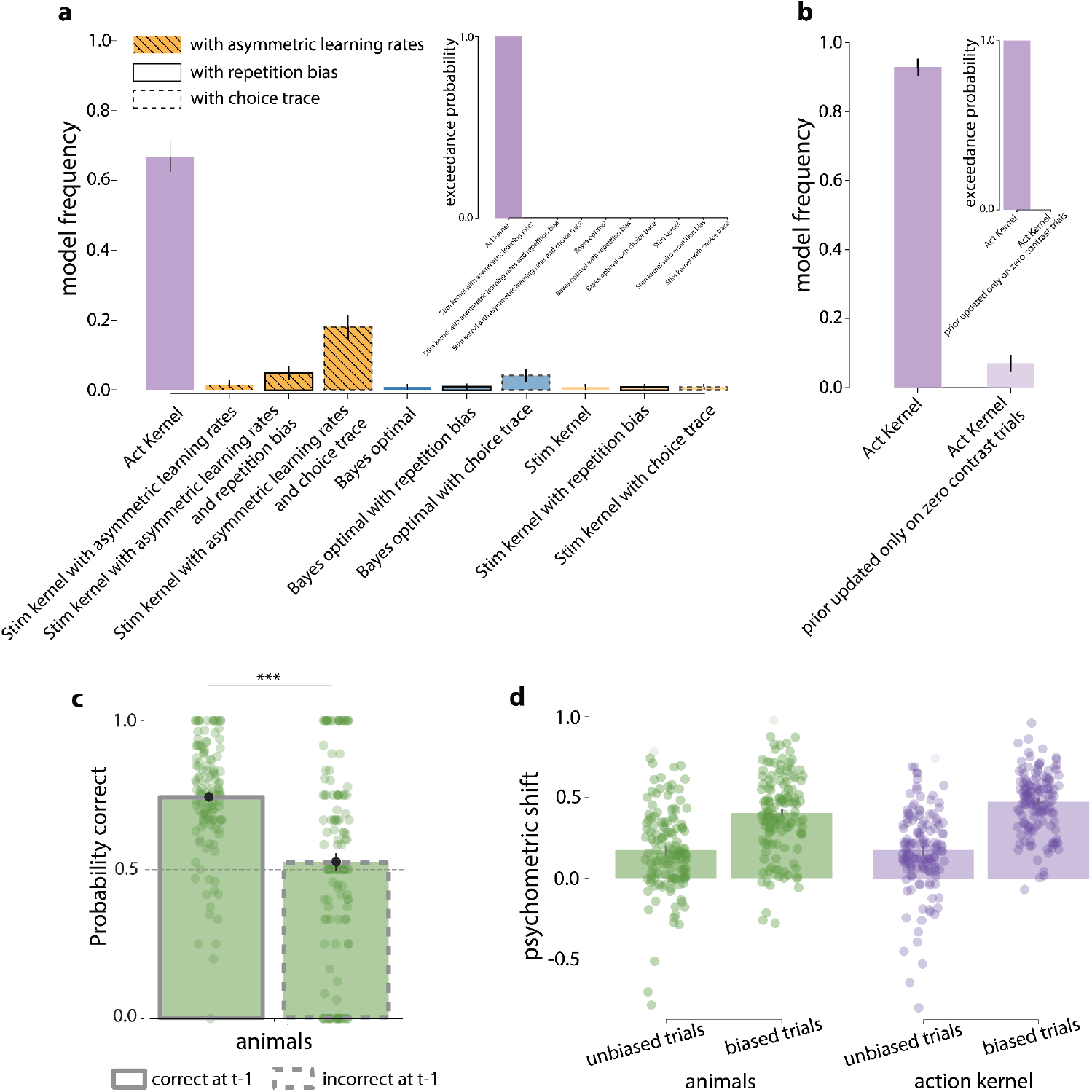
**a**. Bayesian model comparison for 11 behavioral models, considering the possibility of one step repetition bias (i.e. a tendency to repeat the previous choice), multi-step repetition bias (i.e. a tendency to follow an exponentially decaying average of past choices), and, for the stimulus kernel model, the presence of positivity and confirmation biases as asymmetric learning rates (accounting for the possibility to learn differently from positive versus negative rewards and from information that confirms versus contradicts existing beliefs, (Palminteri & Lebreton, 2022). See Methods for more details on the Bayes-optimal, action kernel and stimulus kernel models and Supplementary information for the formal equations of the repetition bias and asymmetrical learning rates. Model frequency (the posterior probability of the model given the subjects’ data, left panel) and exceedance probability (the probability that a model is more likely than any other models, right panel) are shown. The action kernel model offered the best account of the data even when including models with repetition, positivity and confirmation biases (p_exceedance_ > 0.999). **b**. Bayesian model comparison for two behavioral models, the action kernel and a variant that operates only during 0% contrast trials (by calculating an exponentially decaying average of chosen actions at 0% contrast trials). Our comparisons indicate that the action kernel, updating across all contrasts, more effectively explains behavior (exceedance probability > 0.999), suggesting that mice do not limit their subjective prior estimations to 0% contrast trials alone. **c**. Performance on zero contrast trials, distinguishing whether the preceding action was correct or incorrect and considering that the previous contrast was non zero. This analysis mirrors the main analysis in Fig. 4c but is specifically restricted to previous trials with non-zero contrast. When considering behavior within blocks, an agent using an action kernel prior should show a higher percentage of correct responses following a correct, block-consistent action compared to an incorrect one. This is because, on incorrect trials, the prior is updated with an action corresponding to the incorrect stimulus side. Even when limited to previous trials with non-zero contrast, there is a notable difference in the probability of making a correct decision following an incorrect vs. a correct choice (Wilcoxon paired test, t=11734, p=1.1E-15). This finding is confirmation that mice update their priors using information from all contrast levels, not solely zero contrast trials. **d**. Psychometric shift during both the first 90 trials (unbiased) and the other trials (biased) for animals and the action kernel. This shift is determined by analyzing two psychometric curves, one conditioned on the action kernel prior being above 0.5 (favoring the right side) and the other conditioned on the action kernel prior being less than 0.5 (favoring the left side). We fit psychometric functions to these curves (using the psychofit toolbox), and then calculate the psychometric shift as the vertical displacement of these curves at zero contrast. As predicted by the action kernel model, the analysis reveals a significant positive psychometric shift during the unbiased phase (first 90 trials). Furthermore, the shift in the behavioral data is less pronounced during the unbiased period compared to the biased period. This shift decrease during the unbiased phase can be explained by the subjective priors, which were closer to 0.5, mirroring the true block prior set at 0.5 for this period. Specifically, when distinguishing the trials that favor the right side (action kernel prior above 0.5) from those favoring the left side (action kernel prior below 0.5), the underlying action kernel priors remained close to 0.5 during the unbiased period. However, the presence of significant and comparable shifts between the animals and the action kernel model during the unbiased period indicates that mice exhibit a behavioral shift during the unbiased trials.

**Figure S16:**
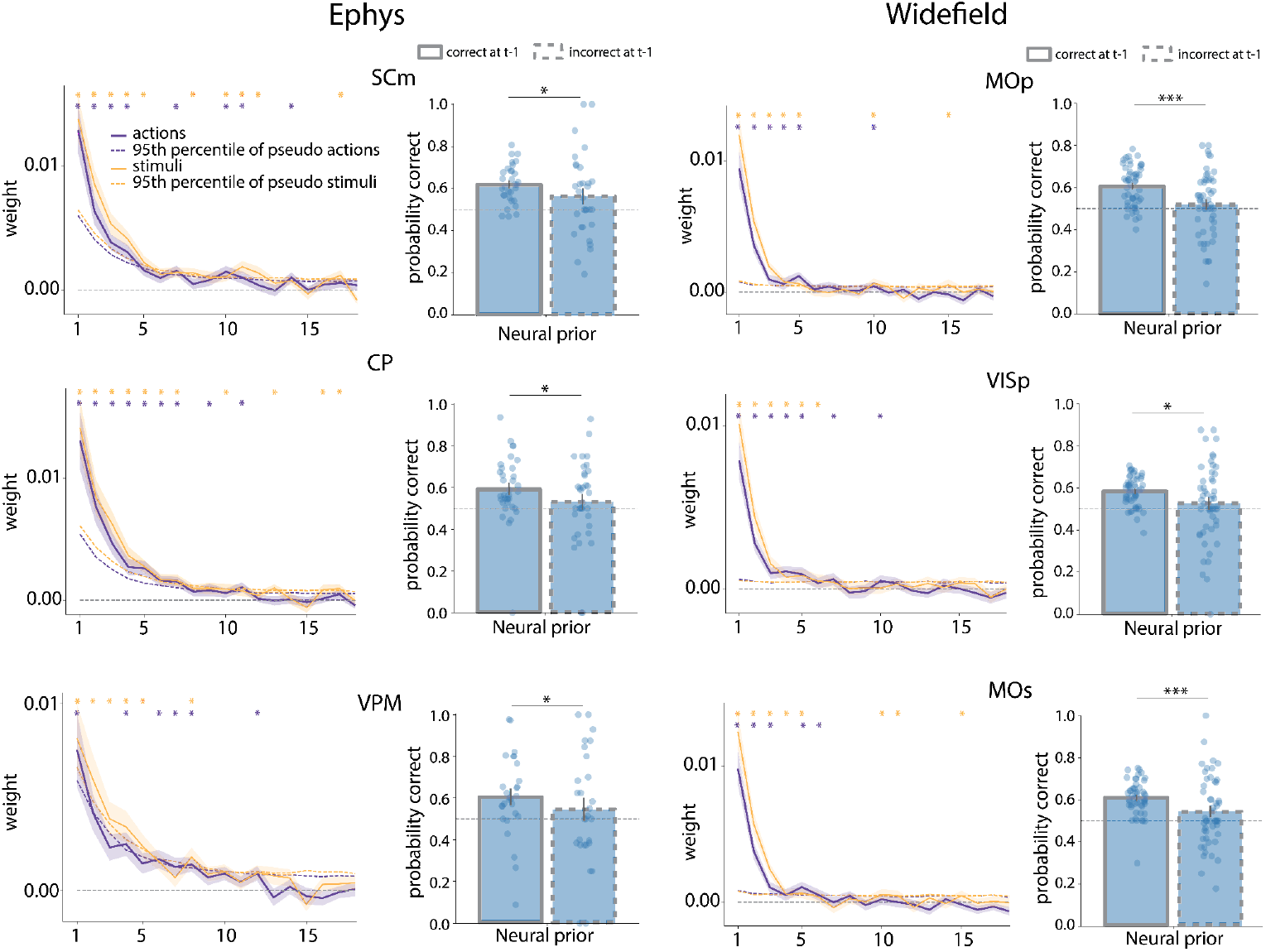
Same analysis as in Fig. 4c,e, but for three specific brain regions using Ephys (SCm, CP, VPM) or WFI data (right column, MOp, VISp, MOs) (* p<0.05, ** p<0.01, *** p<0.001). For the influence of past actions on the decoded Bayes-optimal prior, significance is assessed in the same way as in the main figure 4e (see Methods). For the asymmetry effect, the effect being observed on a brain-wide level, we performed a 1-tailed signed-rank Wilcoxon paired test for assessing significance on the region level.

**Figure S17.**
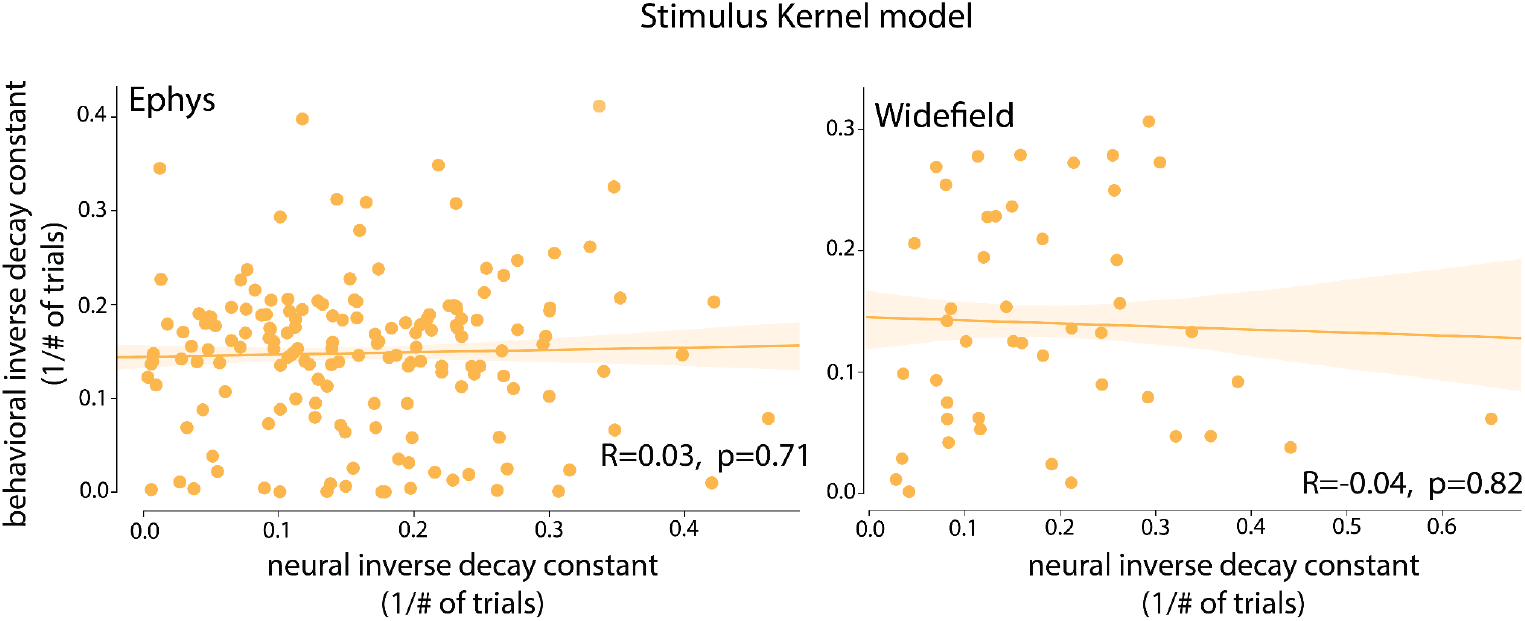
Behavioral inverse decay constants, obtained by fitting the stimulus kernel model to the behavior, as a function of the neural inverse decay constants, obtained by estimating the temporal dependency of the neural signals with respect to previous stimuli (see Methods). The neural and behavioral inverse decay constants are not significantly correlated for either Ephys (Pearson correlation *R*=0.03, *p*=0.71) or WFI (Pearson correlation *R*=-0.04, *p*=0.82)

**Figure S18.**
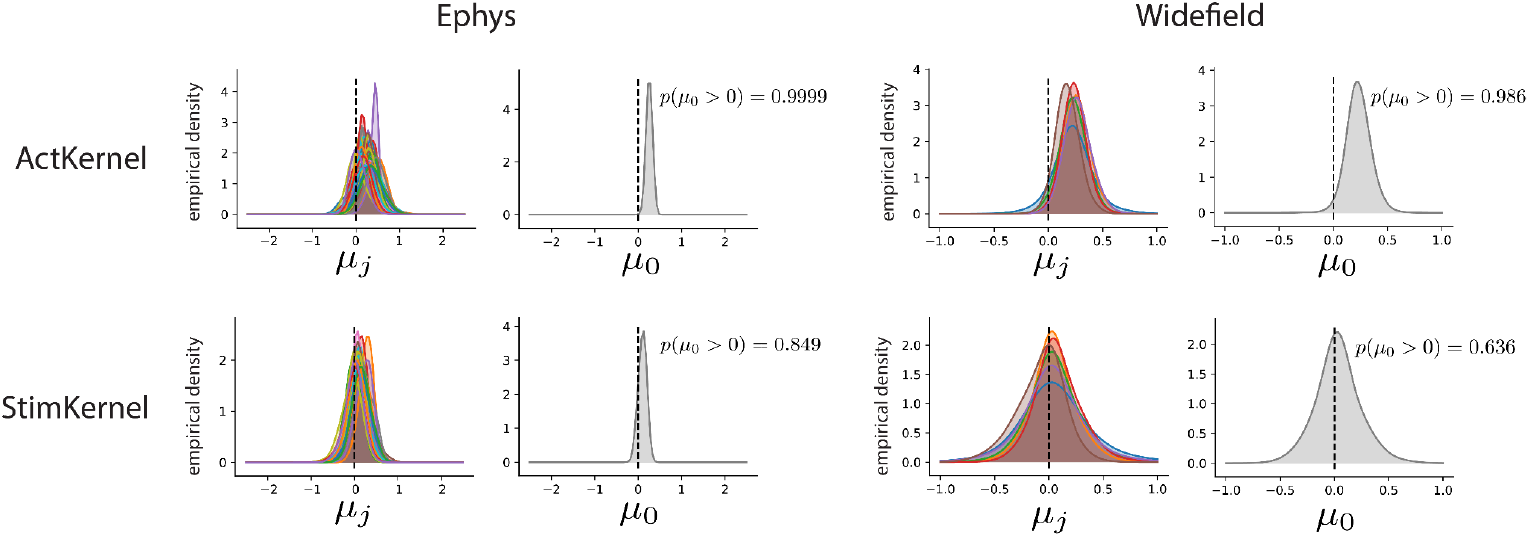
Hierarchical modeling of the neural and behavioral inverse decay constants (also referred to here as learning rates). The parameter μ_*j*_, defined for each mouse *j*, is the slope (the multiplicative coefficient) of the linear regression predicting the neural learning rate from the behavioral learning rate (on the sessions of mouse j). These parameters μ_*j*_ are sampled from a common population level prior with mean μ_0_.The parameter μ_0_, defined at the population level, characterizes an overall relationship between neural and behavioral learning rates. We found that the relationship between neural and behavioral learning rates is significantly positive for the action kernel model (top row), both in electrophysiology (left column) and in widefield imaging (right column), which is not the case for the stimulus Kernel model (bottom row). Furthermore, when testing the difference in means of the population level parameter μ_0_ between action and stimulus kernels, we found that it was significantly greater for the action kernel, both in Ephys and in WFI. Significance was assessed by estimating the means of the μ_0_ distributions for the action and stimulus kernels with the BEST Bayesian test (Kruschke, 2013). In both Ephys and WFI, we found that 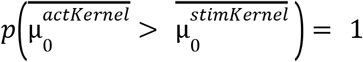 with 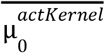 and 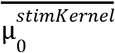 the means of the μ_0_ distributions for the action and stimulus kernels, respectively. Regarding the effect sizes, with the same BEST procedure, we find an effect size of 2.53 in Ephys and 1.96 in widefield (effect sizes greater than 1.3 are commonly considered to be very large (Sullivan & Feinn, 2012)). See Supplementary Information for the full specification of the hierarchical generative model.

**Figure S19.**
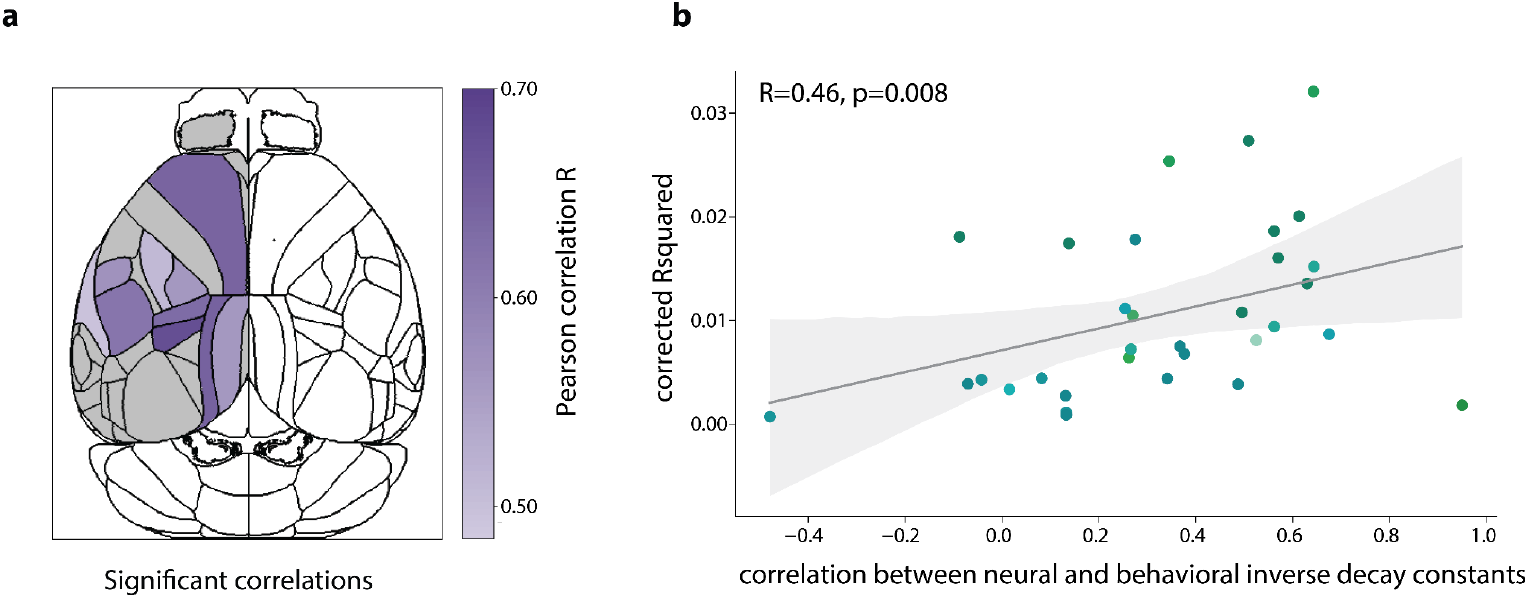
Correlation between behavioral and neural inverse decay constants across sessions at the region level. A decay constant is estimated for each pixel (as in figure 4f, refer to Methods), but now, averages are taken across pixels for each session and specific region. In the analysis figure 4f, session-level learning rates were obtained by averaging across all pixels, regardless of region identity. **a**. Regions with a significant correlation between behavioral and neural inverse decay constants. As expected, only positive correlations emerge as significant. **b**. Correlation between behavioral and neural inverse decay constants is correlated with the prior decoding corrected *R*^*2*^ from the same regions. Remarkably, these two quantities were found to be also correlated (R=0.46, p=0.008). In other words, regions in which the prior decoding *R*^*2*^ is large are also regions which best reflect the behavioral decay constant, i.e., these are the regions that are best correlated with the animals’ cognitive strategies as assessed by the lengths of the action kernels. We did not repeat this analysis with the electrophysiology recordings because we only have a very limited number of significant sessions per region (1-2 for most regions, as opposed to around 20 sessions per region for the WFI data - see Fig. S7a,b)

## References

Ackley, D. H., Hinton, G. E., & Sejnowski, T. J. (1985). A Learning Algorithm for Boltzmann Machines*. Cognitive Science, 9(1), 147–169. 10.1207/s15516709cog0901_7

Andrieu, C., & Thoms, J. (2008). A tutorial on adaptive MCMC. Statistics and Computing, 18(4), 343–373. 10.1007/s11222-008-9110-y

Ashwood, Z. C., Roy, N. A., Stone, I. R., The International Brain Laboratory, Urai, A. E., Churchland, A. K., Pouget, A., & Pillow, J. W. (2022). Mice alternate between discrete strategies during perceptual decision-making. Nature Neuroscience, 25(2), 201–212. 10.1038/s41593-021-01007-z

Bell, A. H., Summerfield, C., Morin, E. L., Malecek, N. J., & Ungerleider, L. G. (2016). Encoding of Stimulus Probability in Macaque Inferior Temporal Cortex. Current Biology, 26(17), 2280–2290. 10.1016/j.cub.2016.07.007

Berkes, P., Orbán, G., Lengyel, M., & Fiser, J. (2011). Spontaneous Cortical Activity Reveals Hallmarks of an Optimal Internal Model of the Environment. Science, 331(6013), 83–87. 10.1126/science.1195870

Biderman, D., Whiteway, M. R., Hurwitz, C., Greenspan, N., Lee, R. S., Vishnubhotla, A., Warren, R., Pedraja, F., Noone, D., Schartner, M., Huntenburg, J. M., Khanal, A., Meijer, G. T., Noel, J.-P., Pan-Vazquez, A., Socha, K. Z., Urai, A. E., The International Brain Laboratory, Cunningham, J. P., … Paninski, L. (2023). Lightning Pose: Improved animal pose estimation via semi-supervised learning, Bayesian ensembling, and cloud-native open-source tools [Preprint]. Neuroscience. 10.1101/2023.04.28.538703

Birman, D., Bonacchi, N., Buchanan, K., Chapuis, G., Meijer, G., Paninski, L., Schartner, M., Svoboda, K., Wells, M., Whiteway, M. R., & Winter, O. (n.d.). Video hardware and software for the International Brain Laboratory.

Bolkan, S. S., Stone, I. R., Pinto, L., Ashwood, Z. C., Iravedra Garcia, J. M., Herman, A. L., Singh, P., Bandi, A., Cox, J., Zimmerman, C. A., Cho, J. R., Engelhard, B., Pillow, J. W., & Witten, I. B. (2022). Opponent control of behavior by dorsomedial striatal pathways depends on task demands and internal state. Nature Neuroscience, 25(3), 345–357. 10.1038/s41593-022-01021-9

Bondy, A. G., Haefner, R. M., & Cumming, B. G. (2018). Feedback determines the structure of correlated variability in primary visual cortex. Nature Neuroscience, 21(4), 598–606. 10.1038/s41593-018-0089-1

Brooks, S. P., & Gelman, A. (1998). General Methods for Monitoring Convergence of Iterative Simulations. Journal of Computational and Graphical Statistics, 7(4), 434–455. 10.1080/10618600.1998.10474787

Busse, L., Ayaz, A., Dhruv, N. T., Katzner, S., Saleem, A. B., Schölvinck, M. L., Zaharia, A. D., & Carandini, M. (2011). The Detection of Visual Contrast in the Behaving Mouse. The Journal of Neuroscience, 31(31), 11351–11361. 10.1523/JNEUROSCI.6689-10.2011

Dabney, W., Kurth-Nelson, Z., Uchida, N., Starkweather, C. K., Hassabis, D., Munos, R., & Botvinick, M. (2020). A distributional code for value in dopamine-based reinforcement learning. Nature, 577(7792), 671–675. 10.1038/s41586-019-1924-6

Echeveste, R., Aitchison, L., Hennequin, G., & Lengyel, M. (2020). Cortical-like dynamics in recurrent circuits optimized for sampling-based probabilistic inference. Nature Neuroscience, 23(9), 1138–1149. 10.1038/s41593-020-0671-1

Elber-Dorozko, L., & Loewenstein, Y. (2018). Striatal action-value neurons reconsidered. eLife, 7, e34248. 10.7554/eLife.34248

Ernst, M. O., & Banks, M. S. (2002). Humans integrate visual and haptic information in a statistically optimal fashion. Nature, 415(6870), 429–433. 10.1038/415429a

Forstmann, B. U. (2010). The neural substrate of prior information in perceptual decision making: A model-based analysis. Frontiers in Human Neuroscience, 4. 10.3389/fnhum.2010.00040

Ganguli, D., & Simoncelli, E. P. (2014). Efficient Sensory Encoding and Bayesian Inference with Heterogeneous Neural Populations. Neural Computation, 26(10), 2103–2134. 10.1162/NECO_a_00638

Haefner, R. M., Berkes, P., & Fiser, J. (2016). Perceptual Decision-Making as Probabilistic Inference by Neural Sampling. Neuron, 90(3), 649–660. 10.1016/j.neuron.2016.03.020

Han, S., & Helmchen, F. (2023). Behavior-relevant top-down cross-modal predictions in mouse neocortex [Preprint]. Neuroscience. 10.1101/2023.04.03.535389

Hanks, T. D., Mazurek, M. E., Kiani, R., Hopp, E., & Shadlen, M. N. (2011). Elapsed Decision Time Affects the Weighting of Prior Probability in a Perceptual Decision Task. Journal of Neuroscience, 31(17), 6339–6352. 10.1523/JNEUROSCI.5613-10.2011

Hansen, K. A., Hillenbrand, S. F., & Ungerleider, L. G. (2012). Human Brain Activity Predicts Individual Differences in Prior Knowledge Use during Decisions. Journal of Cognitive Neuroscience, 24(6), 1462–1475. 10.1162/jocn_a_00224

Harris, K. D. (2020). Nonsense correlations in neuroscience [Preprint]. Neuroscience. 10.1101/2020.11.29.402719

Hoyer, P. O., & Hyvärinen, P. A. (2003). Interpreting neural response variability as Monte Carlo sampling of the posterior. In Advances in Neural Information Processing Systems (Vol. 15, pp. 293–300).

International Brain Lab, Benson, B., Benson, J., Birman, D., Bonacchi, N., Carandini, M., Catarino, J. A., Chapuis, G. A., Churchland, A. K., Dan, Y., Dayan, P., DeWitt, E. E., Engel, T. A., Fabbri, M., Faulkner, M., Fiete, I. R., Findling, C., Freitas-Silva, L., Gercek, B., … Witten, I. B. (2023). A Brain-Wide Map of Neural Activity during Complex Behaviour [Preprint]. Neuroscience. 10.1101/2023.07.04.547681

Ishizu, K., Nishimoto, S., & Funamizu, A. (2023). Localized and global computation for integrating prior value and sensory evidence in the mouse cerebral cortex [Preprint]. Neuroscience. 10.1101/2023.06.06.543645

Jacobs, R. A. (1999). Optimal integration of texture and motion cues to depth. Vision Research, 39(21), 3621–3629. 10.1016/S0042-6989(99)00088-7

Jardri, R., Duverne, S., Litvinova, A. S., & Denève, S. (2017). Experimental evidence for circular inference in schizophrenia. Nature Communications, 8(1), 14218. 10.1038/ncomms14218

Knill, D. C., & Pouget, A. (2004). The Bayesian brain: The role of uncertainty in neural coding and computation. Trends in Neurosciences, 27(12), 712–719. 10.1016/j.tins.2004.10.007

Kok, P., Jehee, J. F. M., & de Lange, F. P. (2012). Less Is More: Expectation Sharpens Representations in the Primary Visual Cortex. Neuron, 75(2), 265–270. 10.1016/j.neuron.2012.04.034

Krasniak, C. (2022). Mesoscale imaging, inactivation, and collaboration in a standardized visual decision-making task. Cold Spring Harbor Laboratory. http://repository.cshl.edu/id/eprint/40616/

Kruschke, J. K. (2013). Bayesian estimation supersedes the t test. Journal of Experimental Psychology: General, 142(2), 573–603. 10.1037/a0029146

Laboratory, International Brain. (2022). Video hardware and software for the International Brain Laboratory. 7030489 Bytes. 10.6084/M9.FIGSHARE.19694452.V1

Laboratory, International Brain. (2023). Data release—Brainwide map—Q4 2022. 12507703 Bytes. 10.6084/M9.FIGSHARE.21400815.V6

Lak, A., Okun, M., Moss, M. M., Gurnani, H., Farrell, K., Wells, M. J., Reddy, C. B., Kepecs, A., Harris, K. D., & Carandini, M. (2020). Dopaminergic and Prefrontal Basis of Learning from Sensory Confidence and Reward Value. Neuron, 105(4), 700–711.e6. 10.1016/j.neuron.2019.11.018

Lange, R. D., & Haefner, R. M. (2022). Task-induced neural covariability as a signature of approximate Bayesian learning and inference. PLOS Computational Biology, 18(3), e1009557. 10.1371/journal.pcbi.1009557

Lau, B., & Glimcher, P. W. (2005). DYNAMIC RESPONSE-BY-RESPONSE MODELS OF MATCHING BEHAVIOR IN RHESUS MONKEYS. Journal of the Experimental Analysis of Behavior, 84(3), 555–579. 10.1901/jeab.2005.110-04

Lima, V., Dellajustina, F. J., Shimoura, R. O., Girardi-Schappo, M., Kamiji, N. L., Pena, R. F. O., & Roque, A. C. (2020). Granger causality in the frequency domain: Derivation and applications. Revista Brasileira de Ensino de Física, 42, e20200007. 10.1590/1806-9126-rbef-2020-0007

Ma, W. J., Beck, J. M., Latham, P. E., & Pouget, A. (2006). Bayesian inference with probabilistic population codes. Nature Neuroscience, 9(11), 1432–1438. 10.1038/nn1790

Mamassian, P., Knill, D. C., & Kersten, D. (1998). The perception of cast shadows. Trends in Cognitive Sciences, 2(8), 288–295. 10.1016/S1364-6613(98)01204-2

Mathis, A., Mamidanna, P., Cury, K. M., Abe, T., Murthy, V. N., Mathis, M. W., & Bethge, M. (2018). DeepLabCut: Markerless pose estimation of user-defined body parts with deep learning. Nature Neuroscience, 21(9), 1281–1289. 10.1038/s41593-018-0209-y

Mayrhofer, J. M., El-Boustani, S., Foustoukos, G., Auffret, M., Tamura, K., & Petersen, C. C. H. (2019). Distinct Contributions of Whisker Sensory Cortex and Tongue-Jaw Motor Cortex in a Goal-Directed Sensorimotor Transformation. Neuron, 103(6), 1034–1043.e5. 10.1016/j.neuron.2019.07.008

Mendonça, A. G., Drugowitsch, J., Vicente, M. I., DeWitt, E. E. J., Pouget, A., & Mainen, Z. F. (2020). The impact of learning on perceptual decisions and its implication for speed-accuracy tradeoffs. Nature Communications, 11(1), 2757. 10.1038/s41467-020-16196-7

Mulder, M. J., Wagenmakers, E.-J., Ratcliff, R., Boekel, W., & Forstmann, B. U. (2012). Bias in the Brain: A Diffusion Model Analysis of Prior Probability and Potential Payoff. Journal of Neuroscience, 32(7), 2335–2343. 10.1523/JNEUROSCI.4156-11.2012

Niv, Y. (2019). Learning task-state representations. Nature Neuroscience, 22(10), 1544–1553. 10.1038/s41593-019-0470-8

Nogueira, R., Abolafia, J. M., Drugowitsch, J., Balaguer-Ballester, E., Sanchez-Vives, M. V., & Moreno-Bote, R. (2017). Lateral orbitofrontal cortex anticipates choices and integrates prior with current information. Nature Communications, 8(1), 14823. 10.1038/ncomms14823

Norton, E. H., Acerbi, L., Ma, W. J., & Landy, M. S. (2019). Human online adaptation to changes in prior probability. PLOS Computational Biology, 15(7), e1006681. 10.1371/journal.pcbi.1006681

Palminteri, S., & Lebreton, M. (2022). The computational roots of positivity and confirmation biases in reinforcement learning. Trends in Cognitive Sciences, 26(7), 607–621. 10.1016/j.tics.2022.04.005

Park, J., Kim, S., Kim, H. R., & Lee, J. (2022). Prior expectation enhances sensorimotor behavior by modulating population tuning and subspace activity in the sensory cortex [Preprint]. Neuroscience. 10.1101/2022.12.04.516847

Pedregosa, F., Varoquaux, G, Gramfort, A., Thirion, B., Grisel, O., Blondel, M., Prettenhofer, P., Weiss, R., Dubourg, V., Vanderplas, J., Passos, A., Coupaneau, D., Brucher, M., Perrot, M., & Duchesnay, E. (2011). Scikit-learn: Machine Learning in Python. 12, 2825–2830.

Platt, M. L., & Glimcher, P. W. (1999). Neural correlates of decision variables in parietal cortex. Nature, 400(6741), 233–238. 10.1038/22268

Rao, V., DeAngelis, G. C., & Snyder, L. H. (2012). Neural Correlates of Prior Expectations of Motion in the Lateral Intraparietal and Middle Temporal Areas. Journal of Neuroscience, 32(29), 10063–10074. 10.1523/JNEUROSCI.5948-11.2012

Sahani, M., & Dayan, P. (2003). Doubly Distributional Population Codes: Simultaneous Representation of Uncertainty and Multiplicity. Neural Computation, 15(10), 2255–2279. 10.1162/089976603322362356

Schaeffer, R., Khona, M., Meshulam, L., International Brain Laboratory, & Fiete, I. R. (2020). Reverse-engineering Recurrent Neural Network solutions to a hierarchical inference task for mice [Preprint]. Neuroscience. 10.1101/2020.06.09.142745

Scott, S. L. (2002). Bayesian Methods for Hidden Markov Models: Recursive Computing in the 21st Century. Journal of the American Statistical Association, 97(457), 337–351. 10.1198/016214502753479464

Soltani, A., & Wang, X.-J. (2010). Synaptic computation underlying probabilistic inference. Nature Neuroscience, 13(1), 112–119. 10.1038/nn.2450

Son, S., Moon, J., Kim, Y.-J., Kang, M.-S., & Lee, J. (2023). Frontal-to-visual information flow explains predictive motion tracking. NeuroImage, 269, 119914. 10.1016/j.neuroimage.2023.119914

Stephan, K. E., Penny, W. D., Daunizeau, J., Moran, R. J., & Friston, K. J. (2009). Bayesian model selection for group studies. NeuroImage, 46(4), 1004–1017. 10.1016/j.neuroimage.2009.03.025

Sugawara, M., & Katahira, K. (2021). Dissociation between asymmetric value updating and perseverance in human reinforcement learning. Scientific Reports, 11(1), 3574. 10.1038/s41598-020-80593-7

Sullivan, G. M., & Feinn, R. (2012). Using Effect Size—Or Why the P Value Is Not Enough. Journal of Graduate Medical Education, 4(3), 279–282. 10.4300/JGME-D-12-00156.1

The International Brain Laboratory, Aguillon-Rodriguez, V., Angelaki, D., Bayer, H., Bonacchi, N., Carandini, M., Cazettes, F., Chapuis, G., Churchland, A. K., Dan, Y., Dewitt, E., Faulkner, M., Forrest, H., Haetzel, L., Häusser, M., Hofer, S. B., Hu, F., Khanal, A., Krasniak, C., … Zador, A. M. (2021). Standardized and reproducible measurement of decision-making in mice. eLife, 10, e63711. 10.7554/eLife.63711

Urai, A. E., De Gee, J. W., Tsetsos, K., & Donner, T. H. (2019). Choice history biases subsequent evidence accumulation. eLife, 8, e46331. 10.7554/eLife.46331

Walker, E. Y., Cotton, R. J., Ma, W. J., & Tolias, A. S. (2020). A neural basis of probabilistic computation in visual cortex. Nature Neuroscience, 23(1), 122–129. 10.1038/s41593-019-0554-5

Wang, Q., Ding, S.-L., Li, Y., Royall, J., Feng, D., Lesnar, P., Graddis, N., Naeemi, M., Facer, B., Ho, A., Dolbeare, T., Blanchard, B., Dee, N., Wakeman, W., Hirokawa, K. E., Szafer, A., Sunkin, S. M., Oh, S. W., Bernard, A., … Ng, L. (2020). The Allen Mouse Brain Common Coordinate Framework: A 3D Reference Atlas. Cell, 181(4), 936–953.e20. 10.1016/j.cell.2020.04.007

Weiss, Y., Simoncelli, E. P., & Adelson, E. H. (2002). Motion illusions as optimal percepts. Nature Neuroscience, 5(6), 598–604. 10.1038/nn0602-858

Zemel, R. S., Dayan, P., & Pouget, A. (1998). Probabilistic Interpretation of Population Codes. Neural Computation, 10(2), 403–430. 10.1162/089976698300017818

