## Supplementary Information for "Brain-wide representations of prior information in mouse decision-making"

November 5, 2024

### 1 Behavioural models

The behavioural models section is divided into four subsections:

**Prior modelling:** describes the estimation of the prior belief as to where the stimulus is going to appear on the next trial.

**Evidence model:** describes how the evidence presented on the screen (the stimulus, in non-zero-contrast trials), is processed by the sensory areas

**Decision policy:** describes how the prior and the evidence are combined to lead to a choice

**Full likelihood:** makes explicit the full likelihood which is used to fit the behavioural model to the mouse's choices.

#### 1.1 Prior modelling

Here, we describe the three different models of the construction of the prior belief as to where the stimulus will appear on the next trial (in the mouse's internal model).

##### 1.1.1 Bayes Optimal: prior from the generative Bayesian model of the task

Let $l_t$ be the current length of the current block. Let $b_t$ be the current block label ($b_t = 0$ indicates unbiased, $b_t = -1$ indicates left-biased and $b_t = 1$ indicates right-biased). Let $s_t$ be the stimulus side ($s_t = -1$ indicates stimulus on left side, $s_t = 1$ indicates stimulus on right side).

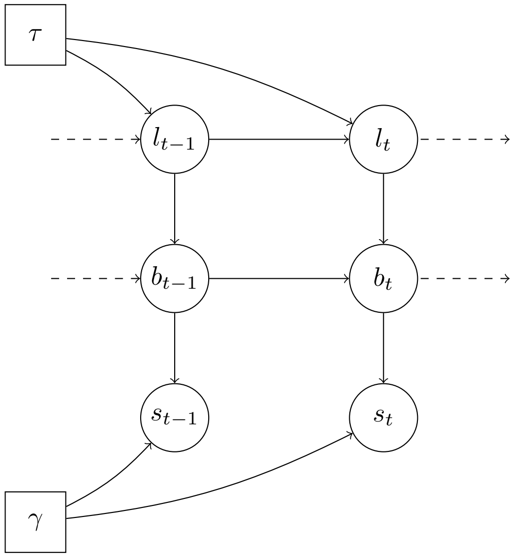

with $\tau = 60$ the scale of the exponential distribution prior over block lengths. Block length has two additional parameters not shown here, the minimum block size (20 trials) and the maximum block size (100 trials). $\gamma = 0.8$ defines the probability that the stimulus appears on a side given the block.

**Analytical equations**

Dynamics over the current length of the current blocks:

$$l_t | l_{t-1} = \begin{cases} 1 & \text{with probability } H_\tau(l_{t-1}) \\ l_{t-1} + 1 & \text{with probability } 1 - H_\tau(l_{t-1}) \end{cases}$$

with $l_1 = 1$, $H_\tau(n) = h(n)/\sum_{100 \geq n' \geq n}[h(n')]$ and $h(n) = \exp(-n/\tau) \cdot [20 \leq n \leq 100]$

Dynamics over blocks:

$$b_t | b_{t-1}, l_t = \begin{cases} b_{t-1} & \text{if } l_t > 1 \\ -b_{t-1} & \text{if } l_t = 1 \text{ and } b_{t-1} \neq 0 \\ U(\{-1, 1\}) & \text{if } l_t = 1 \text{ and } b_{t-1} = 0 \end{cases}$$

With $b_1 = 0$.

Dynamics over the stimulus side (left or right):

$$s_t | b_t = \begin{cases} U(\{-1, 1\}) & \text{if } b_t = 0 \\ b_t & \text{with probability } \gamma \text{ if } b_t \neq 0 \\ -b_t & \text{with probability } 1 - \gamma \text{ if } b_t \neq 0 \end{cases}$$

In other words: $p(s_t|b_t) = (1/2)^{b_t=0} \cdot \gamma^{(b_t \neq 0)\cdot(s_t = b_t)} \cdot (1-\gamma)^{(b_t \neq 0)\cdot(s_t \neq b_t)}$

Importantly, this generative model ignores the structure of the first 90 unbiased trials. It assumes that the first block is unbiased but doesn't force the 90 first trials to be unbiased - the unbiased first block is assumed to follow the same statistics as any other block. Assuming that the first 90 trials were unbiased worsened the explanation of mouse behaviour (Fig. 1).

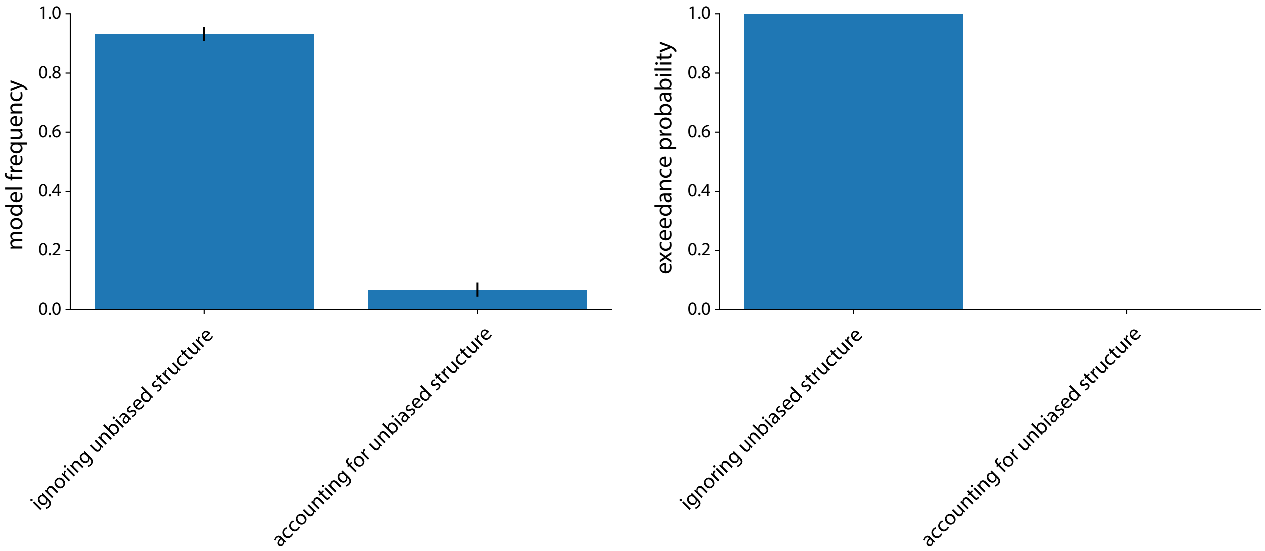

Figure 1: Bayesian Model Selection (BMS) comparing two Bayesian models: one that assumes the first block to be an unbiased block but with the same statistics as the biased blocks (*ignoring unbiased structure*), and one that assumes the first block to be unbiased and of length 90 (*accounting for unbiased structure*). Left is the model frequency and right gives this exceedance probability. The BMS favors the model that ignores the structure of the first unbiased block ($p_{exceedance} > 0.999$)

.

**Inference** To perform inference, meaning to obtain the priors $\pi_t = p\left(s_t = 1 | s_{1:(t-1)}\right)$, we apply the likelihood recursion algorithm [1].

Let us define the forward variables

$$g_t(l_t, b_t) = p(l_t, b_t, s_{1:(t-1)})$$
$$h_t(l_t, b_t) = p(l_t, b_t, s_{1:t})$$

as the joint likelihoods of $l_t, b_t$ and $s_{1:(t-1)}$ or $s_{1:t}$, respectively.

The likelihood recursion (with conditional independencies, see graphical model) gives:

$$\begin{aligned}
h_t(l_t, b_t) &= \sum_{l_{t-1}, b_{t-1}} p(s_t, s_{1:(t-1)}, l_t, b_t, l_{t-1}, b_{t-1}) \\
&= \sum_{l_{t-1}, b_{t-1}} p(s_{1:(t-1)}, l_{t-1}, b_{t-1}|) \cdot p(l_t|l_{t-1}) \cdot p(b_t|b_{t-1}, l_t) \cdot p(s_t|b_t) \\
&= p(s_t|b_t) \cdot \sum_{l_{t-1}, b_{t-1}} h_{t-1}(l_{t-1}, b_{t-1}) \cdot p(l_t|l_{t-1}) \cdot p(b_t|b_{t-1}, l_t) \\
&= p(s_t|b_t) \cdot g_t(l_t, b_t)
\end{aligned}$$

with,

$$g_t(l_t, b_t) = \sum_{l_{t-1}, b_{t-1}} h_{t-1}(l_{t-1}, b_{t-1}) \cdot p(l_t|l_{t-1}) \cdot p(b_t|b_{t-1}, l_t)$$

Obtaining the *prior* $\pi_t = p\left(s_t = 1|s_{1:(t-1)}\right)$ is now straightforward:

$$\pi_t = 0.5 \cdot p\left(b_t = 0|s_{1:(t-1)}\right) + \gamma \cdot p\left(b_t = 1|s_{1:(t-1)}\right) + (1-\gamma) \cdot p\left(b_t = -1|s_{1:(t-1)}\right)$$

with

$$p\left(b_t = k|s_{1:(t-1)}\right) = \frac{\sum_{l_t} g_t\left(l_t, b_t = k\right)}{\sum_{l_t, b_t} g_t\left(l_t, b_t\right)}$$

**Algorithm** Let T be the total number of trials, and with the notation of the previous paragraph.

**Algorithm 1:** Inference of the generative model

**Result:** priors $\pi_t = p(s_t = 1|s_{1:(t-1)})$ at every trials $t$
initialize $g_1(l_1, b_1) = p(l_1, b_1)$ for all $(l_1, b_1)$
**for** $t = 1$*;* $t \leq T$*;* $t{+}{+}$ **do**
  **for** *all* $(l_t, b_t)$ **do**
    **if** $t \geq 2$ **then**
      $g_t(l_t, b_t) = \sum_{l_{t-1}, b_{t-1}} h_{t-1}(l_{t-1}, b_{t-1}) \cdot p(l_t|l_{t-1}) \cdot p(b_t|b_{t-1}, l_t)$
    **end**
    $h_t(l_t, b_t) = p(s_t|b_t) \cdot g_t(l_t, b_t)$
  **end**
  $\pi_t = 0.5 \cdot p\left(b_t = 0|s_{1:(t-1)}\right) + \gamma \cdot p\left(b_t = 1|s_{1:(t-1)}\right) + (1-\gamma) \cdot p\left(b_t = -1|s_{1:(t-1)}\right)$
  with, $p\left(b_t = k|s_{1:(t-1)}\right) = \frac{\sum_{l_t} g_t(l_t, b_t = k)}{\sum_{l_t, b_t} g_t(l_t, b_t)}$
**end**

**Establishing the equivalence between the HMM formulation and the semi-hidden Markov formulation of the task** We verify here that the block length generated with the previously described Markovian process follows the correct truncated exponential distribution.
Let us define the random variable $X_m$ as the length of block number $m$. We can write $X_m$ as the random

variable $(l_t, l_{t+1} = 1)$, the length of the block that ends at trial $t$.
Do we have $P(X_m = k) = \left(\frac{e^{-k/\tau}}{\sum_{n=20}^{100} e^{-n/\tau}}\right)$, for all $k \in [20, 100]$ ?

$$
\begin{aligned}
P(X_m = k) &= P(l_t = k,\ l_{t+1} = 1) \\
&= \prod_{i=20}^{k-1} (1 - H_\tau(i)) \cdot H_\tau(k) \\
&= \left(\frac{e^{-k/\tau}}{\sum_{n=k}^{100} e^{-n/\tau}}\right) \cdot \prod_{i=20}^{k-1} \left(1 - \frac{e^{-i/\tau}}{\sum_{n=i}^{100} e^{-n/\tau}}\right) \propto \left(\frac{e^{-k/\tau}}{\sum_{n=k}^{100} e^{-n/\tau}}\right) \cdot \prod_{i=20}^{k-1} \left(\frac{\sum_{n=i+1}^{100} e^{-n/\tau}}{\sum_{n=i}^{100} e^{-n/\tau}}\right) \\
&= \left(\frac{e^{-k/\tau}}{\sum_{n=20}^{100} e^{-n/\tau}}\right) \cdot \left(\frac{\sum_{n=21}^{100} e^{-n/\tau}}{\sum_{n=21}^{100} e^{-n/\tau}} \ \cdots \ \frac{\sum_{n=k}^{100} e^{-n/\tau}}{\sum_{n=k}^{100} e^{-n/\tau}}\right) \\
&= \left(\frac{e^{-k/\tau}}{\sum_{n=20}^{100} e^{-n/\tau}}\right)
\end{aligned}
$$

which concludes the equivalence

##### 1.1.2 Stimulus Kernel: heuristic prior based on stimulus side integration

A second prior model consists in integrating over previous stimulus sides with an exponentially decaying kernel.

$$\pi_t | \pi_{t-1}, s_{t-1}, \alpha = (1 - \alpha) \cdot \pi_{t-1} + \alpha \cdot (s_{t-1} > 0)$$

where $s_{t-1} \in \{-1, 1\}$ is the stimulus side presented on trial $t-1$ and $\alpha$ is the learning rate.

Given all stimuli observed by the mouse $s_{1:T}$, and the learning rate $\alpha$, obtaining the heuristic prior $\pi_t$ is trivial.

To take into account the possibility of positivity and confirmation biases, we considered an alternative stimulus kernel model with asymmetrical learning rates. For this alternative, we track two variables, $\widehat{\pi}_t^{left}$ and $\widehat{\pi}_t^{right}$, which are unnormalized heuristic priors associated with the left and right stimulus side. The updating rule assumes now the existence of 4 learning rates.

- $\alpha_+^c$ : the learning rate associated with the chosen side if the chosen side was rewarded
- $\alpha_+^u$ : the learning rate associated with the unchosen side if the chosen side was unrewarded
- $\alpha_-^u$ : the learning rate associated with the unchosen side if the chosen side was rewarded
- $\alpha_-^c$ : the learning rate associated with the chosen side if the chosen side was unrewarded

The updating rule for this stimulus-side integration alternative is the following:

$$
\begin{aligned}
\widehat{\pi}_t^{left} &= \widehat{\pi}_{t-1}^{left} + \alpha_t^{left} \cdot \left((s_{t-1} < 0) - \widehat{\pi}_{t-1}^{left}\right) \\
\widehat{\pi}_t^{right} &= \widehat{\pi}_{t-1}^{right} + \alpha_t^{right} \cdot \left((s_{t-1} > 0) - \widehat{\pi}_{t-1}^{right}\right)
\end{aligned}
$$

with $\alpha_t^{left} = (a_{t-1} = -1)\left[\alpha_+^c \cdot (s_{t-1} = a_{t-1}) + \alpha_-^c \cdot (s_{t-1} \neq a_{t-1})\right] + (a_{t-1} = 1)\left[\alpha_+^u \cdot (s_{t-1} = a_{t-1}) + \alpha_-^u \cdot (s_{t-1} \neq a_{t-1})\right]$ and equivalently for $\alpha_t^{right}$.

The prior $\pi_t$ is then set according to:

$$\pi_t = \frac{\widehat{\pi}_t^{right}}{\widehat{\pi}_t^{left} + \widehat{\pi}_t^{right}}$$

##### 1.1.3 Action kernel: heuristic prior based on action integration

A third prior model consists in integrating previous actions with an exponentially decaying kernel.

$$\pi_t | \pi_{t-1}, a_{t-1}, \alpha = (1 - \alpha) \cdot \pi_{t-1} + \alpha \cdot (a_{t-1} > 0)$$

with $a_{t-1} \in \{-1, 1\}$ the action performed by the mouse at time $t-1$ and $\alpha$ the learning rate.

Given all actions performed by the mouse $a_{1:T}$ and the learning rate $\alpha$, obtaining the heuristic prior $\pi_t$ is trivial.

#### 1.2 Evidence model

We present here a generative model of the mouse's internal estimate of the stimulus contrast. We will assume here that the stimulus side is inferred in the optimal way given a noisy stimulus contrast and a subjective prior.
Let $s_t$ be the stimulus side, $c_t$ be the exact stimulus contrast at trial $t$, and $m_t$ its noisy version. We consider the mouse to have a single noisy representation $m_t$ of $c_t$ which it uses to make choices.

##### 1.2.1 Graphical representation

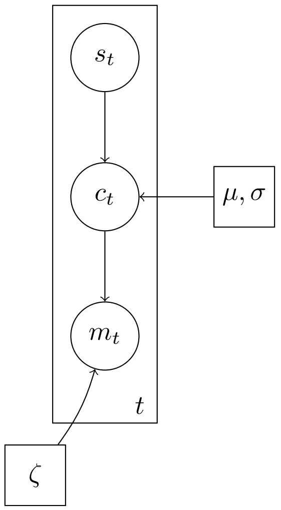

##### 1.2.2 Analytical equations

The noise structure assumed is to be doubly Gaussian:

$$p\left(c_t|s_t = 1\right) \propto f\left(c_t; \mu, \sigma\right) \cdot \mathbb{1}[c_t \geq 0]$$
$$p\left(c_t|s_t = -1\right) \propto f\left(c_t; \mu, \sigma\right) \cdot \mathbb{1}[c_t \leq 0]$$
$$m_t \mid c_t, \zeta \sim \mathcal{N}\left(c_t, \zeta\right)$$

with $f$ the Gaussian probability density function. We will assume that

$$\mu = 0$$
$$\sigma^2 = \sum_{c \in \{-1, -0.25, -0.125, -0.0625, 0, 0.625, 0.125, 0.25, 1\}} 1/9 \cdot c^2 = 0.24$$

We make this assumption of a doubly Gaussian noise structure to make the inference tractable.

##### 1.2.3 Inference

With $s_t$ known, to compute the conditional probability of the measurement $m_t$ given the side $s_t$, we need to marginalize over the contrast $c_t$. This gives;

$$p(m_t \mid s_t = 1, \zeta) = \int_{c_t} p(m_t, c_t \mid s_t = 1, \zeta) dc_t = \int_{c_t} p(c_t \mid s_t = 1) \cdot p(m_t \mid c_t, \zeta) dc_t$$
$$= \int_{c_t > 0} f(c_t; \mu, \sigma) \cdot f(m_t\, ; c_t, \zeta) dc_t = \int_{c_t > 0} \psi_t \cdot f(c_t\, ; \mu_t^*, \sigma_t^*) dc = \psi_t \cdot \Phi\left(\frac{\mu_t^*}{\sigma^*}\right)$$

with $f$ and $\Phi$ the probability density function and cumulative density function of a Gaussian distribution and:

$$\begin{aligned}
\psi_t &= f\left(\mu;\ m_t,\ \sigma^2 + \zeta^2\right) \\
(\sigma^*)^2 &= \frac{1}{\frac{1}{\sigma^2} + \frac{1}{\zeta^2}} \\
\mu_t^* &= (\sigma^*)^2 \cdot \left(\frac{\mu}{\sigma^2} + \frac{m_t}{\zeta^2}\right) = \frac{\sigma^{-2} \cdot \mu + \zeta^{-2} \cdot m_t}{\sigma^{-2} + \zeta^{-2}} = \frac{\zeta^{-2} \cdot m_t}{\sigma^{-2} + \zeta^{-2}}
\end{aligned}$$

#### 1.3 Decision policy of the mouse

##### 1.3.1 Posterior over $s_t$

We assume that the animal computes the posterior $p\left(s_t|m_t, s_{1:(t-1)}\right)$ following Bayes rule, i.e., by combining likelihood and subjective prior.

$$p\left(s_t = 1|m_t, s_{1:(t-1)}\right) \propto \psi_t \cdot \pi_t \cdot \Phi\left(\frac{\mu_t^*}{\sigma^*}\right)$$
$$p\left(s_t = -1|m_t, s_{1:(t-1)}\right) \propto \psi_t \cdot (1-\pi_t) \cdot \left(1 - \Phi\left(\frac{\mu_t^*}{\sigma^*}\right)\right)$$

##### 1.3.2 Optimal Policy with lapse rates

If the animal would seek to maximize the probability of reward, then it should pick the side $s_t$ with a higher posterior probability given its internal measurement $m_t$. This corresponds to the decision rule:

$$\psi_t \cdot \pi_t \cdot \Phi\left(\frac{\mu_t^*}{\sigma^*}\right) < \psi_t \cdot (1-\pi_t) \cdot \left(1 - \Phi\left(\frac{\mu_t^*}{\sigma^*}\right)\right) \implies \text{"choose left"};\ a_t = -1$$
$$\psi_t \cdot \pi_t \cdot \Phi\left(\frac{\mu_t^*}{\sigma^*}\right) > \psi_t \cdot (1-\pi_t) \cdot \left(1 - \Phi\left(\frac{\mu_t^*}{\sigma^*}\right)\right) \implies \text{"choose right"};\ a_t = 1$$
$$\psi_t \cdot \pi_t \cdot \Phi\left(\frac{\mu_t^*}{\sigma^*}\right) = \psi_t \cdot (1-\pi_t) \cdot \left(1 - \Phi\left(\frac{\mu_t^*}{\sigma^*}\right)\right) \implies \text{"choose randomly"}$$

After simplification

$$\mu_t^* < \sigma^* \cdot \Phi^{-1}(1-\pi_t) \implies \text{"choose left"};\ a_t = -1$$
$$\mu_t^* > \sigma^* \cdot \Phi^{-1}(1-\pi_t) \implies \text{"choose right"};\ a_t = 1$$
$$\mu_t^* = \sigma^* \cdot \Phi^{-1}(1-\pi_t) \implies \text{"choose randomly"}$$

where $\Phi^{-1}(.)$ denotes the inverse cumulative normal distribution. Thus, the optimal decision rule corresponds to a simple threshold, where the animal chooses "right" whenever its corrected internal measurement $\mu^*$ exceeds $\sigma^*\Phi^{-1}(1-\pi_t)$ and otherwise chooses "left" (with a random choice in the equality case). When the prior is uniform, $\pi_t = 0.5$, then this simplifies to choosing according to the sign of $m_t$.

When adding a lapse rate $\epsilon_t$, this decision rule can be expressed in full as

$$a_t = \text{sign}\left(\mu_t^* - \sigma^*\Phi^{-1}(1-\pi_t)\right) \quad \text{with probability } 1-\epsilon_t$$
$$\sim \mathcal{U}(\{-1; 1\}) \quad \text{with probability } \epsilon_t$$

with, $\epsilon_t = \epsilon^+ \cdot \mathbb{1}[c_t > 0] + \epsilon^- \cdot \mathbb{1}[c_t < 0] + (\epsilon^+ + \epsilon^-) \cdot 0.5 \cdot \mathbb{1}[c_t = 0]$, $\sigma^2 = 0.24$ and

$$(\sigma^*)^2 = \frac{1}{\frac{1}{\sigma^2} + \frac{1}{\zeta^2}} \qquad \mu_t^* = \frac{\zeta^{-2} \cdot m_t}{\sigma^{-2} + \zeta^{-2}}$$

To take into account the equality case, we assume that sign(0) = (-1 with probability 0.5 else 1)
We assume here contrast-side dependent lapse rates, meaning that the lapse rates will depend on whether the contrast was on the left or right side. Assuming a unique lapse rate worsened behavioural explanation 2. These contrast-side dependent lapse rates would typically occur if mice pay more attention to one side than to the other.

Additionally, two heuristics were incorporated into the decision policy for the Bayes optimal and stimulus kernel models to account for potential autocorrelations in actions. The first is a repetition bias was implemented as follows:

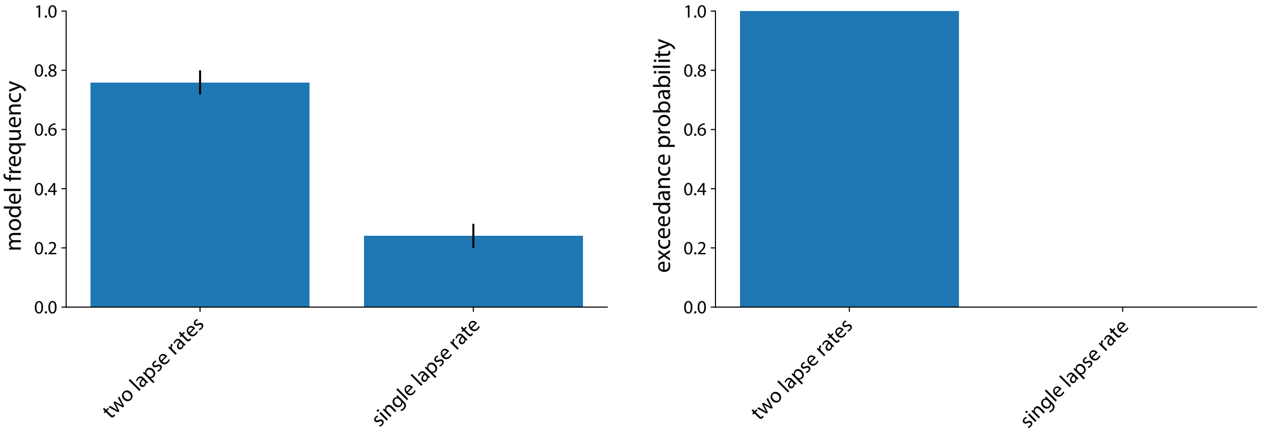

Figure 2: Bayesian Model Selection (BMS) comparing two different decision policies: one with two lapse rates (one for each stimulus side) or one with a single lapse rate. Left is the model frequency and right gives this exceedance probability. The BMS favors the model with two lapse rates ($p_{exceedance} > 0.999$). The Bayes optimal model is assumed for prior computation in both models

.

$$
\begin{aligned}
a_t &= \text{sign}\left(\mu_t^* - \sigma^* \Phi^{-1}\left(1 - \pi_t\right)\right) && \text{with probability } 1 - \epsilon_t - \eta \\
&= a_{t-1} && \text{with probability } \eta \\
&\sim \mathcal{U}(\{-1; 1\}) && \text{with probability } \epsilon_t
\end{aligned}
$$

With $\eta$ the repetition bias parameter.

The second is a multi-step repetition bias (also referred to as choice trace) was implemented as follows:

$$
\begin{aligned}
a_t &= \text{sign}\left(\mu_t^* - \sigma^* \Phi^{-1}\left(1 - \pi_t\right)\right) && \text{with probability } 1 - \epsilon_t - \eta \\
&= F_\lambda(a_{1:(t-1)}) && \text{with probability } \eta \\
&\sim \mathcal{U}(\{-1; 1\}) && \text{with probability } \epsilon_t
\end{aligned}
$$

In this case, $\eta$ denotes the multi-step repetition bias parameter and $F_\lambda$ is a function that applies an exponentially decaying average over previous actions, with $\lambda$ controlling the rate of temporal decay. Again, to take into account the equality case, we assume that sign(0) = (-1 with probability 0.5 else 1)

#### 1.4 Full likelihood

The Markovian process $\{\pi_t,\ t \in [1, T]\}$ corresponds to the priors defined in subsection 1.1. $\theta$ are the parameters of the prior model. $\theta = \emptyset$ if the Bayesian model is considered and $\theta = \{\alpha\}$ for the heuristic models (and $\theta = \left\{\alpha_+^c, \alpha_+^u, \alpha_-^c, \alpha_-^u\right\}$ for the stimulus kernel with choice- and outcome-dependent learning rates).

Let us now write the likelihood of the mouse's action $a_t$ marginalizing out the noisy contrast measurements $m_t$. We write down the equations in the case of a decision policy with lapse rates but the calculations are very similar when additionally considering a repetition bias.

$$p(a_t = 1|c_t, \pi_t, \zeta, \theta, \epsilon_+, \epsilon_-) = \int_{m_t} p(a_t = 1, m_t|c_t, \pi_t) dm_t = \int_{m_t} p(m_t|c_t, \pi_t) \cdot p(a_t = 1|m_t, \pi_t) dm_t$$
$$= \int_{m_t} f\left(m_t; c_t, \zeta\right) \cdot \mathbb{1}\left[\mu_t^* > \sigma^* \Phi^{-1}\left(1 - \pi_t\right)\right] dm_t \cdot (1 - \epsilon_t) + \epsilon_t/2$$

with $\mu_t^* = \frac{\zeta^{-2}}{\sigma^{-2}+\zeta^{-2}} \cdot m_t$, $\epsilon_t = \epsilon^+ \cdot \mathbb{1}[c_t > 0] + \epsilon^- \cdot \mathbb{1}[c_t > 0] + (\epsilon^+ + \epsilon^-) \cdot 0.5 \cdot \mathbb{1}[c_t = 0]$ and f the Gaussian pdf.

For the first term:

$$\int_{m_t} f\left(m_t; c_t, \zeta\right) \cdot \mathbb{1}\left[m_t > \left(\frac{\zeta^2}{\sigma^2} + 1\right) \sigma^* \Phi^{-1}\left(1 - \pi_t\right)\right] dm_t = \int_{\left(\frac{\zeta^2}{\sigma^2}+1\right)\sigma^* \Phi^{-1}(1-\pi_t)}^{+\infty} f\left(m_t; c_t, \zeta\right) dm_t$$
$$= \int_0^{+\infty} f\left(m_t; c_t - \left(\frac{\zeta^2}{\sigma^2} + 1\right) \sigma^* \Phi^{-1}\left(1 - \pi_t\right), \zeta\right) dm_t$$
$$= \Phi\left(\frac{c_t}{\zeta} - \left(\frac{\zeta^2}{\sigma^2} + 1\right) \frac{\sigma^*}{\zeta} \Phi^{-1}\left(1 - \pi_t\right)\right)$$

Thus:

$$p(a_t = 1|c_t, \pi_t, \zeta, \theta, \epsilon_+, \epsilon_-) = \Phi\left(\frac{c_t}{\zeta} - \left(\frac{\zeta^2}{\sigma^2} + 1\right) \frac{\sigma^*}{\zeta} \Phi^{-1}\left(1 - \pi_t\right)\right) \cdot (1 - \epsilon_t) + \frac{\epsilon_t}{2}$$

Given all actions of a mouse on a session $a_{1:T}$ ($a_t \in \{-1, 1\}$) and the corresponding contrasts $c_{1:T}$ ($c_t \in \{-1, 1\}$), we obtain the loglikelihood:

$$\log\ p\left(a_{1:T}|c_{1:T}, \pi_{1:T}, \zeta, \theta, \epsilon_+, \epsilon_-\right) = \sum_{t=1}^{T} \log\ p\left(a_t|c_t, \pi_t, \zeta, \theta, \epsilon_+, \epsilon_-\right)$$
$$= \sum_{t=1}^{T} \mathbb{1}[a_t = 1] \cdot \log\ \rho_t + \mathbb{1}[a_t = -1] \log\ (1 - \rho_t)$$

with

$$\rho_t = \Phi\left(\frac{c_t}{\zeta} - \left(\frac{\zeta^2}{\sigma^2} + 1\right) \frac{\sigma^*}{\zeta} \Phi^{-1}\left(1 - \pi_t\right)\right) \cdot (1 - \epsilon_t) + \frac{\epsilon_t}{2}$$
$$\epsilon_t = \epsilon^+ \cdot \mathbb{1}[c_t > 0] + \epsilon^- \cdot \mathbb{1}[c_t > 0] + (\epsilon^+ + \epsilon^-) \cdot 0.5 \cdot \mathbb{1}[c_t = 0]$$
$$(\sigma^*)^2 = \frac{1}{\frac{1}{\sigma^2} + \frac{1}{\zeta^2}}$$

$\pi_t$ is a function of $\theta$. Inference in the generative model of mouse behaviour implies computing the posterior over the parameters $\theta$, $\epsilon_+$, $\epsilon_-$, $\zeta$.

This is done by implementing a random walk adaptive Metropolis-Hastings [2] [3]. The procedure is adaptive as it learns the covariance matrix of the random walk to obtain relevant samples and acceptance ratios (see articles for more information). Furthermore, 4 chains are run in parallel and the Gelman-Rubin factor is used to assess convergence. In terms of priors, we use uniform distributions ranging from 0 to 1 for both the learning rates and sensory noise, $\zeta$. Furthermore, we set uniform priors between 0 and 0.5 for the lapse rates and repetition biases.

NB: For the Bayes optimal model, we do not perform inference over $\{\tau, \gamma\}$, we fix them to their "correct" values.

### 2 Hierarchical model

We designed a hierarchical model that accounts for two types of variability: within individual mice and within sessions for a given mouse. This model is employed to evaluate the correlation between neural and behavioral learning rates. It establishes session-level parameters that are derived from distributions at the mouse level, which in turn are based on distributions at the population level.

#### 2.1 Model Specification

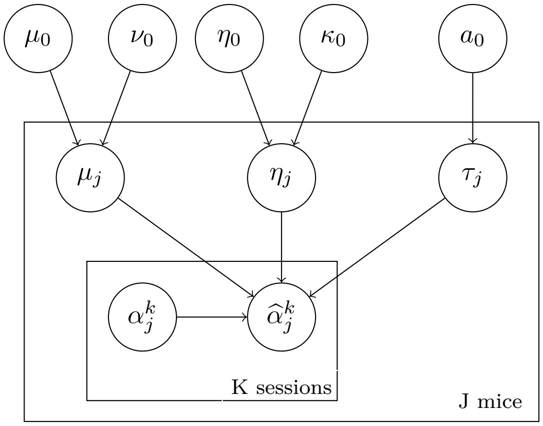

Linear Regression:

$$\widehat{\alpha}_j^k | \mu_j, \eta_j, \tau_j, \alpha_j^k \sim \mathcal{N}(\mu_j \cdot \alpha_j^k + \eta_j, \tau_j)$$

with $\widehat{\alpha}_j^k$ the learning rate (inverse decay constant) estimated from the neural activity and $\alpha_j^k$ the learning rate (inverse decay constant) estimated from behaviour. This linear regression is defined for each session $k$ and mouse $j$. $\mu_j$ is the slope, $\eta_j$ the intercept and $\tau_j$ the standard deviation of the linear regression defined at the mouse-level: these parameters are shared across all sessions for the same mouse.

Mouse-level parameters:

$$\mu_j | \mu_0, \nu_0 \sim \mathcal{N}(\mu_0, \nu_0)$$
$$\eta_j | \eta_0, \kappa_0 \sim \mathcal{N}(\eta_0, \kappa_0)$$
$$\tau_j \sim Exp(a_0)$$

The mouse-level slopes $\mu_j$ and intercepts $\eta_j$ are assumed to follow Gaussian distributions and the standard deviations $\tau_j$ are assumed to follow an exponential distribution. These priors are defined with population-level parameters. $\mu_0$ and $\nu_0$ are the mean and standard deviation of the Gaussian prior over the mouse-level slopes $\mu_j$. $\eta_0$ and $\kappa_0$ are the mean and standard deviation of the Gaussian prior over the mouse-level intercepts $\eta_j$. Lastly, $a_0$ is the parameter of the exponential prior over mouse-level standard deviations $\tau_j$.

Population-level parameters:

$$\mu_0 \sim N(0, 1)$$
$$\nu_0 \sim Exp(0.1)$$
$$\eta_0 \sim N(0, 1)$$
$$\kappa_0 \sim Exp(0.1)$$
$$a_0 \sim Exp(0.1)$$

These last equations describe the priors over the population-level parameters $\mu_0$, $\nu_0$, $\eta_0$, $\kappa_0$ and $a_0$.

#### 2.2 Inference

Inference is done with Metropolis-Hasting, running a chain for 50000 iterations and selecting the second half of samples to compute the posteriors.

The main random variables of interest are $\mu_0$ and $\{\mu_j\}_j$, the slopes at the population level and at the mouse level. These are shown in supplementary figure S15.

### References

[1] Scott, S. L. (2002). Bayesian methods for hidden Markov models: Recursive computing in the 21st century. *Journal of the American statistical Association, 97(457)*, 337-351.

[2] Andrieu, C., & Thoms, J. (2008). A tutorial on adaptive MCMC. *Statistics and computing, 18*, 343-373.

[3] Baker, M. J. (2014). Adaptive Markov chain Monte Carlo sampling and estimation in Mata. *The Stata Journal, 14(3)*, 623-661.
